# Iron-meditated fungal starvation by lupine rhizosphere-associated and extremotolerant *Streptomyces* sp. S29 desferrioxamine production

**DOI:** 10.1101/2020.06.29.145664

**Authors:** Scott A. Jarmusch, Diego Lagos-Susaeta, Emtinan Diab, Oriana Salazar, Juan A. Asenjo, Rainer Ebel, Marcel Jaspars

## Abstract

Siderophores are iron-chelating compounds that aid iron uptake, one of the key strategies for microorganisms to carve out ecological niches in microbially diverse environments. Desferrioxamines are the principal siderophores produced by *Streptomyces* spp. Their biosynthesis has been well studied and as a consequence, the chemical potential of the pathway continues to expand. With all of this in mind, our study aimed to explore extremotolerant and Lupine rhizosphere-derived *Streptomyces* sp. S29 for its potential antifungal capabilities. Cocultivation of isolate S29 was carried out with *Aspergillus niger* and *Botrytis cinerea*, both costly fungal phytopathogens in the wine industry, to simulate their interaction within the rhizosphere. The results indicate that not only is *Streptomyces* sp. S29 extraordinary at producing hydroxamate siderophores but uses siderophore production as a means to ‘starve’ the fungi of iron. High resolution LC-MS/MS followed by GNPS molecular networking was used to observe the datasets for desferrioxamines and guided structure elucidation of new desferrioxamine analogues. Comparing the new chemistry, using tools like molecular networking and MS2LDA, with the known biosynthesis, we show that the chemical potential of the desferrioxamine pathway has further room for exploration.

## INTRODUCTION

The application of *Streptomyces* secondary metabolites goes beyond drug discovery, as they are one of the most important bacteria for crop protection. Their biotechnological application as rhizosphere or endophytic bacteria is proven to confer protection against soilborne pathogens like fungi, nematodes, and other bacteria.^1,2^ They serve as an important factor in ‘disease suppressive soils’, with several mechanisms of action including synthesis of plant growth regulators^3^, antibiotic production^2^, secretion of volatile compounds^4^, and siderophore production.^5^ The last of these is not only a way for bacteria to uptake vital metals from the surrounding soil but also to sequester metals in order to gain a competitive edge against other soilborne organisms.

Rhizosphere-associated microbes are one of the many strategies currently being explored as biocontrol agents. Bacterial communities in the rhizosphere contribute to the extremely complex flow of nutrients as well as the export of antibiotics and other secondary metabolites into the soil.^6,7^ Model organisms from genera like *Bacillus* and *Pseudomonas* have traditionally been promising sources due to their association to the soil and their production of secondary metabolites like lipopeptides but also plant growth promotors.^8,9^ Actinobactera, which are also predominantly soil-associated microbes, have also been explored as biocontrol agents due to their prolific antimicrobial potential.^10^ This combined with the fact that extremotolerant microbes have adapted to the harsh conditions and the potential of these adaptations to be translatable to biotechnologies, extremotolerant microbes look to be promising sources as biocontrol agents.^11^ In Chile, the wine industry is the main agricultural export, estimated at ~$2 billion USD in 2017^12^, thus driving a large research interest around protection of wine-producing grapes against various phytopathogens.

Siderophores are iron-chelating compounds that aid metal uptake in iron deficient environments.^13^ These secondary metabolites are produced by all microbes and four types of siderophore-mediated social interactions have been observed: uptake among clonal cells (sharing), cheating or piracy (which can only happen amongst microbes with the same uptake receptor), competition via locking away and metabolite competition (whichever microbe makes the most cost efficient and effective chelator).^14^ Desferrioxamines (DFO) are the most widely observed group of siderophores in *Streptomyces* spp., used principally for Fe(III) scavenging^13,15^, as well as some other heavy metals.^16–18^ Since the vast majority of organisms require Fe(III) in order to maintain proper cell wall function^13^, as one scavenges for itself, it simultaneously starves other competing organisms, rendering them potentially unviable.

The biosynthetic machinery of desferrioxamines has been well studied^19–21^ and as a consequence, the chemical potential continues to expand. The substrate specificity of DesC-like proteins, similar to acyl CoA-dependent transferases, seems to be the key factor in producing diverse desferrioxamine scaffolds.^19,21^ It has been shown that DesC can accommodate for acetyl, succinyl, and myristol-CoA, with the latter providing insight into the production of acyl desferrioxamines.^19^ Additionally, aryl desferrioxamines, first isolated from *Micrococcus luteus* KLE1011^22^ and later from *Streptomyces* sp. MA37^23^, showed further substrate flexibility, where the latter communication described a new biosynthetic gene cluster (BGC) which contained a DesC homolog (LgoC). This enzyme promiscuity fits ideally with the Screening Hypothesis first proposed by Firn and Jones; secondary metabolism stems from metabolic pathways that ‘maximize diversity and minimize cost’.^24^

Due to the modular nature of desferrioxamines and the presence of peptide bonds throughout their structure, mass spectrometry-based studies have been utilized extensively to not only study their biological impact but also for elucidation of new structures. Previous studies have established standard fragmentation patterns of these siderophores, even as far as elucidating new structures solely based on mass spectrometry.^25–31^ This standardized fragmentation makes the discovery of siderophores much easier thanks to metabolomics-based tools like molecular networking, like Global Natural Products Social Molecular Networking (GNPS). It allows for correlations to be drawn within your dataset and against the GNPS library that relates similar MS/MS spectra to one another, forming clusters or families of similar features.^32^

With all of this in mind, our study originally aimed to explore extremotolerant, rhizosphere-derived *Streptomyces* sp. S29, a novel strain, for antifungal secondary metabolites. Cocultivation of S29 with *Aspergillus niger* and *Botrytis cinerea*, both costly fungal phytopathogens in the wine industry, were carried out in a fashion to allow S29 to dominate the culture, thus eliciting a chemical response from the addition of each fungus. Bioassay-guided fractionation lead to no lead compounds as potential antifungal agents yet post-fermentation streaks of inoculum showed no fungal growth. High resolution LC-MS/MS followed by GNPS molecular networking was used to analyse the data sets. The results indicated that not only is *Streptomyces* sp. S29 extraordinary at producing hydroxamate siderophores, totalling 24 new analogues and 8 previously identified ones confirmed through MS/MS structure elucidation, but uses siderophore production as a means to ‘starve’ the fungi of iron. When grown in coculture with the fungi, 22 additional new analogues and 17 additional previously described were identified, clearly showing a chemical response to the fungi in culture environment. Using MS2LDA to further explore desferrioxamine chemical space, we observed that it is potentially 2.5 times larger than previously anticipated. These results point to the potential of *Streptomyces* sp. S29 as a biocontrol agent to prevent fungal infection and the potential for new desferrioxamine chemistry.

## RESULTS AND DISCUSSION

### *Streptomyces* sp. S29 cocultivation and LC-MS of desferrioxamine-rich fractions

Preliminary fungal bioassay screening of the lupine rhizosphere *Streptomyces* culture collection showed a novel strain, *Streptomyces* sp. S29, inhibited the growth of *Botrytis cinerea* when grown in coculture on solid medium (Figure 1). In an attempt to induce production of antifungal metabolites, the bacteria and fungi were cocultivated in liquid culture in a fashion to allow for *Streptomyces* sp. to be the dominant microbe. Bioassay guided fractionation of the *sec-*butanol fraction lead to no antifungal leads even though inhibition was observed against *B. cinerea* only (Figure S6). Upon LC-MS and ^1^H NMR analysis of the extracts, a profusion of DFOs was observed in all *sec*-butanol fractions, leading to the hypothesis that this *Streptomyces* sp. uses a different mechanism than producing antimycotics to combat the fungal cocultivate, as seen previously in human bacterial infections.^33^ DFOs historically have never exhibited antimicrobial activity and therefore we confidently presume they are not responsible for the inhibition seen in Figure S6.^34^ Time-dependent growth inhibition of *Penicillium* was observed when exposed to varying concentrations of coproporphyrin, a siderophore produced by *Glutamicibacter arilaitensis*^35^, but this activity has never been shown for desferrioxamines. *Streptomyces* sp. S29 was also grown in monoculture on iron deficient medium to induce production of DFOs in an attempt to compare siderophore production to the coculture experiments. *Aspergillus niger* and *Botrytis cinerea* were chosen as the fungal phytopathogens due to their established pathology of wine producing grapes ^36,37^ and the importance of wine as Chile’s main agricultural export. Additionally, these fungi would be encountered in *Streptomyces* sp. S29’s ecological niche, making this an important evaluation of microbes it would encounter in the soil environment.

**Figure 1.**
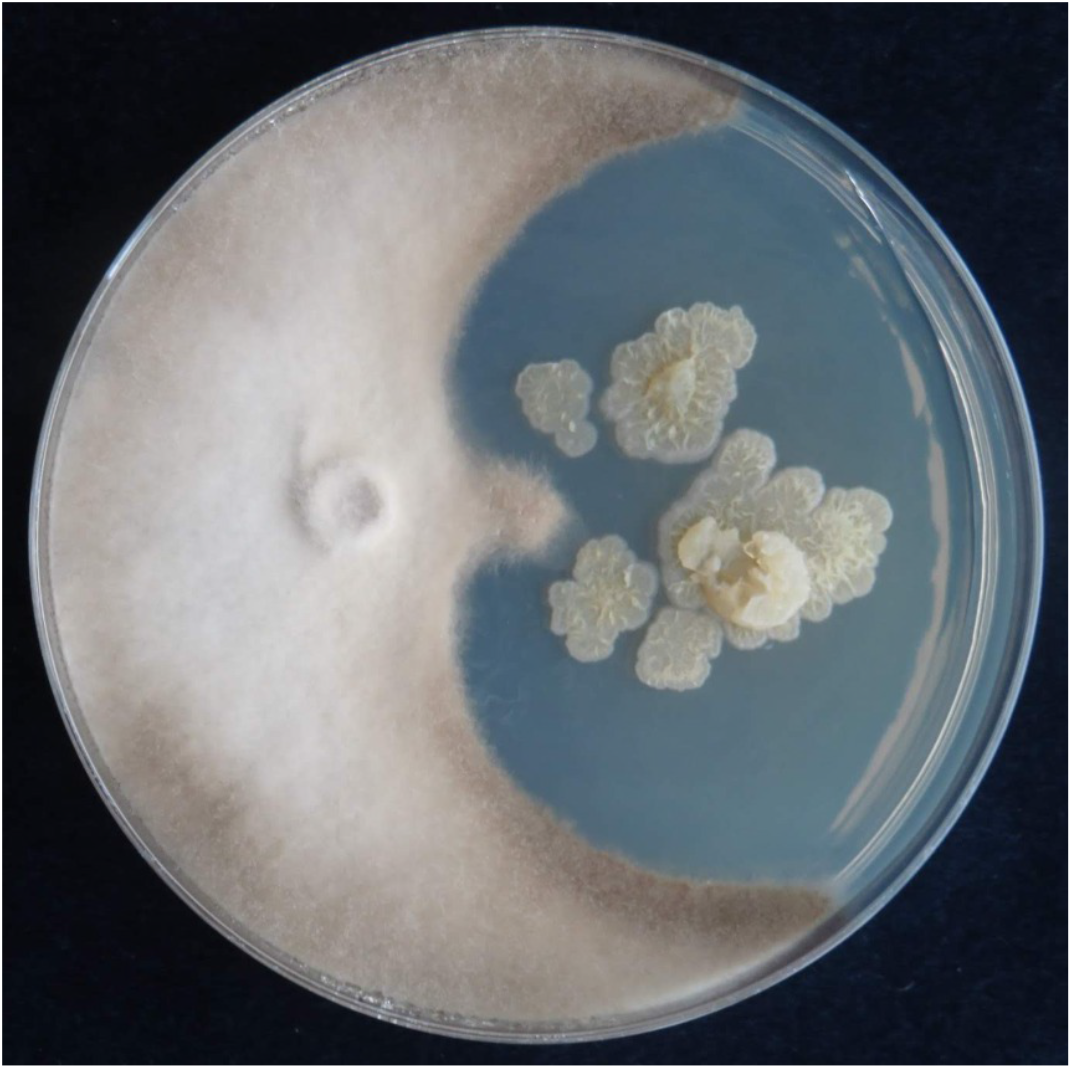
Preliminary antifungal screening on *Streptomyces* isolate S29 (right) grown with *Botrytis cinerea* (left). Image courtesy of Diego Lagos-Susaeta.

Extracts and fractions were evaluated for the presence of two DFOs (B and D1) during the fractionation steps, in order to trace the fractions that contained the highest abundance of siderophores. Presence of DFOs was further evaluated at the final stages of separation based on the molecular families generated from molecular networking, utilizing the known DFOs in the GNPS library as anchor points, which further aided structure elucidation. Based on the LC-MS results, the butanol partitions contained the largest abundance of DFOs, and therefore this data was used to generate a molecular network using GNPS.

### Molecular networking and MS/MS based structure elucidation

The molecular network generated contained 1169 molecular features after removal of features originating from media and instrument blanks. Each subnetwork was approached using GNPS library hits as ‘anchors’ (hexagons in each subnetwork) and then collecting the masses of linked nodes and predicting molecular formula from the raw data. Using these ‘anchors’ not only allows one to relate the nodes directly connected to the anchor, but also gives broader sense of the molecular nature of a cluster. The hydroxamate siderophores were primarily distributed amongst three principle molecular families (Figures 2–4). Subnetwork 1 (Figure 2) contained a number of new acyl DFO D-like analogues that are first described in this study as well as new analogues of acyl DFO B-like derivatives. Subnetwork 2 (Figure 3) is mostly composed of unknown derivatives although a suite of new macrocyclised DFOs and tenacibactin derivatives, first described by Jang et al. ^38^, were observed. The last principle subnetwork contained various Fe(III) and Al(III) complexed FOs (Figure 6). Within this network a new analogue was identified, uC_12_ acyl FO-B, while its non-chelated form was not observed in the data (Figure S32).

**Figure 2.**
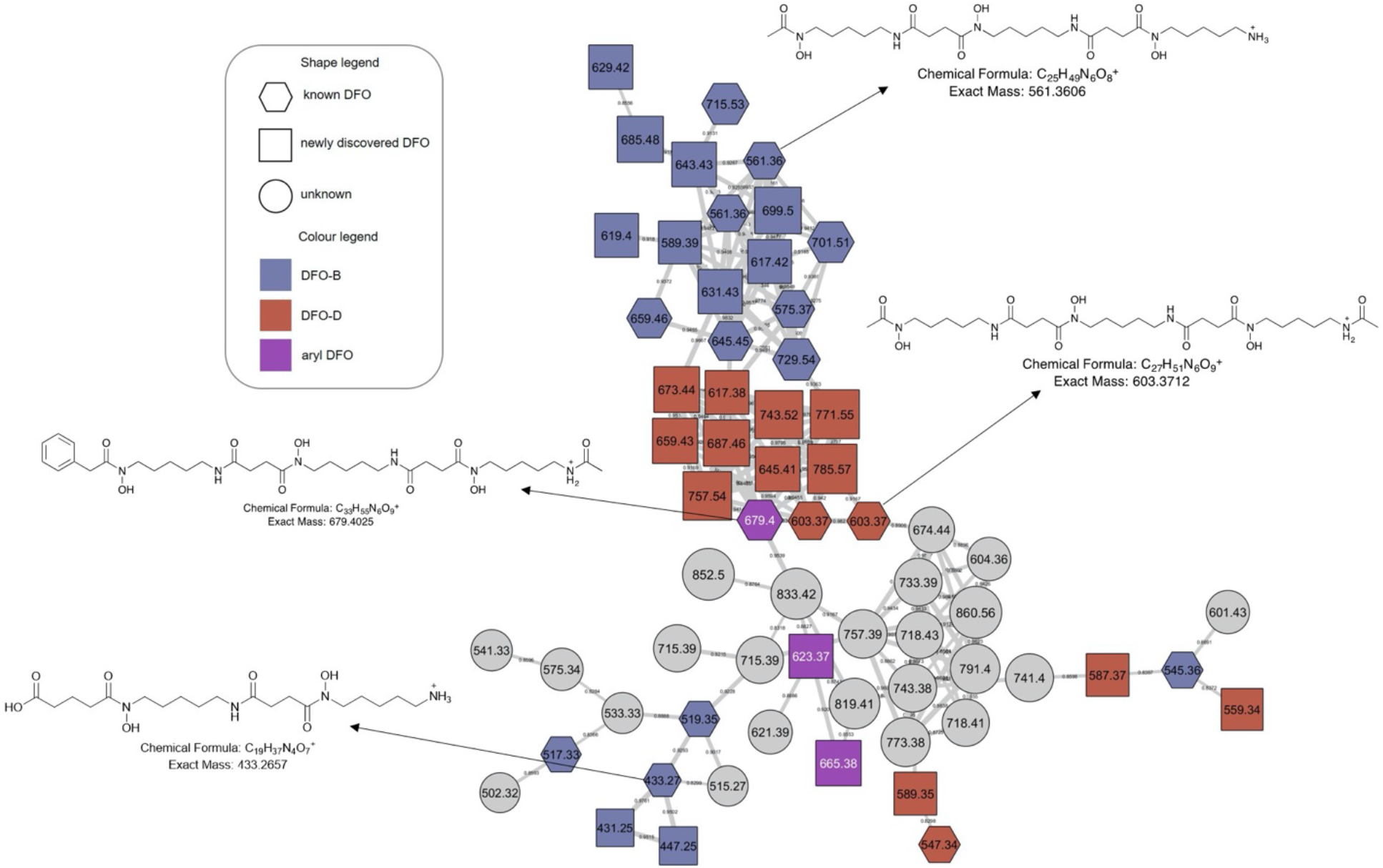
Desferrioxamine subnetwork 1. Distributed amongst the molecular family are DFO-Bs (blue), DFO-Ds (red), and aryl DFOs (purple). Grey nodes represent potential DFOs that were not able to be annotated using MS/MS data only.

**Figure 3.**
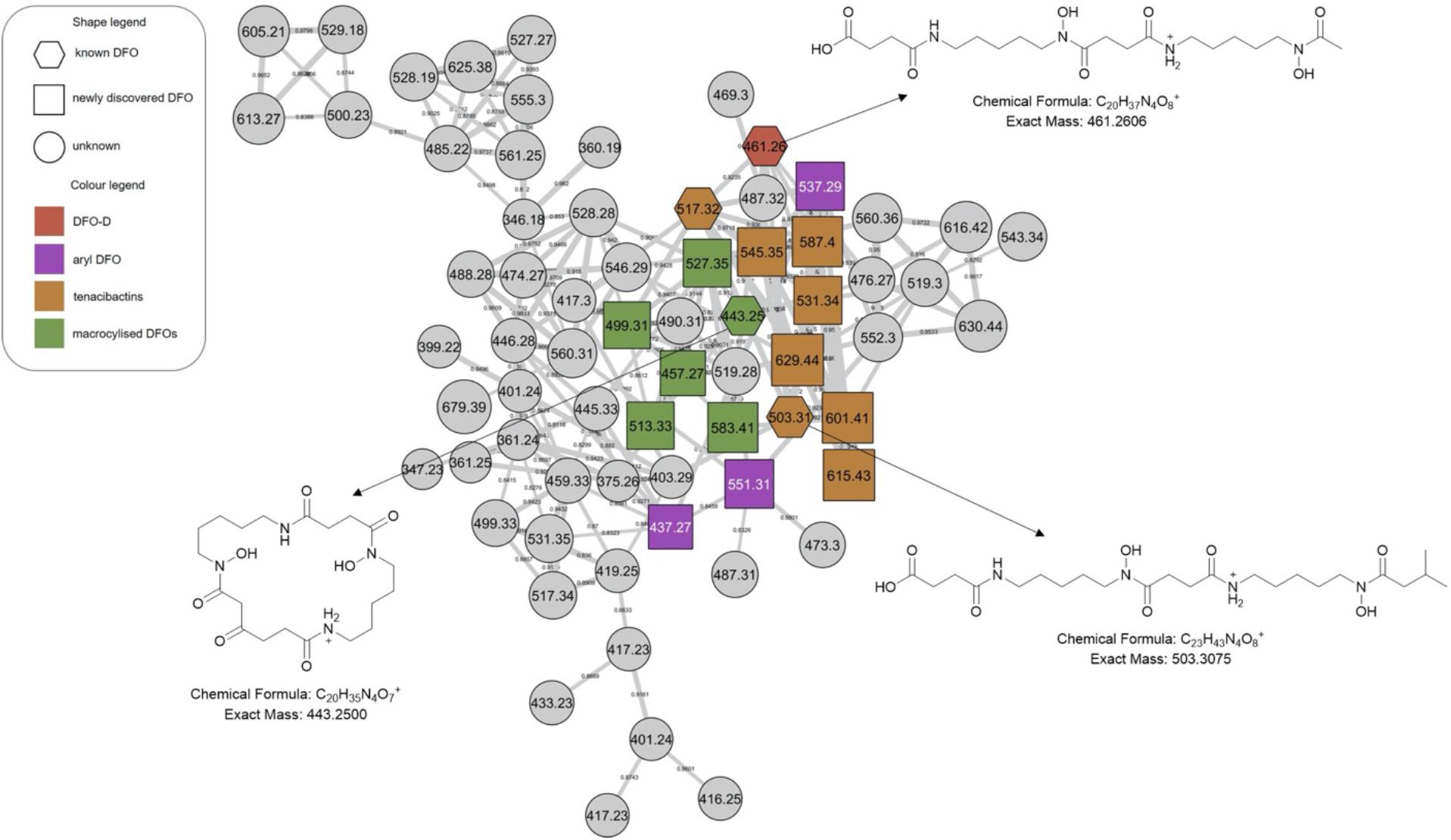
Desferrioxamine subnetwork 2. Within the molecular family are macrocyclised DFOs (green) as well as a number of new tenacibactin derivatives (orange). Grey nodes represent potential DFOs that were not able to be annotated using MS/MS data only.

**Figure 4.**
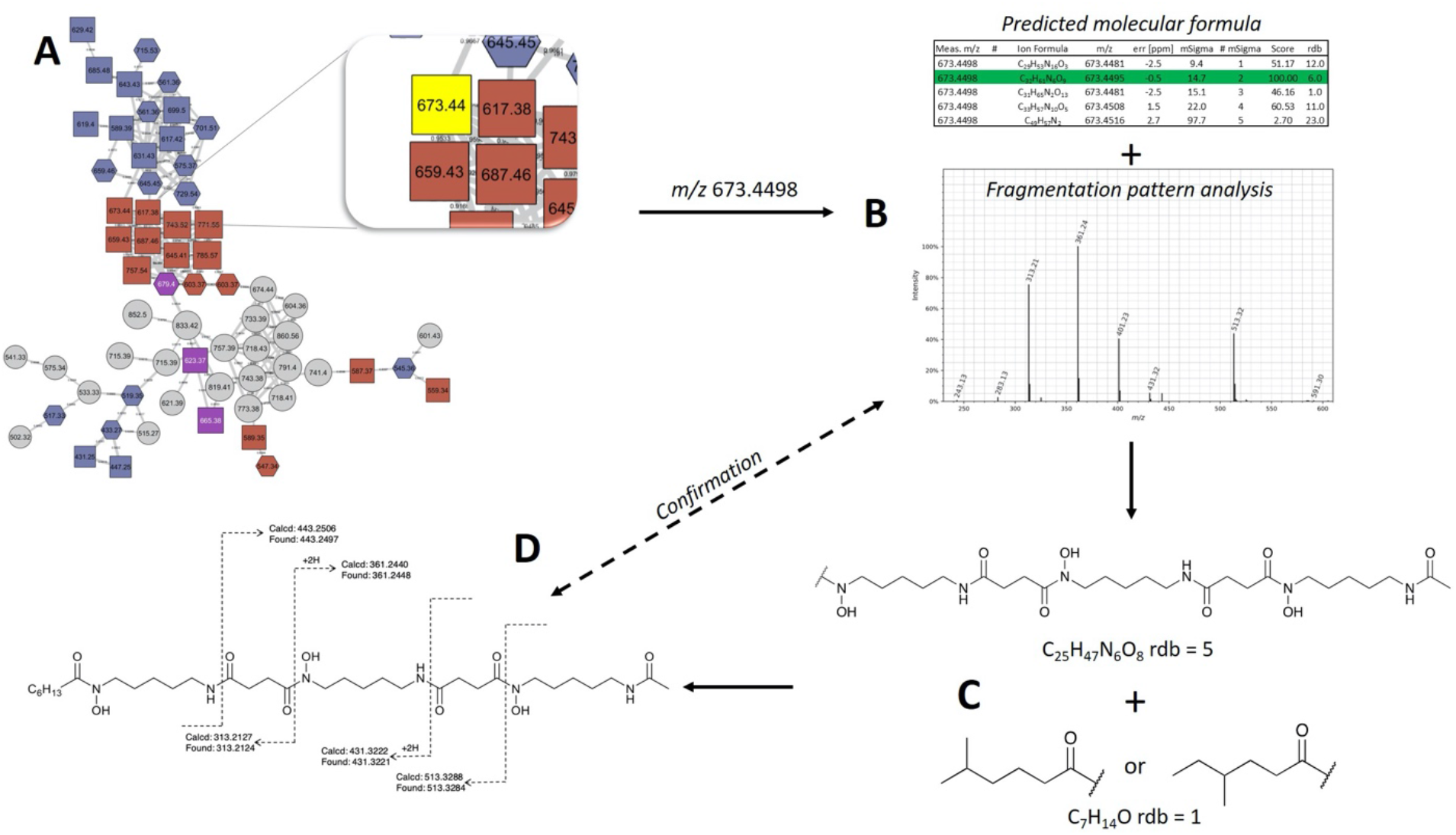
Molecular network and MS/MS based structure determination. (A) molecular networks generated using GNPS were dereplicated and potentially unknown ions were found in the Raw data. (B) Predicted molecular formula were generated and the most likely candidate formula was chosen. MS/MS spectra were analysed for the initial loss of −118 or −160 to determine the N-terminal moiety of *N*-hydroxycadaverine or *N*-acetylhydroxycadaverine, respectively. (C) Based on known structure of desferrioxamines, substructures were proposed for each new metabolite and lastly, (D) putative structures were supported by MS/MS data, although further spectroscopic characterisation would be required for confirmation.

Aryl desferrioxamines related to the previously described acyl ferrioxamine 1 (also known as legonoxamine A) and 2 were also found distributed amongst varying molecular families across the molecular network and do not seem to link into their own cluster such as that seen for the DFO-Ds and DFO-Bs. For the remaining small molecular families that contain DFOs, we observed a group of sodiated adducts, large acyl DFO-Bs, as well as the macrocyclised desferrioxamine E (Figure S1). The remaining nodes that link within the desferrioxamine molecular families were unable to be identified using the annotation workflow described in Figure 4.

MS/MS based structure elucidation was carried out in a similar fashion to studies conducted previously (Figure 4).^25–28^ Early observations of desferrioxamine B and D1 fragmentation showed an initial loss of the N-terminus, corresponding to the loss of either 118.1 or 160.1. Using this as a foundation, structures were either started with a *N*-hydroxycadaverine or a *N*-hydroxy-*N*-acetylcadaverine, respectively. The remainder of structure elucidation was completed stepwise: (1) predicting the molecular formula from the accurate mass and collection of MS/MS data, (2) filling out the structures with appropriate desferrioxamine precursors (i.e. *N*-hydroxycadaverine, putrescine, succinate, acetate, etc.), and (3) taking the remainder of the molecular formula and placing an appropriate acid at the C-terminus that either has precedent from previous studies or could be a potential binding partner with an acyltransferase like coenzyme A.

Overall, 41 new DFO analogues were identified in the molecular network and through manual data processing (Table 1). These range from short acyl chain DFO-Bs, acyl desferrioxamine D-like (DFO-D) analogues which have not previously been described, tenacibactins, aryl DFOs, and long chain dicarboxylate DFOs (Table 1). Additionally, 5 uncertain macrocyclic and 25 known hydroxamate siderophores were detected, including known acyl, aryl DFOs and bisucaberins, giving further credence to the new structure assignments. Their identification, observed presence in monoculture and/or coculture, and identification confidence based on Schymanski’s rules^39^ are present in Table 1. Schymanski’s rule distinguish various levels of confidence for mass spectral-based assignments, where the highest score is 1 (verifiable by MS, MS^2^, RT, and an internal standard) and the lowest score is 5 (exact mass only). The majority of the identified compounds rank at level 2 or level 3 on the Schymanski scale.

**Table 1.** Summary of siderophores identified using molecular networking and manual observations of the data. Nomenclature that are italicized were identified in this study.

|  |  |  |  |  |  |  |  |  |
| --- | --- | --- | --- | --- | --- | --- | --- | --- |
|  | Name | Measured mass<br>[M+H]<sup>+</sup> | Theoretical mass<br>[M+H]<sup>+</sup> | Δ ppm | Sum formula<br>[+H]<sup>+</sup> | ID confidence | Monoculture | Reference |
| cyclic desferrioxamines | Bisu-03 | 429.2344 | 429.2342 | -0.46 | C<sub>19</sub>H<sub>33</sub>N<sub>4</sub>O<sub>7</sub> | 2 |  | Senges et al. |
|  | Bisu-01 | 443.2495 | 443.2500 | -1.13 | C<sub>20</sub>H<sub>35</sub>N<sub>4</sub>O<sub>7</sub> | 2 |  | Senges et al. |
|  | *Uncertain structure* | 457.2660 | 457.2657 | 0.65 | C<sub>21</sub>H<sub>37</sub>N<sub>4</sub>O<sub>7</sub> | 3 | * | *This study* |
|  | *Uncertain structure* | 499.3110 | 499.3126 | -3.20 | C<sub>24</sub>H<sub>43</sub>N<sub>4</sub>O<sub>7</sub> | 3 | * | *This study* |
|  | *Uncertain structure* | 513.3310 | 513.3283 | 5.25 | C<sub>25</sub>H<sub>45</sub>N<sub>4</sub>O<sub>7</sub> | 3 | * | *This study* |
|  | *Uncertain structure* | 527.3450 | 527.3439 | 2.08 | C<sub>26</sub>H<sub>47</sub>N<sub>4</sub>O<sub>7</sub> | 3 | * | *This study* |
|  | *Uncertain structure* | 583.4050 | 583.4065 | -2.57 | C<sub>30</sub>H<sub>55</sub>N<sub>4</sub>O<sub>7</sub> | 3 |  | *This study* |
|  | desferrioxamine E | 601.3557 | 601.3556 | 0.17 | C<sub>27</sub>H<sub>49</sub>N<sub>6</sub>O<sub>9</sub> | 2 |  |  |
| aryl desferrioxamines | legonoxamine B | 337.1754 | 337.1758 | -1.19 | C<sub>17</sub>H<sub>25</sub>N<sub>2</sub>O<sub>5</sub> | 2 |  | Maglangit et al. |
|  | *legonoxamine C* | 351.1912 | 351.1914 | -0.57 | C<sub>18</sub>H<sub>37</sub>N<sub>4</sub>O<sub>5</sub> | 3 |  | *This study* |
|  | *legonoxamine D* | 437.2755 | 437.2758 | -0.69 | C<sub>22</sub>H<sub>37</sub>N<sub>4</sub>O<sub>5</sub> | 3 |  | *This study* |
|  | *legonoxamine E* | 537.2924 | 537.2919 | 0.93 | C<sub>26</sub>H<sub>41</sub>N<sub>4</sub>O<sub>8</sub> | 3 | * | *This study* |
|  | *legonoxamine F* | 551.3088 | 551.3075 | 2.36 | C<sub>27</sub>H<sub>43</sub>N<sub>4</sub>O<sub>8</sub> | 3 |  | *This study* |
|  | *legonoxamine G* | 623.3764 | 623.3763 | 0.16 | C<sub>30</sub>H<sub>51</sub>N<sub>6</sub>O<sub>8</sub> | 3 | * | *This study* |
|  | acyl-ferrioxamine 1/<br>legonoxamine A | 637.3917 | 637.3919 | -0.31 | C<sub>31</sub>H<sub>53</sub>N<sub>6</sub>O<sub>8</sub> | 2 | * | D'Onofrio et al;<br>Maglangit et al. |
|  | *legonoxamine H* | 665.3893 | 665.3869 | 3.61 | C<sub>32</sub>H<sub>53</sub>N<sub>6</sub>O<sub>9</sub> | 3 |  | *This study* |
|  | acyl-ferrioxamine 2 | 679.4025 | 679.4025 | 0.00 | C<sub>33</sub>H<sub>55</sub>N<sub>6</sub>O<sub>9</sub> | 2 | * | D'Onofrio et al. |
|  | *legonoxamine A glycoside* | 799.4456 | 799.4448 | 1.00 | C<sub>37</sub>H<sub>63</sub>N<sub>6</sub>O<sub>13</sub> | 3 | * | *This study* |
| DFO-Bs | *un-pre[5+5\*]* | 431.2500 | 431.2520 | -4.63 | C<sub>19</sub>H<sub>35</sub>N<sub>4</sub>O<sub>7</sub> | 3 |  | *This study* |
|  | *pre[5+5\*]* | 433.2662 | 433.2657 | -1.15 | C<sub>19</sub>H<sub>37</sub>N<sub>4</sub>O<sub>7</sub> | 2 |  | Rütschlin et al. |
|  | *pre[5+5\*] aldehyde* | 447.2450 | 447.2449 | 0.22 | C<sub>19</sub>H<sub>35</sub>N<sub>4</sub>O<sub>8</sub> | 3 |  | *This study* |
|  | *pre[ha+5+5\*]* | 517.3221 | 517.3232 | -2.12 | C<sub>24</sub>H<sub>45</sub>N<sub>4</sub>O<sub>8</sub> | 2 |  | Rütschlin et al. |
|  | abloxime / IC202C | 517.3347 | 517.3344 | 0.58 | C<sub>23</sub>H<sub>45</sub>N<sub>6</sub>O<sub>7</sub> | 2 | * | Iijima et al. |
|  | proferrioxamine G1t | 519.3495 | 519.3501 | -1.16 | C<sub>23</sub>H<sub>47</sub>N<sub>6</sub>O<sub>7</sub> | 2 |  | Feistner et al. |
|  | DesuA1 | 545.3658 | 545.3657 | 0.18 | C<sub>25</sub>H<sub>49</sub>N<sub>6</sub>O<sub>7</sub> | 2 | * | Senges et al. |
|  | DesA1 | 547.3449 | 547.3450 | -0.18 | C<sub>24</sub>H<sub>47</sub>N<sub>6</sub>O<sub>8</sub> | 2 | * | Senges et al. |
|  | Des A2 / IC202B | 557.2880 | 557.2881 | -0.18 | C<sub>23</sub>H<sub>43</sub>N<sub>6</sub>O<sub>8</sub>Al | 2 |  | Iijima et al. |
|  | desferrioxamine B | 561.3602 | 561.3606 | -0.71 | C<sub>25</sub>H<sub>49</sub>N<sub>6</sub>O<sub>8</sub> | 2 | * |  |
|  | desferrioxamine N /desf-05 | 575.3762 | 575.3763 | -0.17 | C<sub>26</sub>H<sub>51</sub>N<sub>6</sub>O<sub>8</sub> | 2 | * | Ejje et al; Senges et al. |
|  | *C3 acyl DFO-B* | 589.3915 | 589.3919 | -0.68 | C<sub>27</sub>H<sub>53</sub>N<sub>6</sub>O<sub>8</sub> | 2 | * | *This study* |
|  | *C4 acyl DFO-B* | 603.4073 | 603.4076 | -0.49 | C<sub>28</sub>H<sub>55</sub>N<sub>6</sub>O<sub>8</sub> | 2 | * | *This study* |
|  | *C5 acyl DFO-B* | 617.4233 | 617.4232 | 0.16 | C<sub>29</sub>H<sub>57</sub>N<sub>6</sub>O<sub>8</sub> | 2 | * | *This study* |
|  | *C4 acyl hydroxylated DFO-B* | 619.4024 | 619.4025 | -0.16 | C<sub>29</sub>H<sub>55</sub>N<sub>6</sub>O<sub>9</sub> | 3 | * | *This study* |
|  | *uC6 acyl DFO-B* | 629.4238 | 629.4232 | 0.95 | C<sub>30</sub>H<sub>57</sub>N<sub>6</sub>O<sub>8</sub> | 3 |  | *This study* |
|  | *C6 acyl DFO-B* | 631.4388 | 631.4389 | -0.16 | C<sub>30</sub>H<sub>59</sub>N<sub>6</sub>O<sub>8</sub> | 2 | * | *This study* |
|  | *uC7 acyl DFO-B* | 643.4389 | 643.4389 | 0.00 | C<sub>31</sub>H<sub>59</sub>N<sub>6</sub>O<sub>8</sub> | 3 |  | *This study* |
|  | *C7 acyl-DFO* | 645.4550 | 645.4545 | 0.77 | C<sub>31</sub>H<sub>61</sub>N<sub>6</sub>O<sub>8</sub> | 2 |  | Traxler et al;<br>Sidebottom et al. |
|  | *C8 acyl-DFO* | 659.4697 | 659.4702 | -0.76 | C<sub>32</sub>H<sub>63</sub>N<sub>6</sub>O<sub>8</sub> | 2 |  | Traxler et al;<br>Sidebottom et al. |

| Table 1 cont. Summary of siderophores identified using molecular networking and manual observations of the data. Nomenclature that are italicized were identified in this study. |  |  |  |  |  |  |  |  |
| --- | --- | --- | --- | --- | --- | --- | --- | --- |
| | Name | Measured mass<br>[M+H]<sup>+</sup> | Theoretical mass<br>[M+H]<sup>+</sup> | $\Delta$ ppm | Sum formula<br>[+H]<sup>+</sup> | ID<br>confidence | Monoculture | Reference |
| DFO-Bs | *uC10 acyl DFO-B* | 685.4856 | 685.4858 | -0.29 | C<sub>34</sub>H<sub>65</sub>N<sub>6</sub>O<sub>8</sub> | 3 |  | *This study* |
|  | *uC11 acyl DFO-B* | 699.5007 | 699.5025 | -2.57 | C<sub>35</sub>H<sub>69</sub>N<sub>6</sub>O<sub>8</sub> | 3 |  | *This study* |
|  | C11 acyl-DFO | 701.5174 | 701.5171 | 0.43 | C<sub>35</sub>H<sub>69</sub>N<sub>6</sub>O<sub>8</sub> | 2 |  | Traxler et al;<br>Sidebottom et al. |
|  | C12 acyl-DFO | 715.5322 | 715.5328 | -0.84 | C<sub>36</sub>H<sub>71</sub>N<sub>6</sub>O<sub>8</sub> | 2 |  | Traxler et al;<br>Sidebottom et al. |
|  | C13 acyl-DFO | 729.5496 | 729.5484 | 1.64 | C<sub>37</sub>H<sub>73</sub>N<sub>6</sub>O<sub>8</sub> | 2 |  | Traxler et al;<br>Sidebottom et al. |
|  | *uC12 acyl FO-B [M+Al]<sup>+</sup>* | 737.4776 | 737.4752 | 3.25 | C<sub>36</sub>H<sub>66</sub>N<sub>6</sub>O<sub>8</sub>Al | 3 |  | *This study* |
|  | *dodecanedioic DFO-B* | 745.5086 | 745.5070 | 2.15 | C<sub>36</sub>H<sub>69</sub>N<sub>6</sub>O<sub>10</sub> | 3 |  | *This study* |
|  | amphiphilic desferrioxamine<br>12 | 745.5438 | 745.5434 | 0.54 | C<sub>37</sub>H<sub>73</sub>N<sub>6</sub>O<sub>9</sub> | 2 |  | Sidebottom et al. |
|  | *tridecanedioic DFO-B* | 759.5221 | 759.5226 | -0.66 | C<sub>37</sub>H<sub>71</sub>N<sub>6</sub>O<sub>10</sub> | 3 |  | *This study* |
|  | amphiphilic desferrioxamine<br>15 | 759.5597 | 759.5590 | 0.92 | C<sub>38</sub>H<sub>75</sub>N<sub>6</sub>O<sub>9</sub> | 2 |  | Sidebottom et al. |
| DFO-Ds | desferrioxamine H | 461.2602 | 461.2606 | -0.87 | C<sub>20</sub>H<sub>37</sub>N<sub>4</sub>O<sub>8</sub> | 2 |  | Adapa et al. |
|  | *glutaric desferrioxamine H* | 475.2760 | 475.2762 | -0.42 | C<sub>21</sub>H<sub>39</sub>N<sub>4</sub>O<sub>8</sub> | 2 |  | *This study* |
|  | *C2 acyl glutaric*<br>*desferrioxamine H* | 489.2900 | 489.2919 | -3.88 | C<sub>22</sub>H<sub>41</sub>N<sub>4</sub>O<sub>8</sub> | 2 |  | *This study* |
|  | *desferrioxamine D4* | 559.3453 | 559.3450 | 0.50 | C<sub>25</sub>H<sub>47</sub>N<sub>6</sub>O<sub>8</sub> | 2 |  | *This study* |
|  | *deoxydesferrioxamine D3* | 587.3757 | 587.3763 | -1.02 | C<sub>27</sub>H<sub>51</sub>N<sub>6</sub>O<sub>8</sub> | 3 | * | *This study* |
|  | *Desferrioxamine D3* | 589.3915 | 589.3919 | -0.68 | C<sub>27</sub>H<sub>53</sub>N<sub>6</sub>O<sub>8</sub> | 2 | * | *This study* |
|  | desferrioxamine D1 | 603.3714 | 603.3712 | 0.33 | C<sub>27</sub>H<sub>51</sub>N<sub>6</sub>O<sub>9</sub> | 2 | * |  |
|  | *C2 acyl DFO-D* | 617.3864 | 617.3869 | -0.81 | C<sub>28</sub>H<sub>53</sub>N<sub>6</sub>O<sub>9</sub> | 2 |  | *This study* |
|  | *C4 acyl DFO-D* | 645.4185 | 645.4182 | 0.46 | C<sub>30</sub>H<sub>57</sub>N<sub>6</sub>O<sub>9</sub> | 2 | * | *This study* |
|  | *C5 acyl DFO-D* | 659.4331 | 659.4338 | -1.06 | C<sub>31</sub>H<sub>59</sub>N<sub>6</sub>O<sub>9</sub> | 2 |  | *This study* |
|  | *C6 acyl DFO-D* | 673.4498 | 673.4495 | 0.45 | C<sub>32</sub>H<sub>61</sub>N<sub>6</sub>O<sub>9</sub> | 2 | * | *This study* |
|  | *C7 acyl DFO-D* | 687.4655 | 687.4651 | 0.58 | C<sub>33</sub>H<sub>63</sub>N<sub>6</sub>O<sub>9</sub> | 2 | * | *This study* |
|  | *C11 acyl DFO-D* | 743.5280 | 743.5277 | 0.40 | C<sub>37</sub>H<sub>71</sub>N<sub>6</sub>O<sub>9</sub> | 2 | * | *This study* |
|  | *C12 acyl DFO-D* | 757.5436 | 757.5434 | 0.26 | C<sub>38</sub>H<sub>73</sub>N<sub>6</sub>O<sub>9</sub> | 2 | * | *This study* |
|  | *C13 acyl DFO-D* | 771.5607 | 771.5590 | 2.20 | C<sub>39</sub>H<sub>75</sub>N<sub>6</sub>O<sub>9</sub> | 2 | * | *This study* |
|  | *C14 acyl DFO-D* | 785.5749 | 785.5747 | 0.25 | C<sub>40</sub>H<sub>77</sub>N<sub>6</sub>O<sub>9</sub> | 2 | * | *This study* |
| tenacibactins | tenacibactin C | 503.3070 | 503.3075 | 0.99 | C<sub>23</sub>H<sub>43</sub>N<sub>4</sub>O<sub>8</sub> | 3 |  | Jang et al. |
|  | *tenacibactin E* | 531.3380 | 531.3388 | -1.15 | C<sub>25</sub>H<sub>47</sub>N<sub>4</sub>O<sub>8</sub> | 3 | * | *This study* |
|  | *tenacibactin F* | 545.3545 | 545.3545 | 0.00 | C<sub>26</sub>H<sub>49</sub>N<sub>4</sub>O<sub>8</sub> | 3 | * | *This study* |
|  | *tenacibactin G* | 587.3990 | 587.4014 | 4.08 | C<sub>29</sub>H<sub>55</sub>N<sub>4</sub>O<sub>8</sub> | 3 |  | *This study* |
|  | *tenacibactin H* | 601.4168 | 601.4171 | -0.49 | C<sub>30</sub>H<sub>57</sub>N<sub>4</sub>O<sub>8</sub> | 3 | * | *This study* |
|  | *tenacibactin I* | 615.4326 | 615.4327 | -0.16 | C<sub>31</sub>H<sub>59</sub>N<sub>4</sub>O<sub>8</sub> | 3 |  | *This study* |
|  | *tenacibactin J* | 629.4484 | 629.4484 | 0.00 | C<sub>32</sub>H<sub>61</sub>N<sub>4</sub>O<sub>8</sub> | 3 |  | *This study* |

### Acyl DFO production by *Streptomyces* sp. S29

*Streptomyces* sp. S29 was able to produce twelve new acyl DFO-Bs similar to those previously described.^26,27^ Of these 12, five were short acyl chain derivatives (C_3_, C_4_, C_4_ hydroxylated, C_5_, and C_6_), five were monounsaturated derivatives (uC_6_, uC_7_, uC_10_, uC_11_, and uC_12_), and two were derivatives containing glutaric acid at the C-terminus, similar to those seen by Rütschlin et al.^40^ Pertaining to the unsaturated derivatives, placement of the alkene at the position α to the carbonyl is based off previous studies also reporting DFOs containing unsaturated acyl chains.^27,41^ Additionally, the position of the hydroxy in the C_4_ derivative was based on previous work showing multiple amphiphilic peptide-based siderophores contain hydroxylation at the β-position.^27,42^

We propose henceforth that the previously known ‘acyl DFOs’ should be renamed ‘acyl DFO-B’ due to the observation in this study of analogues that contain a *N*-acetyl-*N*-hydroxycadaverine terminus, similar to desferrioxamine D. The new acyl DFO-D analogues discovered here also range from short, medium and long acyl chained derivatives (Table 1). It is likely the acyl DFOs are branched analogues as previous studies noted that it was unlikely the chains were fully linear since *Streptomyces* spp. utilise branched starter units in fatty acid synthesis.^26,27^ This is also the case for the suite of tenacibactin analogues observed in the data. These branched acyl chain siderophores were originally observed in *Tenacibaculum* sp. A4K-17 ^38^, and are essentially truncated versions of acyl DFO-Bs that lack the N-terminal *N*-hydroxycadaverine.

*Streptomyces coelicolor* has shown broad substrate specificity in regard to medium and long acyl chain incorporation into desferrioxamine structures^19^, but not short acyl chained groups, where *Streptomyces* sp. S29 can facilitate production of short, medium, and long acyl chained DFO-B analogues. Traxler et al. showed the ability of other Actinobactera to react to the presence of *S. coelicolor*. through the production of acyl DFO-B analogues.^26^ Interestingly, we see this same phenomenon for the coculture of *Streptomyces* sp. S29 and the fungi. Medium and long chained acyl DFO-Bs (including unsaturated derivatives) were almost exclusively produced in coculture whereas short, medium and long chained acyl DFO-Ds were produced under both mono-and coculture conditions. The tenacibactin analogues were distributed amongst both cultures (Table 1). It was hypothesized these long-chained derivatives might be a way for the uptake of these analogues to bypass the DesE uptake mechanism, instead they ‘hand-off’ the Fe(III) via membrane bound ferrioxamines.^19,43^

### Aryl DFO production by *Streptomyces* sp. S29

The production of seven new aryl DFOs related to legonoxamine A and B, which were also observed in this study (Table 1), shows the biosynthetic flexibility of *Streptomyces* sp. S29. These structures range from being smaller, potential oligomers like legonoxamine C (*m/z* 351.1912, Figure S12) to the larger legonoxamine H (*m/z* 665.3893, Figure S17). A few interesting legonoxamine derivatives include legonoxamine F (Figure S15), putrescine-containing legonoxamine G (Figure S15) and legonoxamine A glycoside (Figure S18).

Legonoxamine F contains one 1,6-diaminohexane instead of the typical 1,5-diaminopentane. This is a well-established phenomenon in siderophore biosynthesis^31,44^ linking it to the promiscuity of the S29 *des* biosynthetic gene cluster. Legonoxamine G contains a *N*-hydroxyputrescine moiety due to the initial loss of 104 Da instead of the typical 118 Da associated to *N*-hydroxycadaverine. Putrebactin is the archetypal putrescine containing siderophore^45^, and as its biosynthetic gene cluster has been shown to lack genes coding for production of putrescine, it is assumed that instead, this precursor is sequestered from metabolite pools.^46^ Interestingly, researchers have also found that under putrescine-depleted conditions, *Shewanella putrefaciens* shifts to producing desferrioxamine B when cadaverine precursors are available.^47^ In desferrioxamine biosynthesis, incorporation of the putrescine precursor in desferrioxamine A1 has been shown before in multiple species^30,44,48^ and now in this study with the discovery of legonoxamine G.

Legonoxamine A glycoside was observed using the predicted molecular formula (C_37_H_62_N_6_O_13_) and the loss of a sugar (−162 Da). An additional putative diglycosylated DFO (*m/z* 899.4827), with a predicted molecular formula of C_38_H_70_N_6_O_18_, was observed in the mass spectrum but upon selected ion monitoring (SIM), produced insufficient fragment ions for structural characterisation. The production of glycosylated DFOs has only been reported once before: nocardamin glucuronide, isolated from *Streptomyces* sp. 80H647.^49^ There are only a few reports of microbial enzymes that produce glucuronides^50^, yet, the addition of glucose or other hexoses are more commonplace in bacterial natural products. We propose legonoxamine A glycoside undergoes a similar glycosylation mechanism to nocardamin glucuronide, occurring on a *N*-hydroxyl.

### Macrocyclised desferrioxamines

We observe desferrioxamine E in the molecular network (Figure S1) as well as several other macrocyclic DFOs. In Figure 3, these are anchored by bisucaberin 01 (*m/z* 443), first described by Senges et al.^28^ Based on the molecular formulas, fragmentation patterns and the previously established structure, we observe multiple additional macrocyclic derivatives connected via bisucaberin 01 but their structures were not definable based on MS/MS data alone (Table 1). The putative macrocyclised derivatives extend by successive CH_2_’s, differing from the study by Senges et al. which reduce in size, but the location of this extension requires confirmation via NMR. Similar to acyl DFOs, macrocyclic derivatives have also shown to have flexible biosynthesis in feeding studies, including the incorporation of 1,8-diaminooctane moieties.^51^ There was no observation of putrebactin, or avaroferrin in the culture despite the similar biosynthesis with bisucaberins and desferrioxamines.

### Dicarboxylic acid desferrioxamine analogues

Two new desferrioxamines, observed only in coculture extracts, with dicarboxylic acid C-termini were also elucidated. Dodecanedioic DFO-B (Figure S33) has an *m/z* of 745.5086, corresponding to the molecular formula of C_36_H_68_N_6_O_10_. Tridecanedioic DFO-B (Figure S34) has an *m/z* of 759.5221, corresponding to the molecular formula of C_37_H_70_N_6_O_10_. This is the first reported instance of large dicarboxylic acids being incorporated into DFO structures yet there are examples of larger dicarboxylic acids being bound to CoA, like pimelic acid, therefore we cannot rule the possibility out.

Microbial cytochrome p450s are responsible for adding complexity to fatty acids to be used as building blocks for more complex metabolites or as signalling molecules.^52^ Previous studies have shown the bacterial production of dicarboxylic acids might occur via cytochrome p450 monoxygenase.^53^ In order to determine the presence of enzymes that facilitate ω-oxidation in S29, protein sequences that belong to the long-chain fatty acid ω-monooxygenase enzyme class (EC 1.14.14.80) and from fatty acid ω-oxidation gene ontology (GO:0010430) were obtained from the Uniprot database. BLAST results of these sequences against the S29 draft genome reveals just one hit (supplementary data), comprising 3 orthologs with mid-term amino acid identity (40-50%), a methyl-branched lipid ω-hydroxylase (cyp124) from Mycobacterium spp. (~43% identity) (Uniprot: P9WPP2, P0A517 & P9WPP3). Pfam sequence search confirms cytochrome P450 domain for this protein and further experimentation could confirm its ω-monooxygenase activity. Searching Uniprot under these search criteria returns only a handful of bacterial sequences with a couple belonging to Actinobactera, therefore, the possibilities to track this activity in the S29 genome are very narrow resulting in an inconclusive result regarding the presence of an ω-monooxygenase in S29.

### Ferrioxamines and DFO presence in pure and coculture experiments

A number of metal complexed ferrioxamines were observed in subnetwork 2 (Figure 6). Fe(III) and Al^3+^ are the only metals complexed with the variety of desferrioxamines present, similar to studies before.^26,27^ Although the binding affinity for Fe(III) is greater than Al(III)^54^, it appears that this difference in affinity is not enough to prevent Al(III) chelation. The importance of iron has already been addressed in regard to bacteria and cell maintenance but Al(III) is toxic and has no direct role in metabolism and seems to be an consequence of the chemical ecology of these metabolites.

The molecular network was also used to evaluate the production of the DFOs between the two cocultivations with *A. niger* and *B. cinerea* and the *Streptomyces* sp. S29 monoculture (Figure S3). The results showed that 32 of the 71 ions were present in the monoculture, whereas all of the remaining were found in coculture extracts only (Table 1). There was no qualitative significant difference in the siderophores produced under cocultivation with *A. niger* compared to those produced in cocultivation with *B. cinerea*, indicating the potential broad response to a fungal microbe in the environment. These results might point to DFO and siderophore production in a broader context as a common chemical response to other microbes, whether it be bacterial or fungal, in close environmental proximity to the primary organism.

### MS2LDA analysis

In order to further evaluate the overall chemical space that DFOs inhabit in the molecular network, MS2LDA was performed. MS2LDA is a technique which identifies molecular ‘fingerprints’ in fragmentation patterns and assigns these into motifs (13 Mass2Motifs were manually curated). By merging the two networks (GNPS molecular network and MS2LDA) after motifs were annotated, it is clear that further DFO chemical space is there to be discovered. Figure S4 represents the overall results of this merged network where we have highlighted the previously identified features (107) and edges (970) from the original molecular networking job and all of the nodes (351) and edges (1991) that contain DFO motifs identified through MS2LDA. This near tripling of features by inclusion of the MS2LDA workflow doesn’t necessarily correlate to an increase in metabolites, but upon validation of the data, there seems to be very minimal features corresponding to isotopes or adducts.

In order to evaluate the accuracy of this analysis, a characterized DFO in this study (legonoxamine E, Figure 5) and an unknown DFO observed using MS2LDA (*m/z* 650.37, [M+2H]^2+^) were more closely analysed (Figure S5). The unknown desferrioxamine is a singleton (Figure S5B) in both the original network and the merged network, yet, MS2LDA identifies it contains DFO motifs and upon closer inspection and prediction of its molecular formula (C_59_H_103_N_12_O_20_). The potential of this putative DFO being a dimer was ruled out based on fragmentation analysis lacking the monomer. Elucidation of its full structure was not able to be completed with MS/MS data alone, therefore, further studies are underway to characterise its structure. Nonetheless, this example displays the utility of MS2LDA to evaluate the chemical potential of a bacterium and its potential use in these types of studies in the future.

**Figure 5.**
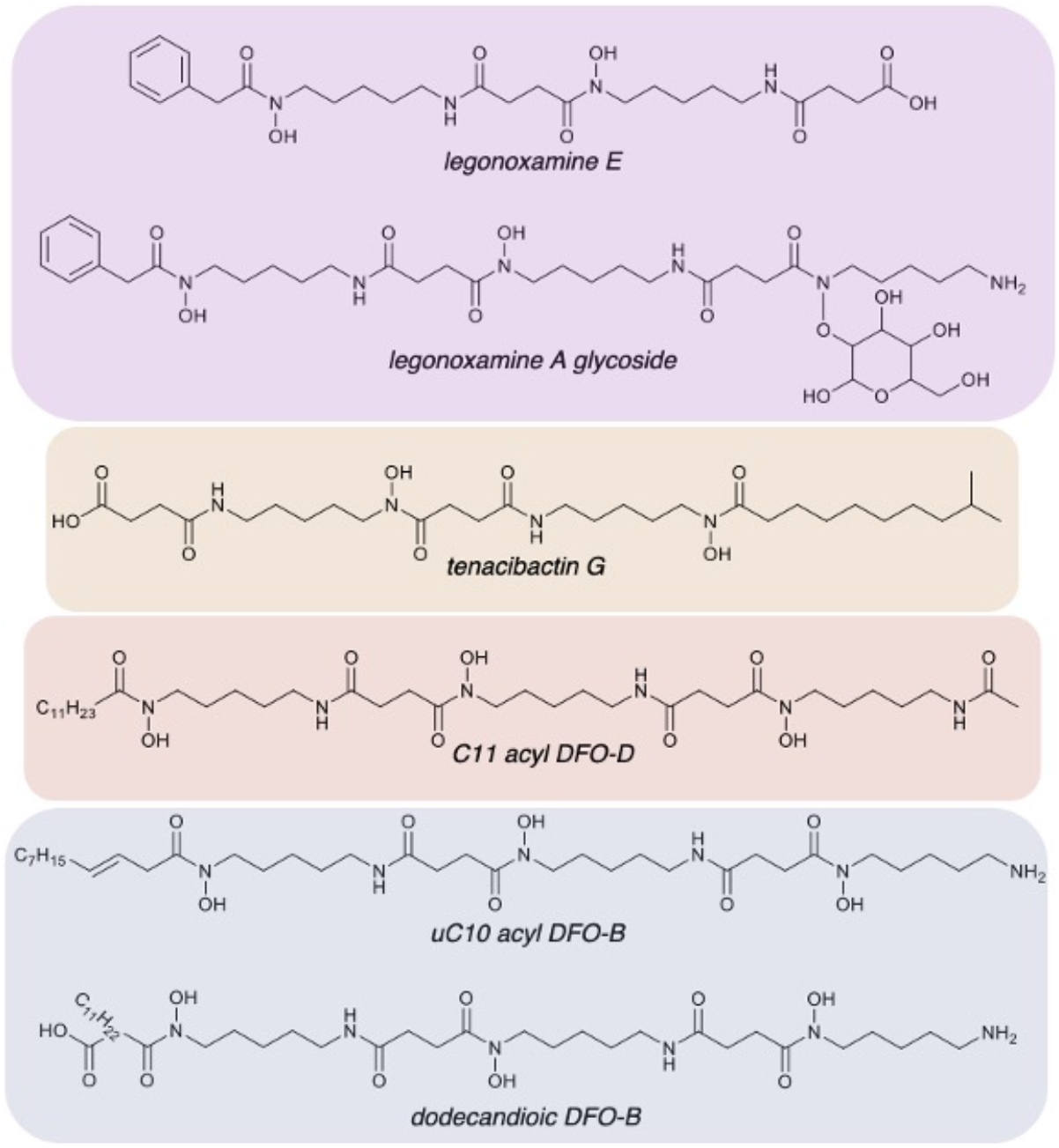
Representation of desferrioxamine structural diversity of *Streptomyces* sp. S29. (A-B) New legonoxamine derivatives (purple), tenacibactins (orange), DFO-Ds (red), and a variety of DFO-Bs (blue) were all found. Full MS/MS data and annotations can be found in supplementary for all 41 new analogues discovered.

**Figure 6.**
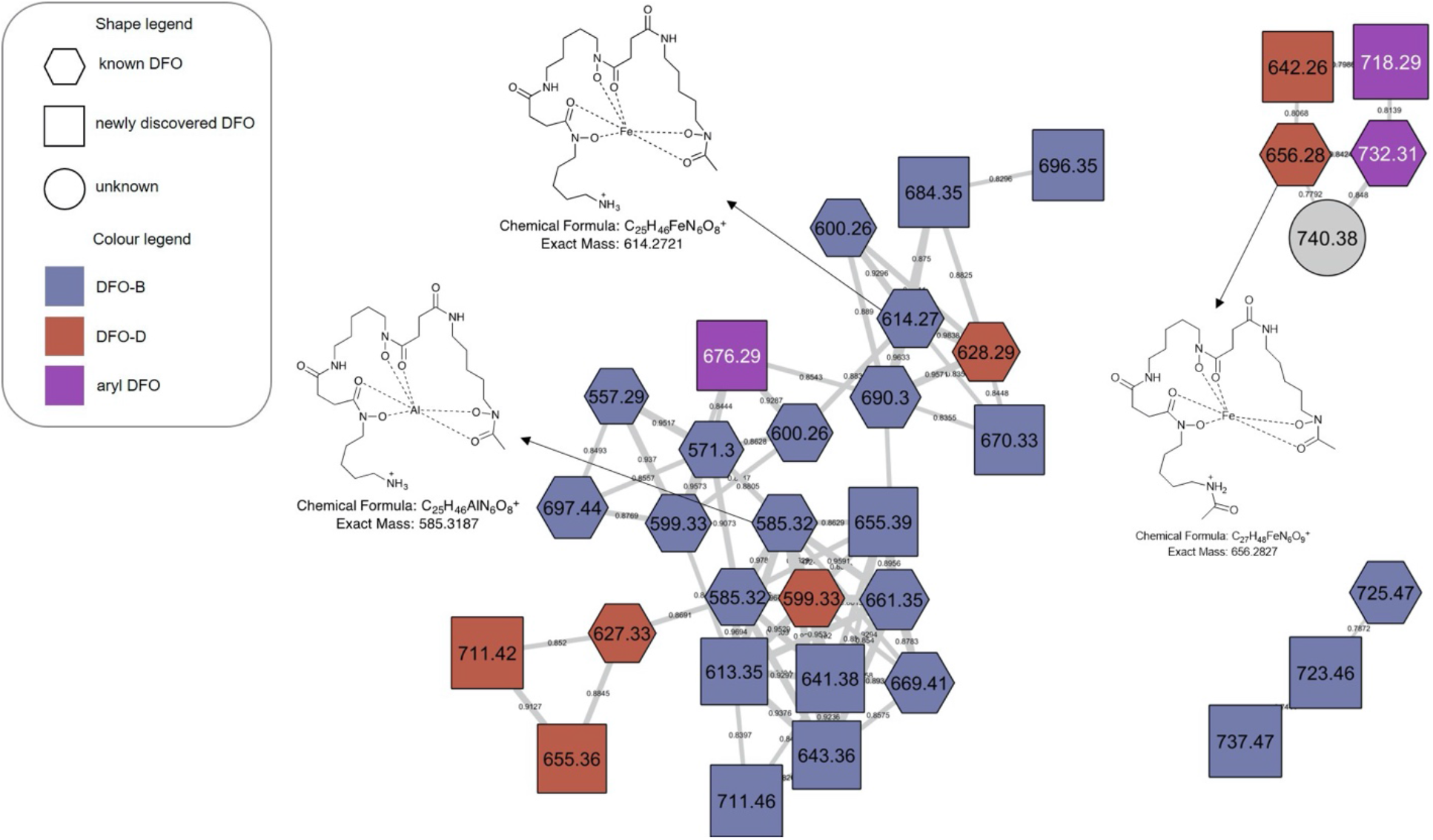
Metal-complexed molecular families. The GNPS library was able to identify multiple metal ion complexes (hexagonal nodes). All other features observed in the molecular network were assignable as Fe(III) or Al(III) bound ferrioxamines. Al(III) analogue of new desferrioxamine analogue –uC12 acyl FO-B. Table S1 contains accurate mass information on all complexed ions found in this figure.

### *Streptomyces* sp. S29 siderophore biosynthetic gene clusters

DFOs are biosynthesized by nonribosomal peptide synthetase independent siderophore (NIS) pathways and the presence of at least one gene encoding a homolog to IucA/IucC has become recognized as a marker of NIS biosynthetic gene clusters.^55^ DFO biosynthesis is encoded by the highly conserved *des* biosynthetic gene cluster (*desEFABCD* genes) found in many *Streptomyces* species ^21^. Using FeGenie^56^, the identification of iron-related genes revealed 27 siderophore biosynthesis and 11 transport related genes in the *Streptomyces* sp. S29 genome. These siderophore biosynthesis related genes are grouped in five biosynthetic gene clusters and antiSMASH ^57^ predictions reveal that three of the five contain at least one gene homolog to IucA/IucC and the other two correspond to NRPS dependent pathways.

Region 7 (Figure S53) contains a siderophore BGC in *Streptomyces* sp. S29 with high homology to *desA*-*D* genes and to the ferric-siderophore uptake and utilization genes (*desE*-*F*). This *des* cluster follows the same structure as many *Streptomyces* spp., including the position of iron boxes upstream putative *desE* (s29_002291) and *desA* (s29_002293) genes (Fig S31). DesC of the *des* BGC from *Streptomyces* sp. S29 (s29_2295) exhibits high identity to DesC from S. coelicolor (72%) and to LgoC from *Streptomyces* sp. MA37 (69%). In *S. coelicolor* A3(2), transcription of *des* BGC is repressed by the divalent metal-dependent regulatory (DmdR) protein DmdR1 and derepressed by iron limitation.^58^ BLASTN/P analysis revealed a putative DmdR repressor in *Streptomyces* sp. S29 (s29_002782) with 89% nucleotide and 93% amino acid identity to DmdR1 (AJ271797.1 / CAC28070.1). BLASTN analyses identified eight putative DmdR-binding sites (iron boxes) in *Streptomyces* sp. S29 by query of putative iron boxes found upstream in different ORFs in *S. coelicolor* A3(2) genome.^19^

Acyltransferases involved in hydroxamate biosynthesis normally exhibit narrow substrate tolerance (e.g. IucB, which shows a high degree of specificity towards acetyl-CoA).^19^ The array of DFOs produced by *S. coelicolor* (B, E, G1 and amphiphilic) and aryl DFOs (legonoxamine A-B) produced by *Streptomyces* sp. MA37, has been attributed to substrate tolerant acyltransferases DesC and LgoC. ^19,59^ Senges et al. acknowledged in their evaluation of the *Streptomyces chartreusis* secretome that in iron depleted minimal medium they also observed additional desferrioxamines produced and that the overall biosynthetic potential may be much broader than what has been predicted.^28^ Cloning and more detailed analysis of the *Streptomyces* sp. S29 *des* gene cluster is ongoing to link the chemical potential shown in this study.

In conclusion, the chemistry observed in the *Streptomyces* sp. S29 secretome shows it potential as an agent to prevent fungal infection, evidenced by its locking away of Fe(III) via overproduction of desferrioxamines. New analogues were observed in both monoculture and cocultivation studies with fungal phytopathogens, *A. niger* and *B. cinerea*. *Streptomyces* sp. S29 seems to produce an arsenal of desferrioxamine analogues using any available precursor materials available whether that be putrescine, cadaverine, aryl, acyl, or long chained dicarboxylates. The role of bacterial communities in plant and crop protection is becoming more important due to the growing world population, with a strong emphasis on improving strategies at preventing phytopathogenic infection. Overall, the production of 46 new and 25 previously reported siderophores indicates this *Streptomyces* sp. should be studied further for its plant protective properties as a biocontrol agent.

## EXPERIMENTAL SECTION

### Strain collection

*Streptomyces* sp. S29 was isolated from a soil sample collected from Lupine plant (*Lupinus oreophilus*) roots originating from a polyextreme enviroment at Quebrada Nacimiento, Socaire, San Pedro de Atacama, Atacama Desert (23°37’06’’S;67°50’56’’W-3,646 m.a.s.l). One aliquot was heated at 100 °C for 5 min and directly sprinkled onto Gause #1 and Starch-Casein plates, both complemented with nystatin, cycloheximide and nalidixic acid. Another aliquot was resuspended in ¼ Ringer solution, heated at 60 °C for 10 min and dilutions of 10^−1⁄2^ and 10^−1^ were spread onto agar plates. Plates were incubated at 30 °C for ~1 week and colonies were subcultured on ISP2 plates. Based on morphological characteristics of the colony and antifungal activity experiments against *B. cinerea* and *A. niger*, 16S rRNA sequence of S29 isolate was used to identify and confirm its taxonomy using EzBioCloud database (3) (data not shown).

### Phytopathogens

*Aspergillus niger* and *Botrytis cinerea*, biochemically characterized field samples, were provided by Laboratorio de microbiología, Facultad de Ciencias naturales, matemática y medio ambiente, Universidad Tecnológica Metropolitana (UTEM), Santiago, Chile. Molecular identification of these strains was carried out by sequencing of internal transcribed spacer (ITS) region amplified by PCR using universal primers ITS1 and ITS4 (supplementary data).

### Biological screening

Preliminary antifungal screening of the lupine rhizosphere *Streptomyces* collection was carried out starting from spore stocks, where Streptomyces isolates were grown on ISP2 at 30°C for 1 week. On PDA plates, two inverted agar plugs (different isolates) from these ISP2 plates, set at 2 cm from the centre, were grown at 30°C O.N. After this, the centre of the PDA plates was inoculated using 10 μL of a 5×10^4^ spores/mL spore stock of a phytopathogen (*Botrytis/Fusarium*) and grown at 25°C. Inhibition observations were carried out at 4, 11 and 17 days from pathogen inoculation.

Disc diffusion assays were adapted for the microbes tested using the Clinical and Laboratory Standards Institute M44 Method for Antifungal Disk Diffusion Susceptibility Testing of Yeasts.^60^ Extract concentrations were standardised to 200 μg/mL for all bioassays conducted.

In order to conduct post-fermentation streaks, 50 μL of fermentation was aseptically transferred onto a ISP2 agar plate. 24- and 48-hour incubation (28°C) intervals were checked for fungal growth. Due to the faster growth rate of the fungi, we would expect any presence of fungal spores to lead to the observation of mycelial growth, yet no fungal growth was observed.

### Cocultivation and extraction

A set of experiments were carried out on these three organisms in order to determine the effect of the fungal culture being added to the *Streptomyces* sp. (a) *Streptomyces* sp. S29 alone: a single colony was aseptically transferred to 100 mL of iron deficient medium (ISP2), shaken at room temperature at 150 RPM for 7 days, in order to induce production of siderophores. (b) *Streptomyces* sp. S29 + *Aspergillus niger*: 100 mL of bacterial culture was grown for 5 days, at room temperature and shaken at 150 RPM. On the second day of bacterial growth, an independent axenic culture of *A. niger* (spores) was grown for 3 days (10 mL of PDA liquid medium, room temperature and 150 RPM shaking). On the fifth day of bacterial growth, the fungal liquid culture was transferred axenically to the bacterial culture and allowed to grow an addition 2 days. (c) *Streptomyces* sp. S29 + *Botrytis cinerea*: the same procedure as described in experiment b was followed. Additionally, each experiment contained 3 g/50 mL of sterilised glass beads to prevent aggregation. Each coculture was then scaled up to 3 L total volume in an attempt to produce sufficient secondary metabolites for structure elucidation.

Each liquid fermentation included Diaion HP-20 resin (3 g/50 mL) that was sterilised with culture medium. After 7 days, the resin was vacuum filtered, subsequently washed five times with milli-q water and finally macerated exhaustively in methanol (3×). The methanolic extract was then dried on a rotary evaporator and subsequently underwent Kupchan partitioning.^61^ Finally, the *sec*-butanol fraction underwent C18 solid-phase extraction (Phenomenex), with each separation yielding 5 fractions of increasing methanol and these samples were subsequently prepared for LC-MS studies.

The 100% methanol fractions from the *Botrytis* and *Aspergillus* co-culture were subjected to further separation via reversed phase-HPLC with a Waters Sunfire C18 semi-preparative column (5 μM, 100Å, 250 × 10 mm) wand a solvent system of A–95/5 MeOH/H_2_O and B-MeOH. The Agilent 1200 HPLC utilized a solvent gradient of 0% B to 100% B at 2.0 mL/min over 50 min.

### Liquid chromatography – mass spectrometry

All runs were analysed using an Agilent 1290 UHPLC coupled to Bruker MAXIS II Q-ToF mass spectrometer. LC utilizes a Phenomenex Kinetex XB-C_18_ (2.6 mM, 100 × 2.1 mm) column with a mobile phase of 5% MeCN+0.1% formic acid to 100% MeCN+0.1% formic acid in 11 min. Bruker MAXIS II has a mass range of *m/z* 100-2000, capillary voltage 4.5 kV, nebulizer gas 5.0 bar, dry gas 12.0 l/min, and dry temperature of 220 °C. MS/MS experiments were conducted under Auto MS/MS scan mode with a step collision energy from 80-200%. Samples were prepared at a concentration of 0.3 mg/mL. Raw data files were analysed using Bruker Daltonics 3.6.

### GNPS Molecular Networking and MS2LDA

A molecular network was created using the online workflow (https://ccms-ucsd.github.io/GNPSDocumentation/) on the GNPS website (http://gnps.ucsd.edu). The data was filtered by removing all MS/MS fragment ions within +/− 17 Da of the precursor m/z. MS/MS spectra were window filtered by choosing only the top 6 fragment ions in the +/− 50Da window throughout the spectrum. The precursor ion mass tolerance was set to Da and a MS/MS fragment ion tolerance of 0.02 Da. A network was then created where edges were filtered to have a cosine score above 0.7 and more than 5 matched peaks. Further, edges between two nodes were kept in the network if and only if each of the nodes appeared in each other’s respective top 10 most similar nodes. Finally, the maximum size of a molecular family was set to 100, and the lowest scoring edges were removed from molecular families until the molecular family size was below this threshold. The spectra in the network were then searched against GNPS’ spectral libraries. The library spectra were filtered in the same manner as the input data. All matches kept between network spectra and library spectra were required to have a score above 0.7 and at least 6 matched peaks. All features originating from the medium/instrument blanks were excluded from the final molecular network.

For documentation on running MS2LDA, the user is referred to the original publication and guidelines, which are also present at ms2lda.org.^62^ All spectral images in the supplementary information were created using Metabolomics Spectrum Resolver (https://metabolomics-usi.ucsd.edu/).^63^

All siderophores dereplicated and elucidated in this study can be found in Table 1. The structure elucidation of each new derivative and its corresponding MS/MS spectra were generated using Metabolomics Spectrum Resolver are displayed in the supplementary information (Table S2 and Figures S7-S52). All MS/MS spectra were also annotated and deposited in the GNPS library, as well as the original datasets (Table S2).

### All data files and workflows can be found at the links below

GNPS molecular networking job: https://gnps.ucsd.edu/ProteoSAFe/status.jsp?task=3c73305ef7ed48e19cb18e50b8b6b2bd

MS2LDA annotation experiment: http://ms2lda.org/basicviz/summary/1319/

MASSIVE: doi:10.25345/C5772V

### Streptomyces sp. S29 genome sequencing and assembly

*Streptomyces* sp. S29 total DNA was extracted from liquid culture using DNeasy UltraClean Microbial Kit (Qiagen). Whole genome sequencing (WGS) of the strain was performed by an Illumina MiSeq (2 x 300 PE). Nanopore sequencing was performed using a Rapid Sequencing Kit (SQK-RAD004) and MinIon Flow Cell (R9.4.1). The sequence was assembled using Canu, polished using Racon, Illumina dataset was mapped onto the contigs using BWA mem and then corrected using Pilon. The genome was annotated using NCBI pgap software (6).

## ASSOCIATED CONTENT

### Supplementary Information

Tables of known and new desferrioxamine derivatives and their corresponding CCMS Library IDs and Metabolomics Resolver Images (Table S2), disc diffusion assay results, desferrioxamine containing subnetworks, new desferrioxamine analogues with annotated structures and MS^2^ spectra, ITS regions for fungal phytopathogens, cytochrome p450 enzyme sequence.

### Author Contributions

DLS^†^ and ED^†^ contributed equally to this work.

### Conflicts of interest

There are no conflicts of interest to declare.

## Acknowledgements

SAJ would like to thank the University of Aberdeen for funding doctoral studies through and Elphinstone Scholarship. DL would like to thank the Agencia Nacional de Investigación y Desarrollo (ANID) for funding doctoral studies through ‘Beca nacional de doctorado’ Scholarship. ED would like to thank the Ministry of Higher Education and Scientific Research - Sudan, together with the University of Khartoum, for joint funding of master’s studies. We would like to thank to Valeria Razmilic and Jean Franco Castro for their valuable advice and work in the setup of Lupine *Streptomyces* culture collection. We would also like to thank the support team at GNPS and Justin J.J. van der Hooft at MS2LDA for help with data deposition and for help at any stage of running the relevant workflows.

## Notes

### Competing Interest Statement

The authors have declared no competing interest.

