## Supplementary information for "Iron-meditated fungal starvation by lupine rhizosphere-associated and extremotolerant *Streptomyces* sp. S29 desferrioxamine production"

### Table of Contents


Table S1. Summary of metal complexed desferrioxamines ........................................ S3

Table S2. Summary of CCMS library and Metabolomics Spectrum Resolver Identifiers ........................................ S4-S6

Figure S1. Additional DFO containing molecular families ........................................ S7

Figure S2. Structure-centric whole molecular network ........................................ S8

Figure S3. Coculture-centric whole molecular network ........................................ S9

Figure S4. MS2LDA merged molecular network ........................................ S10

Figure S5. MS2LDA Mass2Motifs examples and proof of concept ........................................ S11

Figure S6. Disc diffusion-based bioassay-guided fractionation ........................................ S12

Figure S7-S52. Annotated desferrioxamine structures and MS/MS spectra for new analogues ........................................ S12-S58

Figure S53. *Streptomyces* sp. S29 des biosynthetic gene cluster ........................................ S59

ITS regions for fungal phytopathogens ........................................ S60

S29 omega-monoxygenase BLAST hit ........................................ S61

Table S1. Ferrioxamine and Al(III)-complexed metabolites observed in Figure 6.

| | $[M-2H+Fe]^{+}$ | | | $[M-2H+Al]^{+}$ | | |
|---|---|---|---|---|---|---|
| Name | Predicted | Observed | Δ ppm | Predicted | Observed | Δ ppm |
| Des A2 / IC202B | - | - | - | 557.2881 | 557.288 | 0.18 |
| ferrioxamine A1 | 600.2572 | 600.2565 | -1.17 | 571.3038 | 571.3033 | 0.88 |
| ferrioxamine B | 614.2728 | 614.2723 | -0.81 | 585.3194 | 585.3188 | 1.03 |
| ferrioxamine N | 628.2885 | 628.2883 | -0.32 | 599.3351 | 599.3349 | 0.33 |
| C3 acyl FO-B | | | | 613.3500 | 613.3460 | 6.50 |
| ferrioxamine D3 | 642.2670 | 642.2630 | 6.22 | | | |
| ferrioxamine D1 | 656.2834 | 656.2822 | -1.83 | 627.3300 | 627.3287 | 2.07 |
| C4 acyl FO-B | 656.3191 | 656.3200 | -1.37 | | | |
| C5 acyl FO-B | 670.3354 | 670.3341 | -1.94 | 641.3820 | 641.3807 | 2.03 |
| C4 hydroxyl acyl FO-B | - | - | - | 643.3613 | 643.3599 | 2.18 |
| legonoxamine G | 676.2885 | 676.2871 | -2.07 | 647.3351 | 647.3338 | 2.01 |
| C3 acyl FO-D | | | | 655.3606 | 655.3560 | -7.02 |
| C6 acyl FO-B | 684.3511 | 684.3497 | -2.05 | 655.3977 | 655.3937 | 6.10 |
| acyl ferrioxamine 1/ legonoxamine A | 690.3041 | 690.3027 | -2.03 | 661.3507 | 661.3492 | 2.27 |
| uC7 acyl FO-B | 696.3504 | 696.3480 | 3.44 | | | |
| C7 acyl FO-B | 698.3667 | 698.3655 | -1.72 | 669.4133 | 669.4106 | 4.03 |
| C9 acyl FO-B | | | | 697.4439 | 697.4420 | -2.72 |
| C7 acyl FO-D | - | - | - | 711.4239 | 711.4199 | 5.62 |
| C10 acyl FO-B | - | - | - | 711.4592 | 711.4573 | 2.60 |
| legonoxamine H | 718.2983 | 718.2940 | 5.98 | | | |
| uC11 acyl FO-B | - | - | - | 723.4613 | 723.4622 | -1.18 |
| C11 acyl FO-B | - | - | - | 725.4759 | 725.4745 | 1.93 |
| acyl ferrioxamine 2 | 732.3140 | 732.3130 | 1.36 | | | |
| *uC12 acyl FO-B | | | | 737.4769 | 737.4730 | -5.28 |

Table S2. CCMS Library IDs and Metabolomics Resolver Image IDs for metabolites found in this study.

| Name | Measured mass [M+H]+ | CCMS Library ID | Metabolomics Resolver ID |
|---|---|---|---|
| **macrocylised desferrioxamines** | | | |
| Bisu-03 | 429.2344 | CCMSLIB00003739989 | mzspec:GNPSTASK-4b53c4f871bb416fb5ed51300d6c8d29:spectra/specs_ms.mgf:scan:2258 |
| Bisu-01 | 443.2495 | CCMSLIB00003739987 | mzspec:GNPSTASK-a172e23b485b4fbea34650f2bf3c0ec2:spectra/specs_ms.mgf:scan:580 |
| desferrioxamine E | 601.3557 | CCMSLIB00000001621 | mzspec:GNPSTASK-4b53c4f871bb416fb5ed51300d6c8d29:spectra/specs_ms.mgf:scan:1028 |
| **aryl desferrioxamines** | | | |
| legonoxamine B | 337.1754 | CCMSLIB00005723622 | mzspec:GNPSTASK-a172e23b485b4fbea34650f2bf3c0ec2:spectra/specs_ms.mgf:scan:1539 |
| legonoxamine C | 351.1912 | CCMSLIB00005723608 | mzspec:GNPSTASK-16d50fb5de564636964680ad8c037b7c:spectra/specs_ms.mgf:scan:827 |
| legonoxamine D | 437.2755 | CCMSLIB00005723623 | mzspec:GNPSTASK-a172e23b485b4fbea34650f2bf3c0ec2:spectra/specs_ms.mgf:scan:2064 |
| legonoxamine E | 537.2924 | CCMSLIB00005723624 | mzspec:GNPSTASK-a172e23b485b4fbea34650f2bf3c0ec2:spectra/specs_ms.mgf:scan:1850 |
| legonoxamine F | 551.3088 | CCMSLIB00005723609 | mzspec:GNPSTASK-01f346e93bcb4c469014b16207c56409:spectra/specs_ms.mgf:scan:1240 |
| legonoxamine G | 623.3764 | CCMSLIB00005723625 | mzspec:GNPSTASK-4b53c4f871bb416fb5ed51300d6c8d29:spectra/specs_ms.mgf:scan:567 |
| acyl ferrioxamine 1/ legonoxamine A | 637.3917 | CCMSLIB00005723626 | mzspec:GNPSTASK-4b53c4f871bb416fb5ed51300d6c8d29:spectra/specs_ms.mgf:scan:304 |
| legonoxamine H | 665.3893 | CCMSLIB00005723627 | mzspec:GNPSTASK-4b53c4f871bb416fb5ed51300d6c8d29:spectra/specs_ms.mgf:scan:333 |
| (acyl ferrioxamine 1/legonoxamine A) + Al | 661.3492 | CCMSLIB00005723666 | mzspec:GNPSTASK-c017395e82c64b25a38904c12c5b81ec:spectra/specs_ms.mgf:scan:3084 |
| legonoxamine G +Fe | 676.2871 | CCMSLIB00005723658 | mzspec:GNPSTASK-4b53c4f871bb416fb5ed51300d6c8d29:spectra/specs_ms.mgf:scan:3046 |
| acyl ferrioxamine 2 | 679.4025 | CCMSLIB00005723628 | mzspec:GNPSTASK-4b53c4f871bb416fb5ed51300d6c8d29:spectra/specs_ms.mgf:scan:1538 |
| acyl ferrioxamine 1/legonoxamine A | 690.3027 | CCMSLIB00005723660 | mzspec:GNPSTASK-4b53c4f871bb416fb5ed51300d6c8d29:spectra/specs_ms.mgf:scan:555 |
| legonoxamine A glycoside | 799.4456 | CCMSLIB00005723629 | mzspec:GNPSTASK-f16063a00560422d9abc5930d48f492f:spectra/specs_ms.mgf:scan:568 |
| **DFO-Bs** | | | |
| un-pre[5+5*] | 431.2500 | CCMSLIB00005724363 | mzspec:GNPS:TASK-3c73305ef7ed48e19cb18e50b8b6b2bd-spectra/specs_ms.mgf:scan:2201 |
| pre[5+5*] | 433.2662 | CCMSLIB00005724364 | mzspec:GNPSTASK-16d50fb5de564636964680ad8c037b7c:spectra/specs_ms.mgf:scan:730 |
| pre[5+5*] aldehyde | 447.2450 | CCMSLIB00005724365 | mzspec:GNPS:TASK-3c73305ef7ed48e19cb18e50b8b6b2bd-spectra/specs_ms.mgf:scan:2356 |
| pre[ha+5+5*] | 517.3221 | CCMSLIB00005724366 | mzspec:GNPS:TASK-3c73305ef7ed48e19cb18e50b8b6b2bd-spectra/specs_ms.mgf:scan:2775 |
| abloxime / IC202C | 517.3347 | CCMSLIB00005723632 | mzspec:GNPSTASK-4b53c4f871bb416fb5ed51300d6c8d29:spectra/specs_ms.mgf:scan:1795 |
| proferrioxamine G1t | 519.3495 | CCMSLIB00005723607 | mzspec:GNPSTASK-4b53c4f871bb416fb5ed51300d6c8d29:spectra/specs_ms.mgf:scan:1775 |
| DesuA1 | 545.3658 | CCMSLIB00003739981 | mzspec:GNPSTASK-4b53c4f871bb416fb5ed51300d6c8d29:spectra/specs_ms.mgf:scan:64 |
| DesA1 | 547.3449 | CCMSLIB00003739980 | mzspec:GNPSTASK-4b53c4f871bb416fb5ed51300d6c8d29:spectra/specs_ms.mgf:scan:523 |
| Name | Measured mass [M+H]+ | CCMS Library ID | Metabolomics Resolver ID |
| Des A2 / IC202B | 557.2880 | CCMSLIB00005723654 | mzspec:GNPSTASK-4b53c4f871bb416fb5ed51300d6c8d29:spectra/specs_ms.mgf:scan:2183 |
| desferrioxamine B | 561.3602 | CCMSLIB00000075313 | mzspec:GNPSTASK-4b53c4f871bb416fb5ed51300d6c8d29:spectra/specs_ms.mgf:scan:69 |
| ferrioxamine A1 + Al | 571.3033 | CCMSLIB00005723662 | mzspec:GNPSTASK-4b53c4f871bb416fb5ed51300d6c8d29:spectra/specs_ms.mgf:scan:1746 |

| | | | |
|---|---|---|---|
| desferrioxamine N /desf-05 | 575.3762 | CCMSLIB00003739977 | mzspec:GNPSTASK-4b53c4f871bb416fb5ed51300d6c8d29:spectra/specs_ms.mgf:scan:272 |
| C3 acyl DFO-B | 589.3915 | CCMSLIB00005723634 | mzspec:GNPSTASK-4b53c4f871bb416fb5ed51300d6c8d29:spectra/specs_ms.mgf:scan:1826 |
| ferrioxamine N + Al | 599.3349 | CCMSLIB00005723663 | mzspec:GNPSTASK-4b53c4f871bb416fb5ed51300d6c8d29:spectra/specs_ms.mgf:scan:1772 |
| ferrioxamine A1+ Fe | 600.2565 | CCMSLIB00005723655 | mzspec:GNPSTASK-4b53c4f871bb416fb5ed51300d6c8d29:spectra/specs_ms.mgf:scan:233 |
| C4 acyl DFO-B | 603.4073 | CCMSLIB00005723636 | mzspec:GNPSTASK-4b53c4f871bb416fb5ed51300d6c8d29:spectra/specs_ms.mgf:scan:1836 |
| C5 acyl DFO-B | 617.4233 | CCMSLIB00005724368 | mzspec:GNPS:TASK-3c73305ef7ed48e19cb18e50b8b6b2bd-spectra/specs_ms.mgf:scan:3269 |
| C4 acyl hydroxylated DFO-B | 619.4024 | CCMSLIB00005723638 | zspec:GNPSTASK-4b53c4f871bb416fb5ed51300d6c8d29:spectra/specs_ms.mgf:scan:547 |
| ferrioxamine N + Fe | 628.2883 | CCMSLIB00005723656 | mzspec:GNPSTASK-4b53c4f871bb416fb5ed51300d6c8d29:spectra/specs_ms.mgf:scan:1780 |
| uC6 acyl DFO-B | 629.4238 | CCMSLIB00005724369 | mzspec:GNPS:TASK-3c73305ef7ed48e19cb18e50b8b6b2bd-spectra/specs_ms.mgf:scan:3332 |
| C6 acyl DFO-B | 631.4388 | CCMSLIB00005723639 | mzspec:GNPSTASK-4b53c4f871bb416fb5ed51300d6c8d29:spectra/specs_ms.mgf:scan:593 |
| C5 acyl FO-B + Al | 641.3807 | CCMSLIB00005723673 | mzspec:GNPSTASK-01f346e93bcb4c469014b16207c56409:spectra/specs_ms.mgf:scan:841 |
| C4 hydroxyl acyl FO-B | 643.3599 | CCMSLIB00005723665 | mzspec:GNPSTASK-c017395e82c64b25a38904c12c5b81ec:spectra/specs_ms.mgf:scan:2990 |
| uC7 acyl DFO-B | 643.4389 | CCMSLIB00005724370 | mzspec:GNPS:TASK-3c73305ef7ed48e19cb18e50b8b6b2bd-spectra/specs_ms.mgf:scan:3408 |
| C7 acyl-DFO | 645.4550 | CCMSLIB00000072059 | mzspec:GNPSTASK-4b53c4f871bb416fb5ed51300d6c8d29:spectra/specs_ms.mgf:scan:611 |
| C6 acyl FO-B + Al | 655.3937 | CCMSLIB00005723672 | mzspec:GNPSTASK-4bd8abbc22fe4d439c8ee86ba408ba27:spectra/specs_ms.mgf:scan:110 |
| C8 acyl-DFO | 659.4697 | CCMSLIB00000072060 | mzspec:GNPSTASK-4b53c4f871bb416fb5ed51300d6c8d29:spectra/specs_ms.mgf:scan:1916 |
| C7 acyl FO-B + Al | 669.4106 | CCMSLIB00005723671 | mzspec:GNPSTASK-4bd8abbc22fe4d439c8ee86ba408ba27:spectra/specs_ms.mgf:scan:133 |
| C5 acyl FO-B + Fe | 670.3341 | CCMSLIB00005723670 | mzspec:GNPSTASK-16d50fb5de564636964680ad8c037b7c:spectra/specs_ms.mgf:scan:781 |
| C6 acyl FO-B | 684.3497 | CCMSLIB00005723659 | mzspec:GNPSTASK-4b53c4f871bb416fb5ed51300d6c8d29:spectra/specs_ms.mgf:scan:3075 |
| uC10 acyl DFO-B | 685.4856 | CCMSLIB00005723643 | mzspec:GNPSTASK-4b53c4f871bb416fb5ed51300d6c8d29:spectra/specs_ms.mgf:scan:1937 |
| C7 acyl FO-B | 698.3655 | CCMSLIB00005723661 | mzspec:GNPSTASK-4b53c4f871bb416fb5ed51300d6c8d29:spectra/specs_ms.mgf:scan:3094 |
| uC11 acyl DFO-B | 699.5007 | CCMSLIB00005724371 | mzspec:GNPS:TASK-3c73305ef7ed48e19cb18e50b8b6b2bd-spectra/specs_ms.mgf:scan:3702 |
| C11 acyl-DFO | 701.5174 | CCMSLIB00000072046 | mzspec:GNPSTASK-4b53c4f871bb416fb5ed51300d6c8d29:spectra/specs_ms.mgf:scan:664 |
| C10 acyl FO-B + Al | 711.4573 | CCMSLIB00005723669 | mzspec:GNPSTASK-11c83a4db6cd4a82a30449fe74d5fa48:spectra/specs_ms.mgf:scan:112 |
| C12 acyl-DFO | 715.5322 | CCMSLIB00000072049 | mzspec:GNPSTASK-4b53c4f871bb416fb5ed51300d6c8d29:spectra/specs_ms.mgf:scan:1980 |
| C13 acyl-DFO | 729.5496 | CCMSLIB00000072052 | mzspec:GNPSTASK-4b53c4f871bb416fb5ed51300d6c8d29:spectra/specs_ms.mgf:scan:1998 |
| uC11 acyl FO-B | 723.4613 | CCMSLIB00005723645 | mzspec:GNPSTASK-16d50fb5de564636964680ad8c037b7c:spectra/specs_ms.mgf:scan:427 |
| C11 acyl FO-B + Al | 725.4745 | CCMSLIB00005723668 | mzspec:GNPSTASK-11c83a4db6cd4a82a30449fe74d5fa48:spectra/specs_ms.mgf:scan:126 |
| uC12 acyl FO-B $[M+Al]^+$ | 737.4776 | CCMSLIB00005724372 | mzspec:GNPS:TASK-3c73305ef7ed48e19cb18e50b8b6b2bd-spectra/specs_ms.mgf:scan:3843 |
| Name | Measured mass [M+H]+ | CCMS Library ID | Metabolomics Resolver ID |
| dodecanedioic DFO-B | 745.5086 | CCMSLIB00005723647 | mzspec:GNPSTASK-4b53c4f871bb416fb5ed51300d6c8d29:spectra/specs_ms.mgf:scan:1917 |
| amphiphilic desferrioxamine 12 | 745.5438 | CCMSLIB00005723611 | mzspec:GNPSTASK-4b53c4f871bb416fb5ed51300d6c8d29:spectra/specs_ms.mgf:scan:1932 |
| tridecanedioic DFO-B | 759.5221 | CCMSLIB00005723649 | mzspec:GNPSTASK-4b53c4f871bb416fb5ed51300d6c8d29:spectra/specs_ms.mgf:scan:1930 |

| | | | |
|---|---|---|---|
| amphiphilic desferrioxamine 15 | 759.5597 | CCMSLIB00005723610 | mzspec:GNPSTASK-4b53c4f871bb416fb5ed51300d6c8d29:spectra/specs_ms.mgf:scan:1944 |
| **DFO-Ds** | | | |
| desferrioxamine H | 461.2602 | CCMSLIB00005723630 | mzspec:GNPSTASK-a172e23b485b4fbea34650f2bf3c0ec2:spectra/specs_ms.mgf:scan:561 |
| glutaric desferrioxamine H | 475.2760 | CCMSLIB00005723631 | mzspec:GNPSTASK-a172e23b485b4fbea34650f2bf3c0ec2:spectra/specs_ms.mgf:scan:832 |
| C2 acyl desferrioxamine H | 489.2900 | CCMSLIB00005724373 | mzspec:GNPS:TASK-3c73305ef7ed48e19cb18e50b8b6b2bd-spectra/specs_ms.mgf:scan:2629 |
| desferrioxamine D4 | 559.3453 | CCMSLIB00005724367 | mzspec:GNPS:TASK-3c73305ef7ed48e19cb18e50b8b6b2bd-spectra/specs_ms.mgf:scan:2988 |
| deoxydesferrioxamine D3 | 587.3757 | CCMSLIB00005724374 | mzspec:GNPS:TASK-3c73305ef7ed48e19cb18e50b8b6b2bd-spectra/specs_ms.mgf:scan:3130 |
| Desferrioxamine D3 | 589.3915 | CCMSLIB00005723633 | mzspec:GNPSTASK-4b53c4f871bb416fb5ed51300d6c8d29:spectra/specs_ms.mgf:scan:1474 |
| desferrioxamine D1 | 603.3714 | CCMSLIB00005723635 | mzspec:GNPSTASK-4b53c4f871bb416fb5ed51300d6c8d29:spectra/specs_ms.mgf:scan:294 |
| C2 acyl DFO-D | 617.3864 | CCMSLIB00005724375 | mzspec:GNPS:TASK-3c73305ef7ed48e19cb18e50b8b6b2bd-spectra/specs_ms.mgf:scan:3266 |
| ferrioxamine D1 + Al | 627.3287 | CCMSLIB00005723664 | mzspec:GNPSTASK-4b53c4f871bb416fb5ed51300d6c8d29:spectra/specs_ms.mgf:scan:1449 |
| C4 acyl DFO-D | 645.4185 | CCMSLIB00005723640 | mzspec:GNPSTASK-4b53c4f871bb416fb5ed51300d6c8d29:spectra/specs_ms.mgf:scan:595 |
| ferrioxamine D1 + Fe | 656.2822 | CCMSLIB00005723657 | mzspec:GNPSTASK-4b53c4f871bb416fb5ed51300d6c8d29:spectra/specs_ms.mgf:scan:1455 |
| C5 acyl DFO-D | 659.4276 | CCMSLIB00005723641 | mzspec:GNPSTASK-4b53c4f871bb416fb5ed51300d6c8d29:spectra/specs_ms.mgf:scan:1905 |
| C6 acyl DFO-D | 673.4498 | CCMSLIB00005723642 | mzspec:GNPSTASK-4b53c4f871bb416fb5ed51300d6c8d29:spectra/specs_ms.mgf:scan:629 |
| C7 acyl DFO-D | 687.4655 | CCMSLIB00005723644 | mzspec:GNPSTASK-4b53c4f871bb416fb5ed51300d6c8d29:spectra/specs_ms.mgf:scan:646 |
| C7 acyl FO-D + Al | 711.4199 | CCMSLIB00005723667 | mzspec:GNPSTASK-c017395e82c64b25a38904c12c5b81ec:spectra/specs_ms.mgf:scan:3267 |
| C11 acyl DFO-D | 743.5280 | CCMSLIB00005723646 | mzspec:GNPSTASK-4b53c4f871bb416fb5ed51300d6c8d29:spectra/specs_ms.mgf:scan:2013 |
| C12 acyl DFO-D | 757.5436 | CCMSLIB00005723648 | mzspec:GNPSTASK-4b53c4f871bb416fb5ed51300d6c8d29:spectra/specs_ms.mgf:scan:2031 |
| C13 acyl DFO-D | 771.5607 | CCMSLIB00005723650 | mzspec:GNPSTASK-4b53c4f871bb416fb5ed51300d6c8d29:spectra/specs_ms.mgf:scan:740 |
| C14 acyl DFO-D | 785.5749 | CCMSLIB00005723651 | mzspec:GNPSTASK-4b53c4f871bb416fb5ed51300d6c8d29:spectra/specs_ms.mgf:scan:750 |
| **tenacibactins** | | | |
| tenacibactin C | 503.3070 | CCMSLIB00005724342 | mzspec:GNPS:TASK-3c73305ef7ed48e19cb18e50b8b6b2bd-spectra/specs_ms.mgf:scan:2705 |
| tenacibactin E | 531.3380 | CCMSLIB00005724344 | mzspec:GNPS:TASK-3c73305ef7ed48e19cb18e50b8b6b2bd-spectra/specs_ms.mgf:scan:2861 |
| tenacibactin F | 545.3545 | CCMSLIB00005724343 | mzspec:GNPS:TASK-3c73305ef7ed48e19cb18e50b8b6b2bd-spectra/specs_ms.mgf:scan:2925 |
| tenacibactin G | 587.3990 | CCMSLIB00005724345 | mzspec:GNPS:TASK-3c73305ef7ed48e19cb18e50b8b6b2bd-spectra/specs_ms.mgf:scan:3131 |
| tenacibactin H | 601.4168 | CCMSLIB00005724346 | mzspec:GNPS:TASK-3c73305ef7ed48e19cb18e50b8b6b2bd-spectra/specs_ms.mgf:scan:3192 |
| tenacibactin I | 615.4326 | CCMSLIB00005724347 | mzspec:GNPS:TASK-3c73305ef7ed48e19cb18e50b8b6b2bd-spectra/specs_ms.mgf:scan:3255 |
| tenacibactin J | 629.4484 | CCMSLIB00005724348 | mzspec:GNPS:TASK-3c73305ef7ed48e19cb18e50b8b6b2bd-spectra/specs_ms.mgf:scan:3334 |

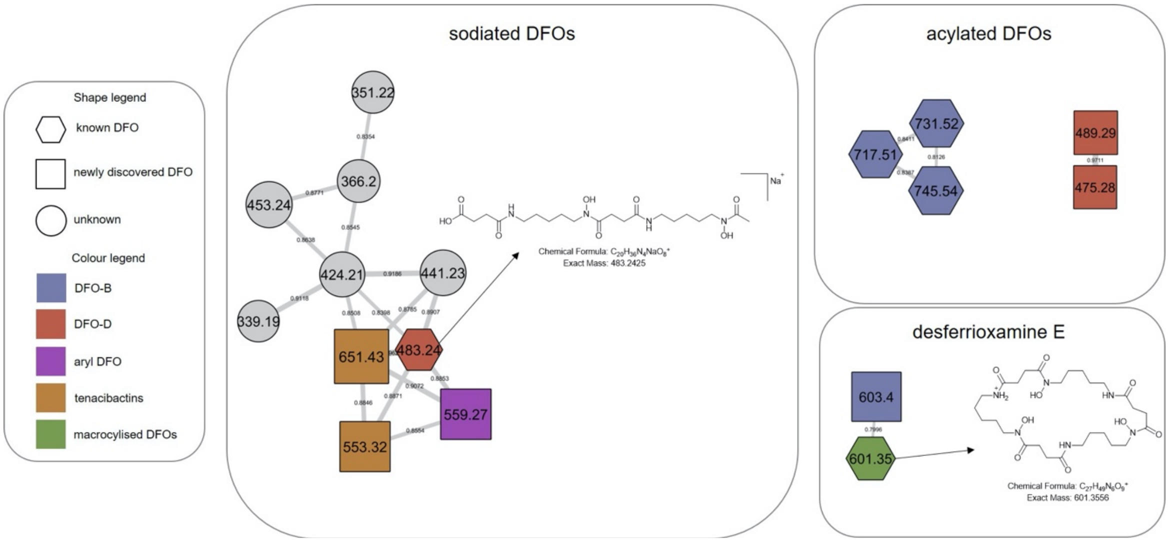


Figure S1. Three additional subnetworks containing distinct clustering. The sodiated cluster represents some $Na^+$ adducts observed in the MS data. The acylated DFOs are a mixture of acylated unsaturated derivatives and some small desferrioxamine H derivatives. The last group shows the placement of desferrioxamine E, one of the macrocyclic DFOs proven to be produced by *Streptomyces* spp. All of the identities of the nodes annotated can be found in Table S1.

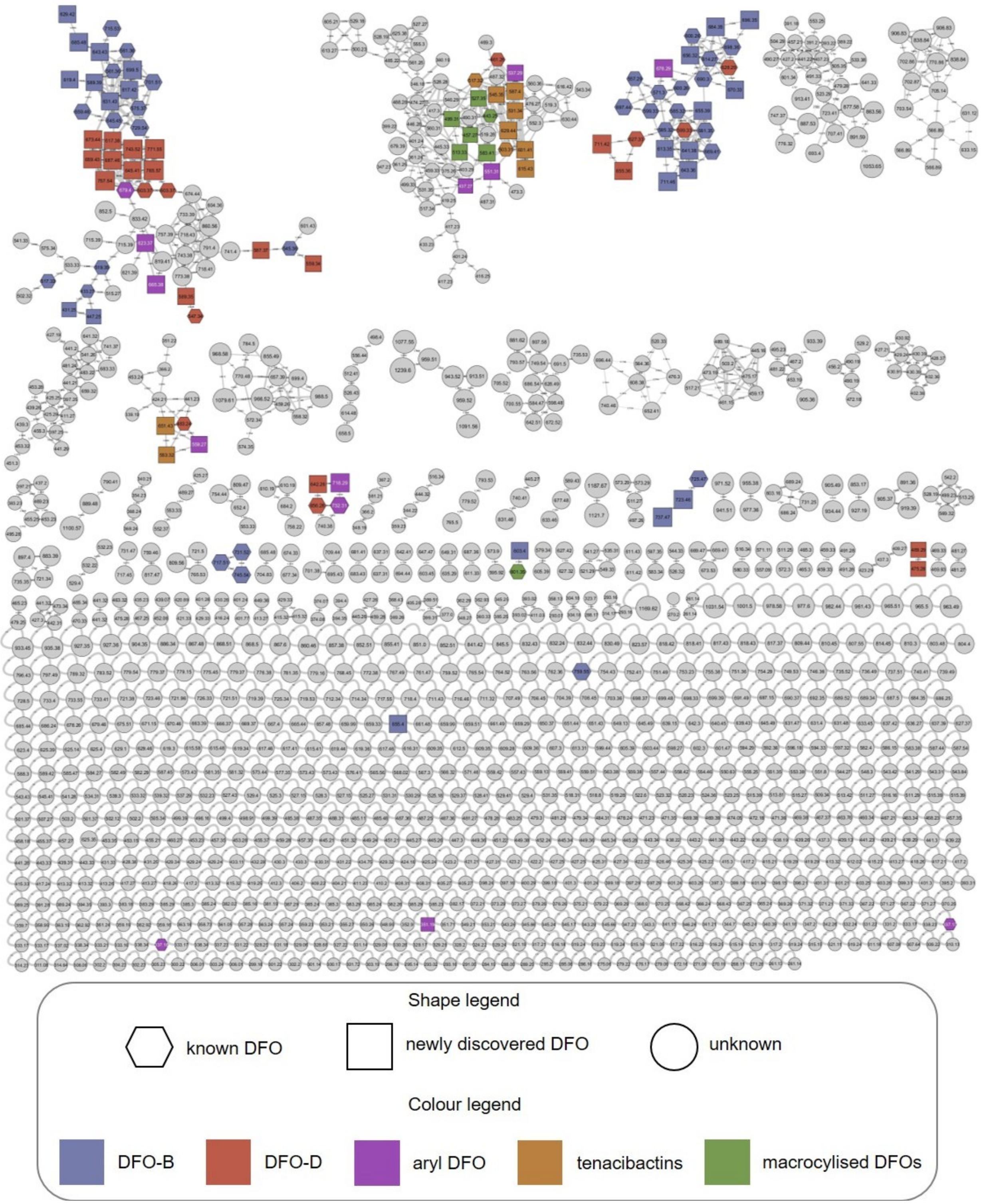


Figure S2. Molecular network annotated with different structural DFO families as represented in the legend.

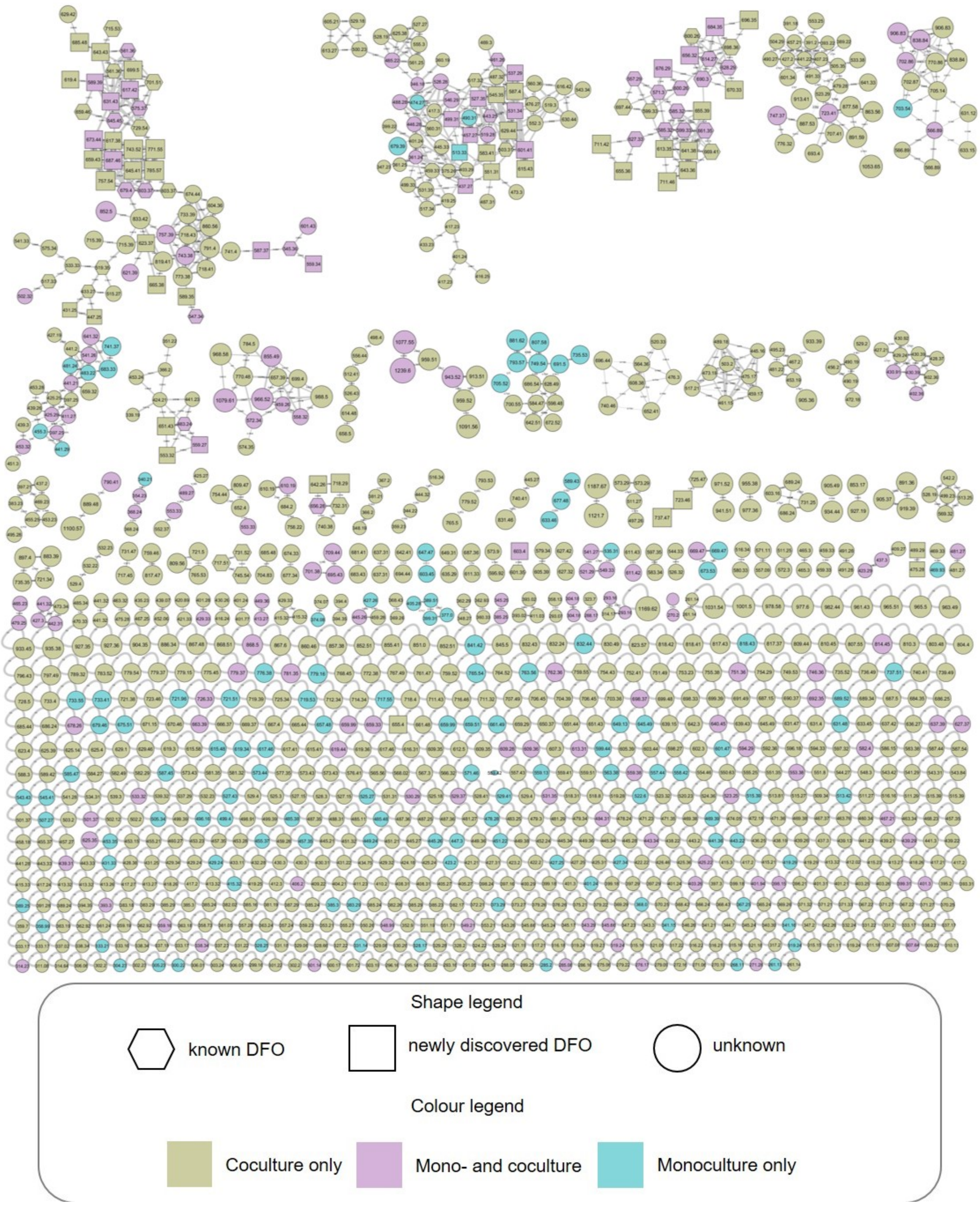


Figure S3. Molecular network annotated with the origin of each node and whether they derive from *Streptomyces* sp. S29 monoculture, both *Streptomyces* sp. S29 and the fungal cocultivation, or only the cocultivation samples.

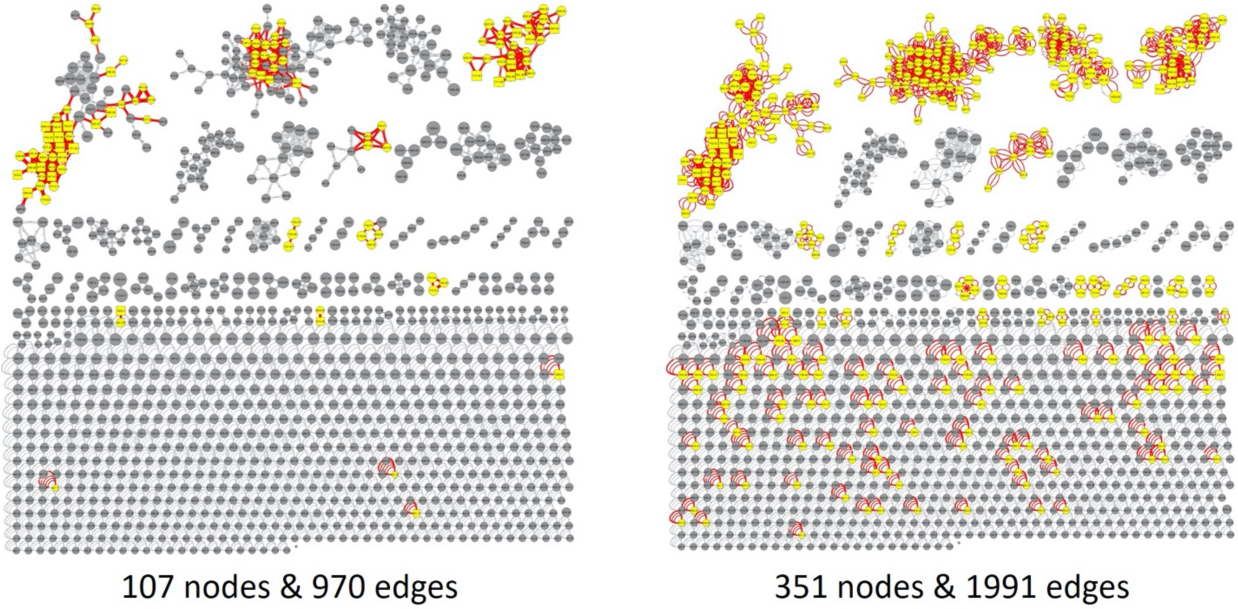


Figure S4. Merged GNPS molecular network and MS2LDA. (Left) Highlighted nodes and edges representing desferrioxamines identified from the original molecular network generated in Figure 2. (Right) Highlighted nodes and edges representing nodes and edges that contain manually curated desferrioxamine motifs via MS2LDA.

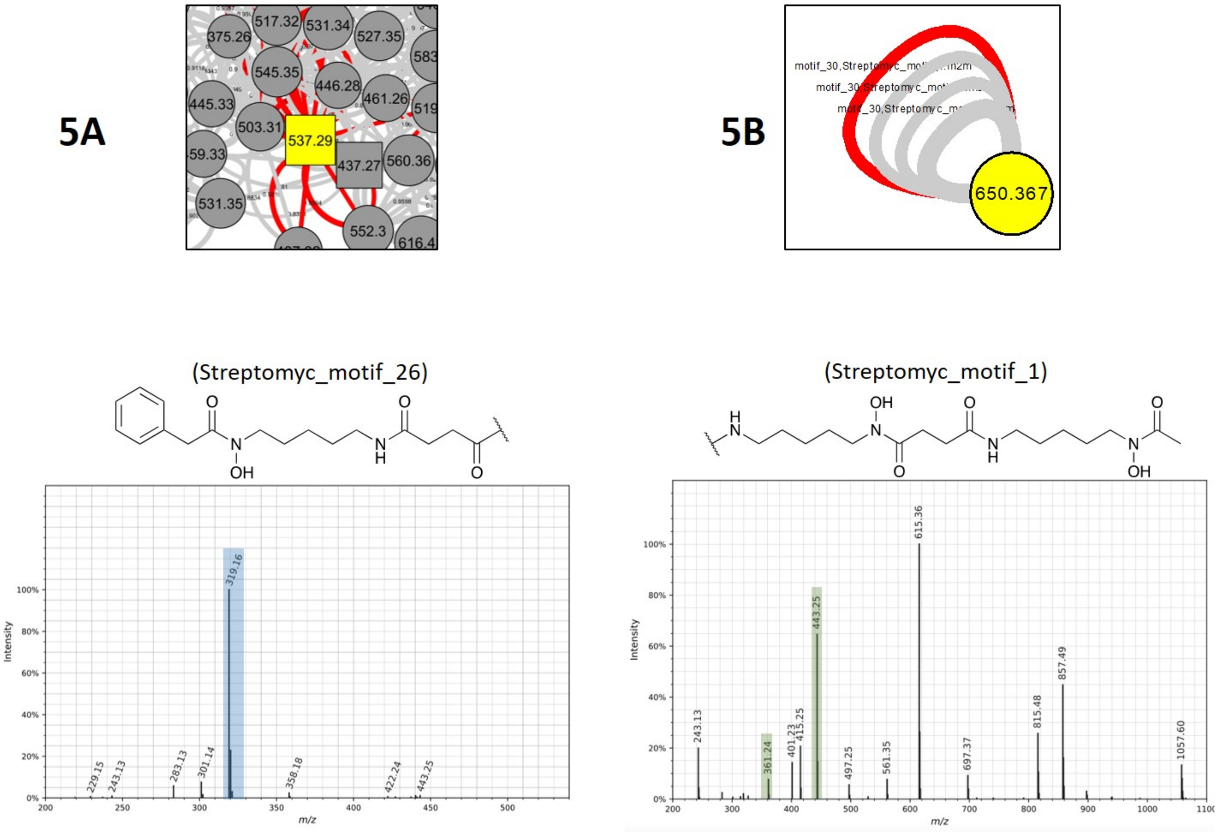


Figure S5. Merged GNPS molecular network and MS2LDA validations. (5A) Node representing legonoxamine E (Figure S13), the manually curated desferrioxamine motif identified in MS2LDA and the MS/MS fragmentation of this metabolite with the motif highlighted. (5B) Node representing an unknown desferrioxamine, the manually curated desferrioxamine motif identified in MS2LDA and the MS/MS fragmentation of this metabolite with the motif highlighted. Motifs can be found in MotifDB at http://ms2lda.org/motifdb/motif_set/32/.

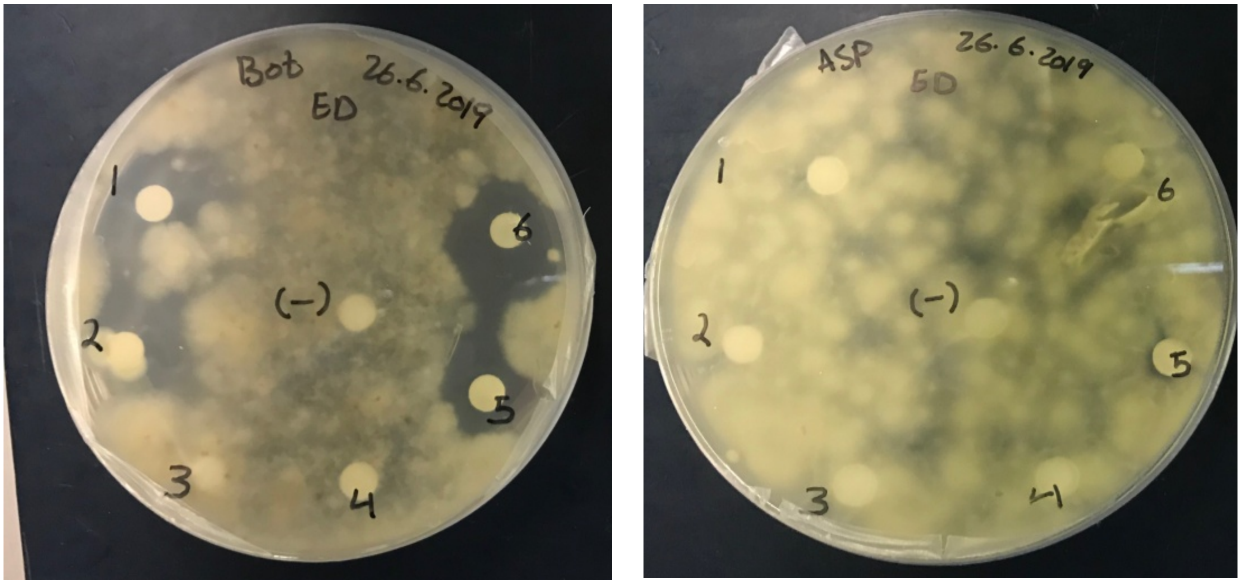


| Sample Number | Fraction ID |
|---|---|
| 1 | S29 + *A. niger* DCM |
| 2 | S29 + *A. niger* Methanol/Water |
| 3 | S29 + *A. niger* sec-butanol 25% MeOH SPE |
| 4 | S29 + *A. niger* sec-butanol 50% MeOH SPE |
| 5 | S29 + *A. niger* sec-butanol 100% MeOH SPE |
| 6 | S29 + *A. niger* n-hexane |

Figure S6. Disc diffusion assay of *Streptomyces* sp. S29 + *Aspergillus niger* coculture extracts tested against *Botrytis cinerea* (left) and *Aspergillus niger* (right). It is clear that the *sec*-butanol fractions, largely containing desferrioxamines described, show no biological activity. Sample number 5 in the *Botrytis cinerea* plate does show some inhibition of the fungi but due to previous studies showing desferrioxamines contain no antifungal activity, it was presumed to be resulting from another metabolite in the extract.

### macrocylised DFOs

Figure S7. MS/MS for reported compound: ***m/z 457.2660***

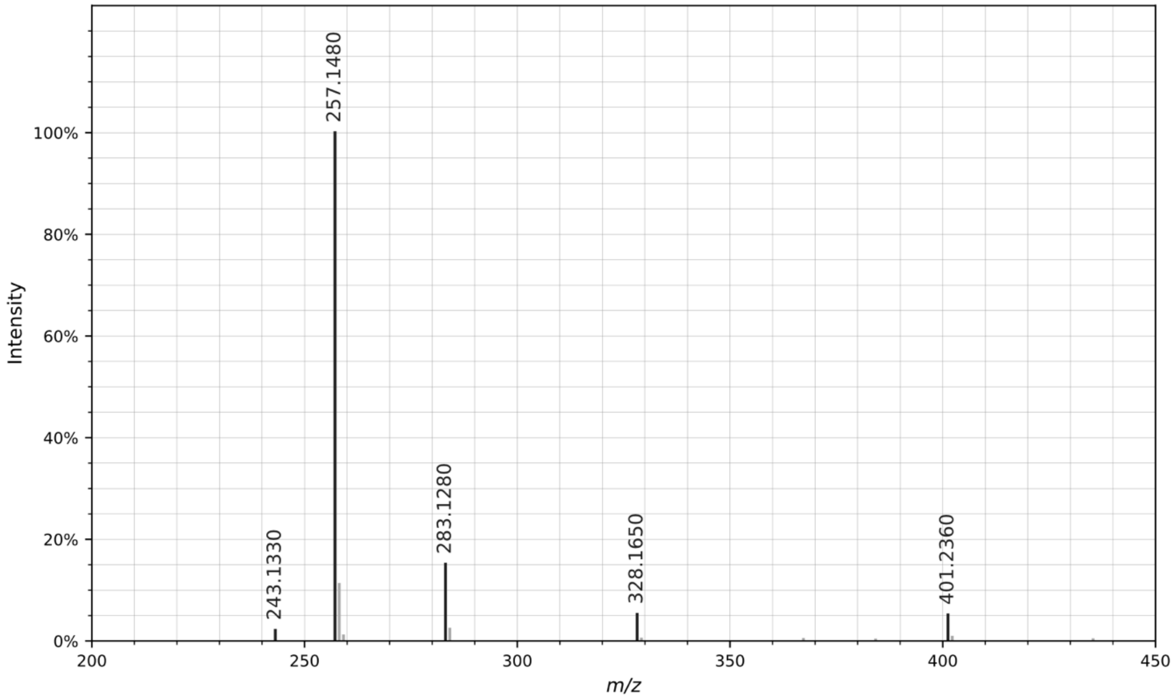


CID MS/MS spectrum of the $[M+H]^+$ ion of *m/z* 457.2600. All fragment ions in the annotated structure match ions in the spectrum. Other fragment ions present in the spectrum are products of non-sequential fragmentation and are present in all DFOs.

Figure S8. Annotation of MS/MS for reported compounds: ***m/z 499.3110***

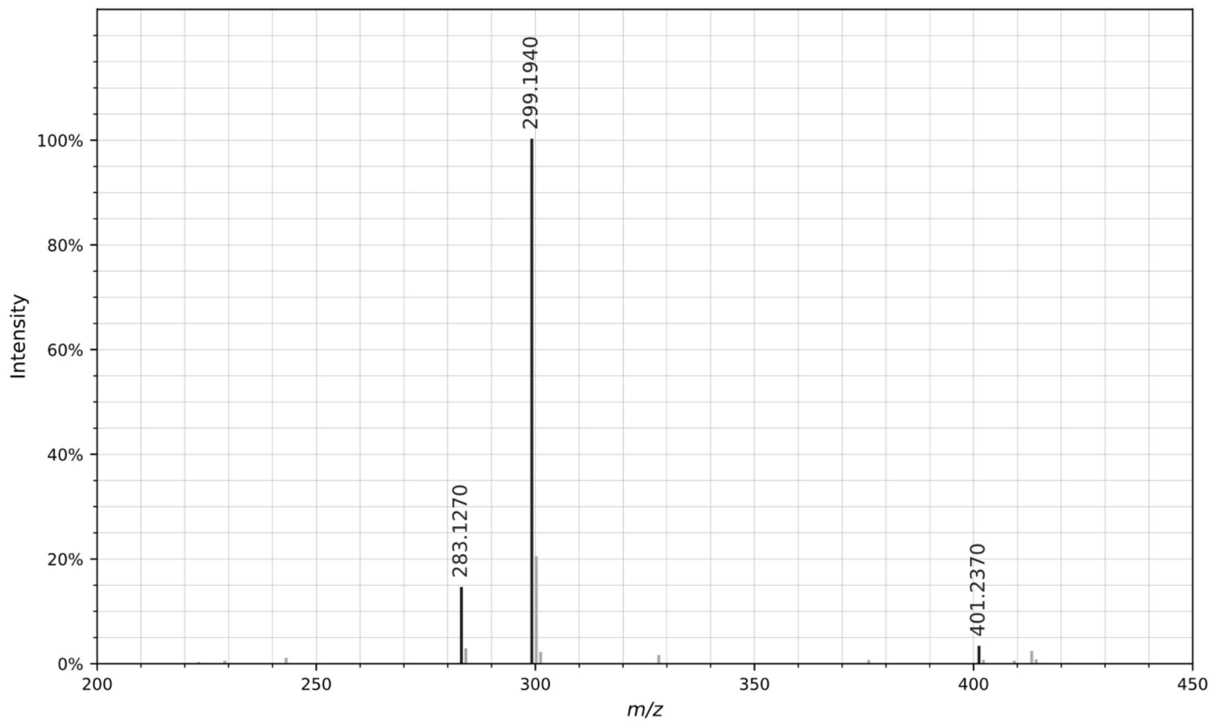


CID MS/MS spectrum of the $[M+H]^+$ ion of *m/z* 499.3110. All fragment ions in the annotated structure match ions in the spectrum. Other fragment ions present in the spectrum are products of non-sequential fragmentation and are present in all DFOs.

Figure S9. Annotation of MS/MS for reported compounds: ***m/z* 513.3310**

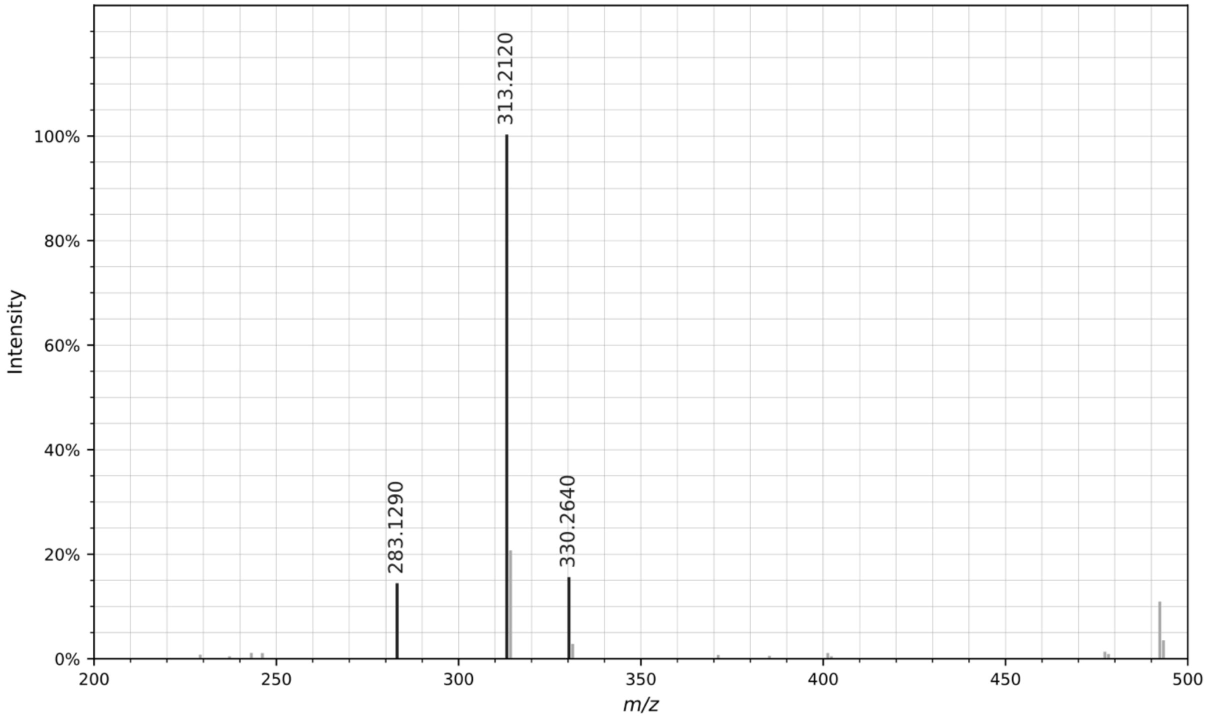


CID MS/MS spectrum of the $[M+H]^+$ ion of *m/z* 513.3310. All fragment ions in the annotated structure match ions in the spectrum. Other fragment ions present in the spectrum are products of non-sequential fragmentation and are present in all DFOs.

Figure S10. Annotation of MS/MS for reported compounds: ***m/z 527.3450***

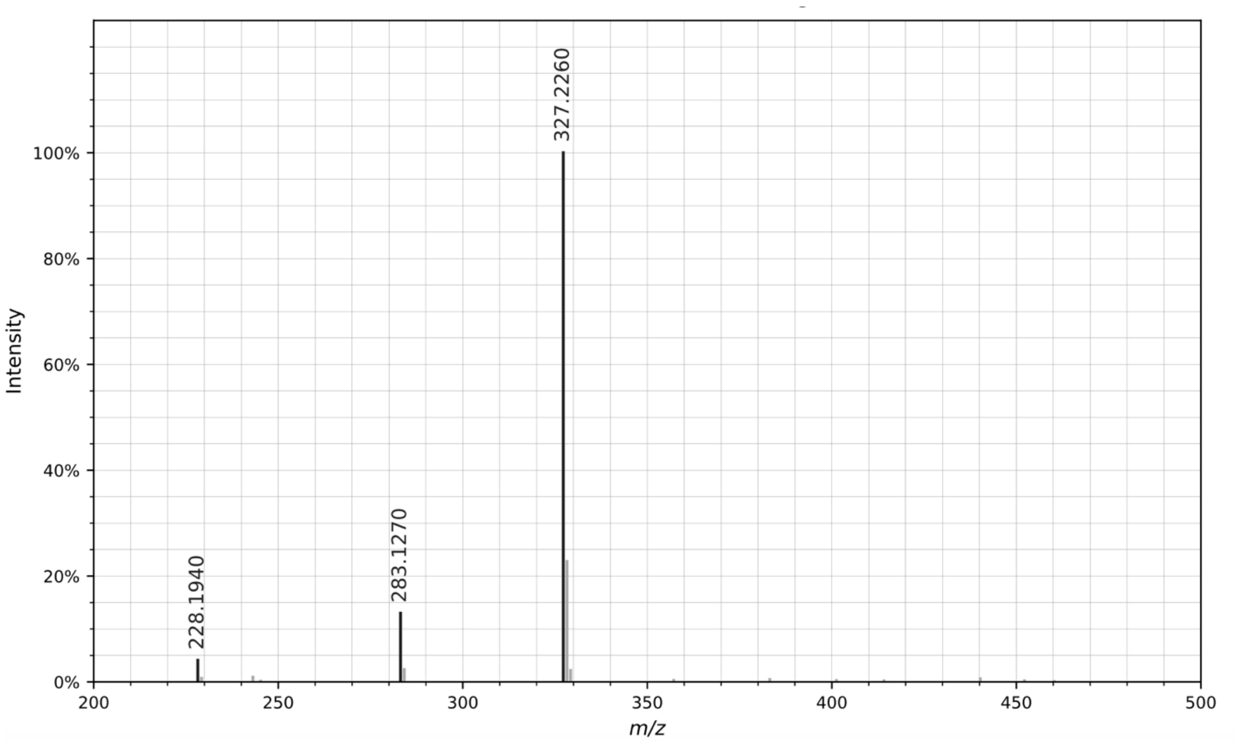


CID MS/MS spectrum of the $[M+H]^+$ ion of *m/z* 527.3450. All fragment ions in the annotated structure match ions in the spectrum. Other fragment ions present in the spectrum are products of non-sequential fragmentation and are present in all DFOs.

Figure S11. Annotation of MS/MS for reported compounds: ***m/z 583.4050***

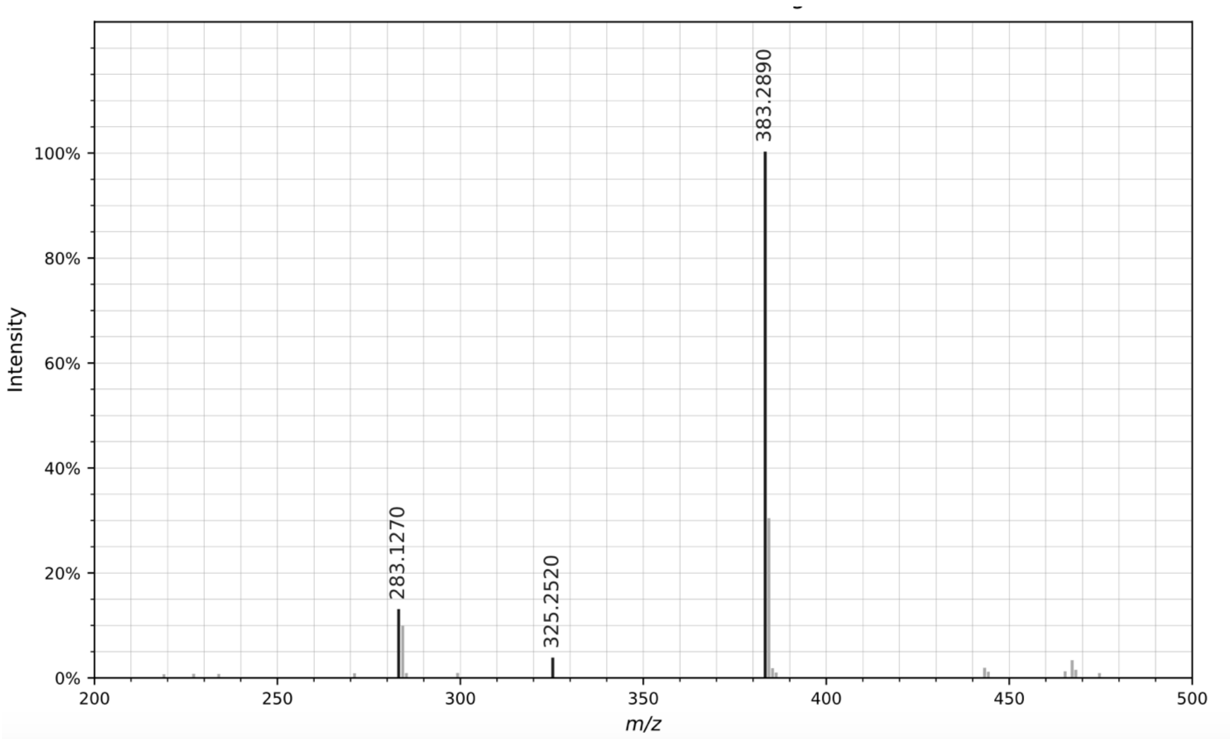


CID MS/MS spectrum of the $[M+H]^+$ ion of *m/z* 583.4050. All fragment ions in the annotated structure match ions in the spectrum. Other fragment ions present in the spectrum are products of non-sequential fragmentation and are present in all DFOs.

### aryl desferrioxamines

Figure S12. Annotation of MS/MS for reported compounds: **legonoxamine C**

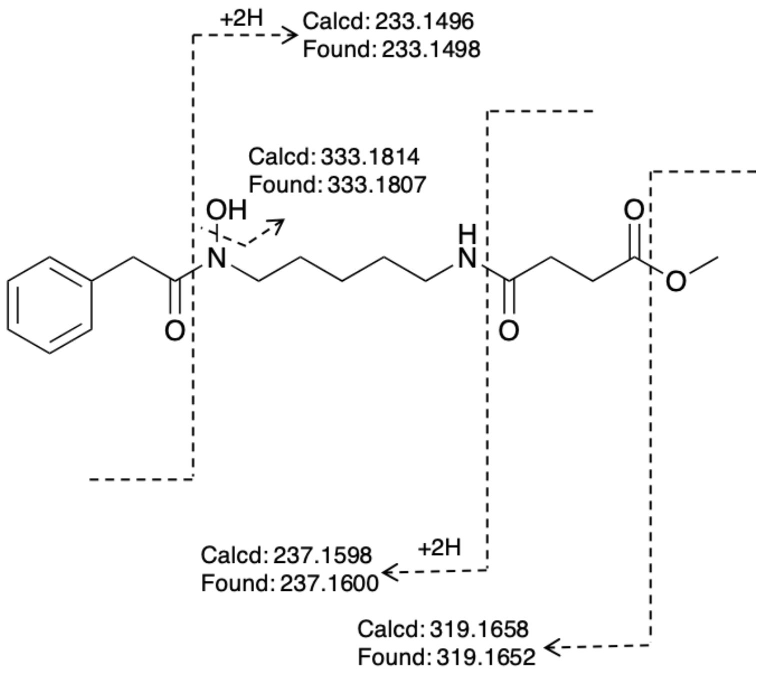


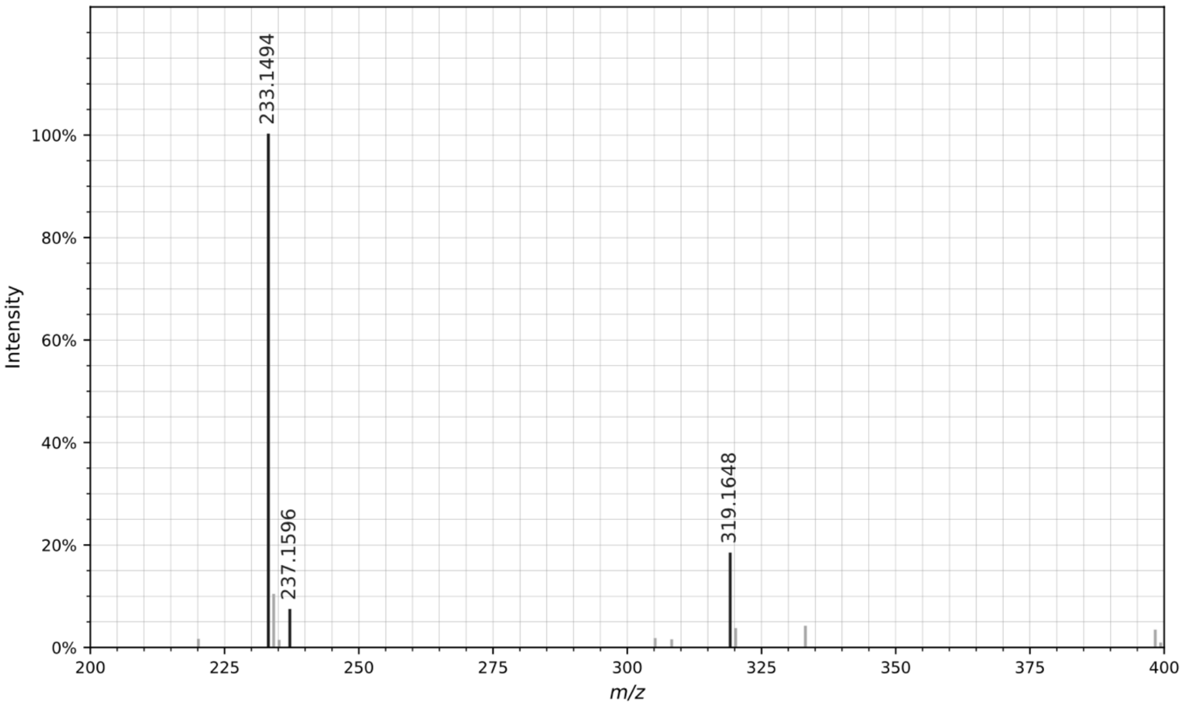


CID MS/MS spectrum of the $[M+H]^+$ ion of legonoxamine C. All fragment ions in the annotated structure match ions in the spectrum. Other fragment ions present in the spectrum are products of non-sequential fragmentation and are present in all DFOs.

Figure S13. Annotation of MS/MS for reported compounds: **legonoxamine D**

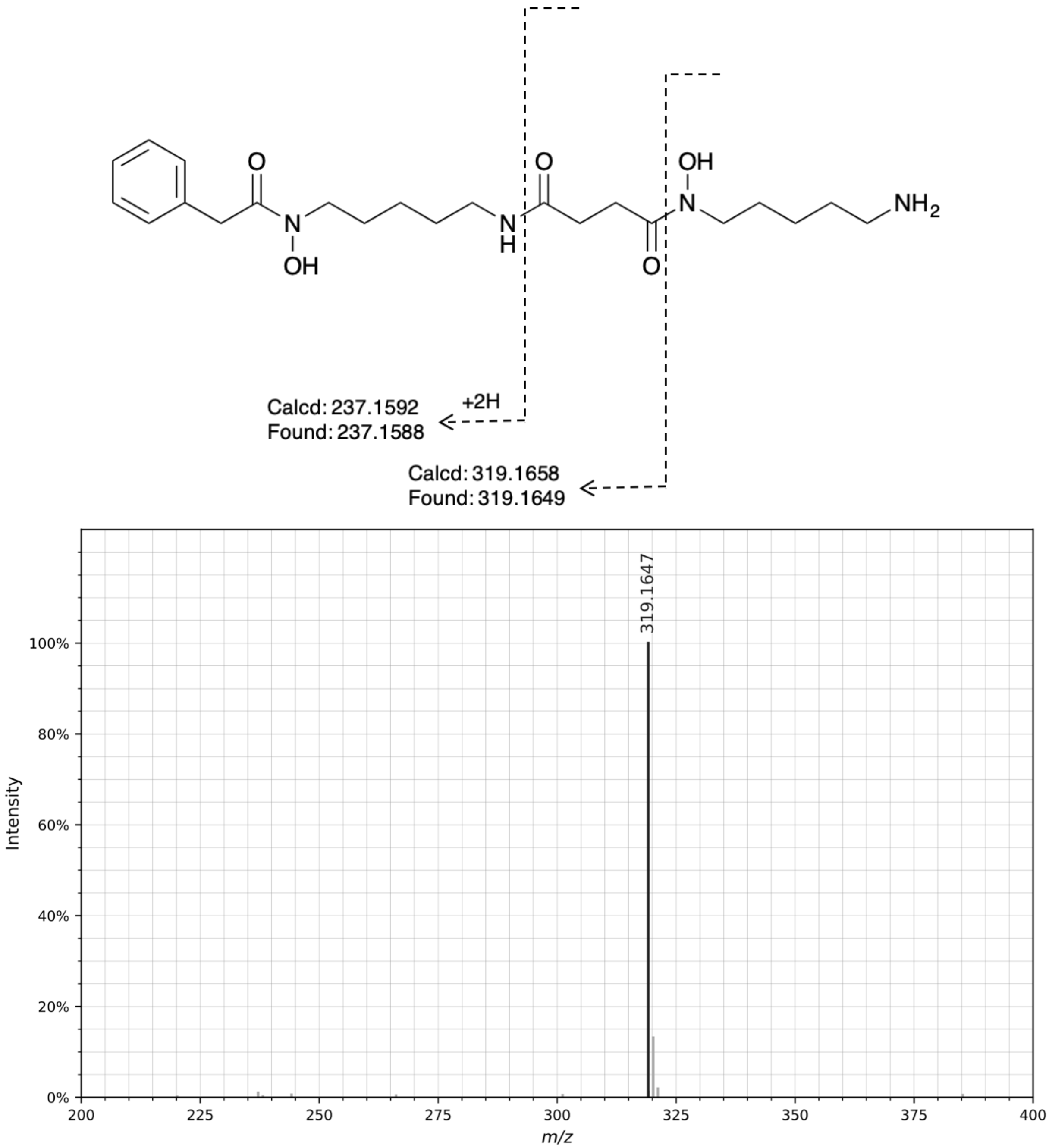


CID MS/MS spectrum of the $[M+H]^+$ ion of legonoxamine D. All fragment ions in the annotated structure match ions in the spectrum. Other fragment ions present in the spectrum are products of non-sequential fragmentation and are present in all DFOs.

Figure S14. Annotation of MS/MS for reported compounds: **legonoxamine E**

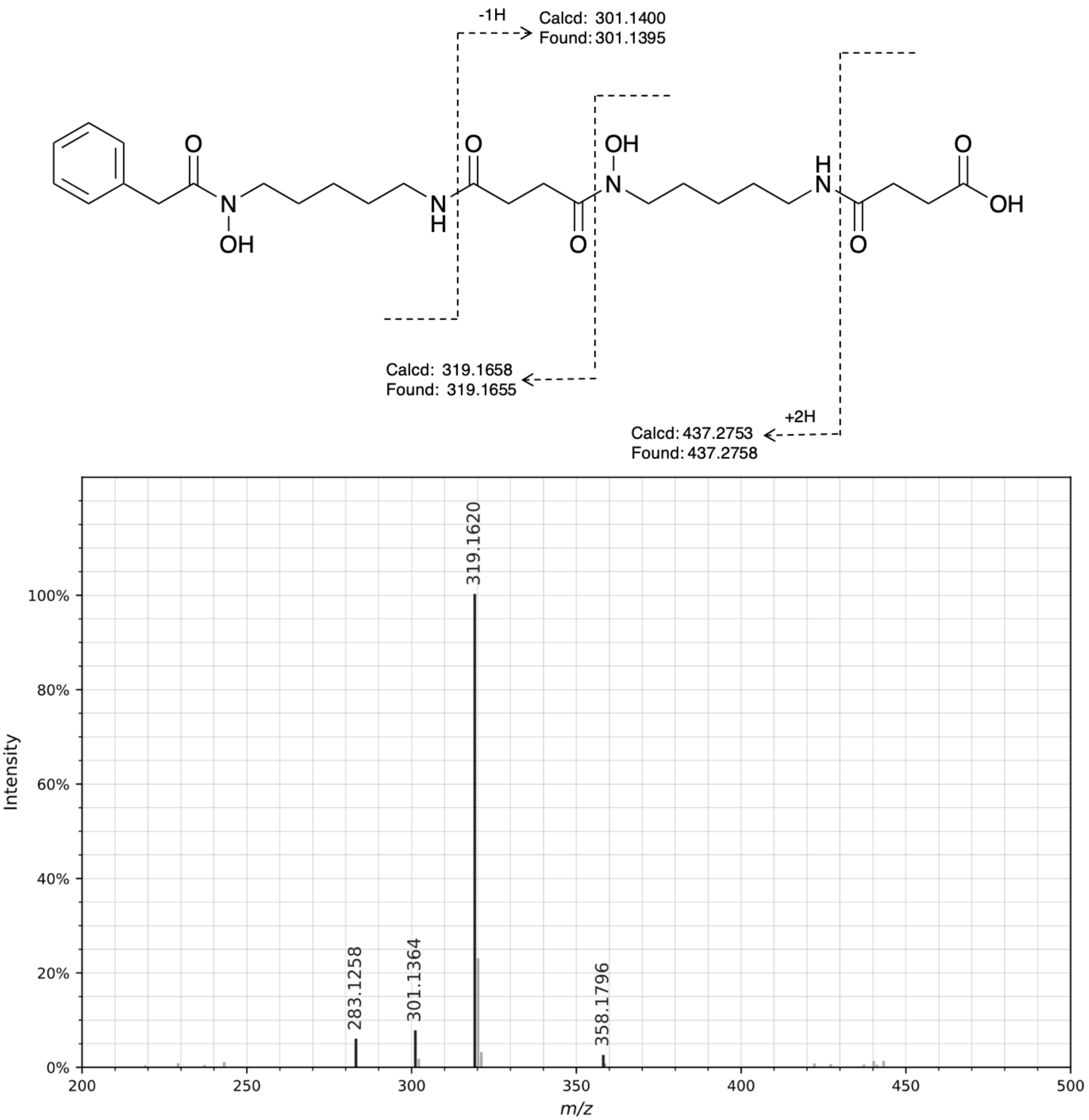


CID MS/MS spectrum of the $[M+H]^+$ ion of legonoxamine E. All fragment ions in the annotated structure match ions in the spectrum. Other fragment ions present in the spectrum are products of non-sequential fragmentation and are present in all DFOs.

Figure S15. Annotation of MS/MS for reported compounds: **legonoxamine F**

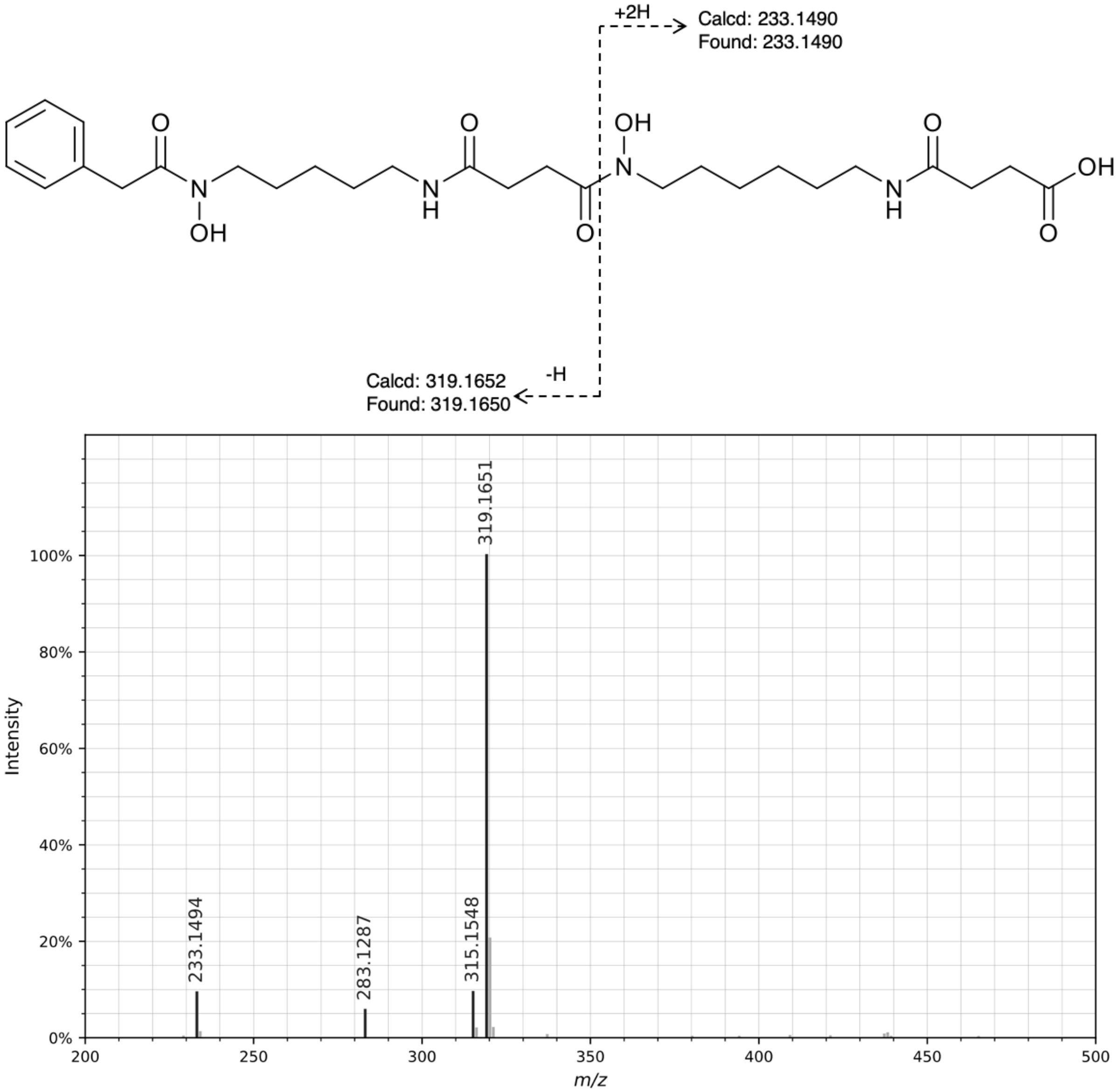


CID MS/MS spectrum of the $[M+H]^+$ ion of legonoxamine F. All fragment ions in the annotated structure match ions in the spectrum. Other fragment ions present in the spectrum are products of non-sequential fragmentation and are present in all DFOs.

Figure S16.   Annotation of MS/MS for reported compounds: **legonoxamine G**

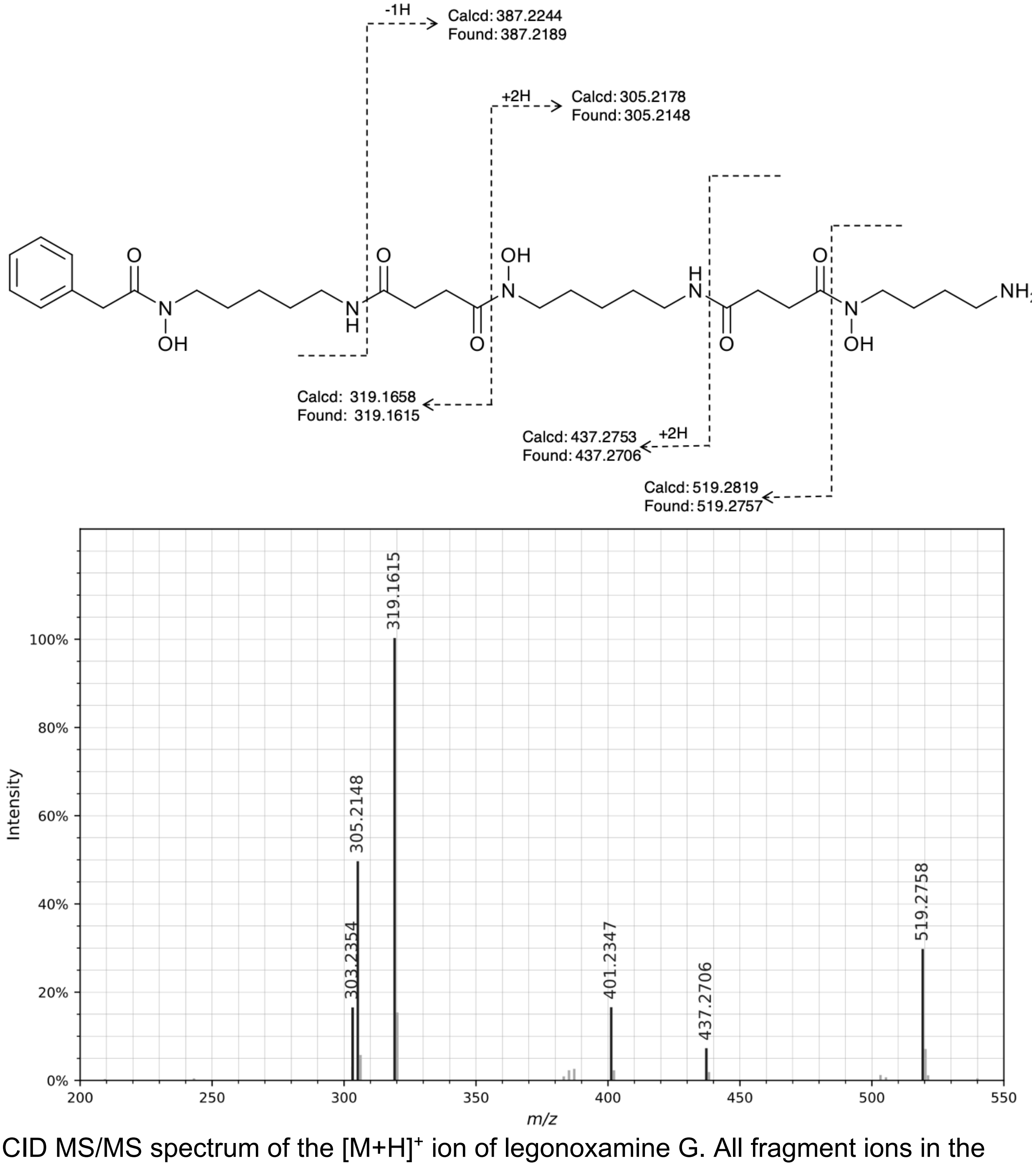


CID MS/MS spectrum of the $[M+H]^+$ ion of legonoxamine G. All fragment ions in the annotated structure match ions in the spectrum. Other fragment ions present in the spectrum are products of non-sequential fragmentation and are present in all DFOs.

Figure S17. Annotation of MS/MS for reported compounds: **legonoxamine H**

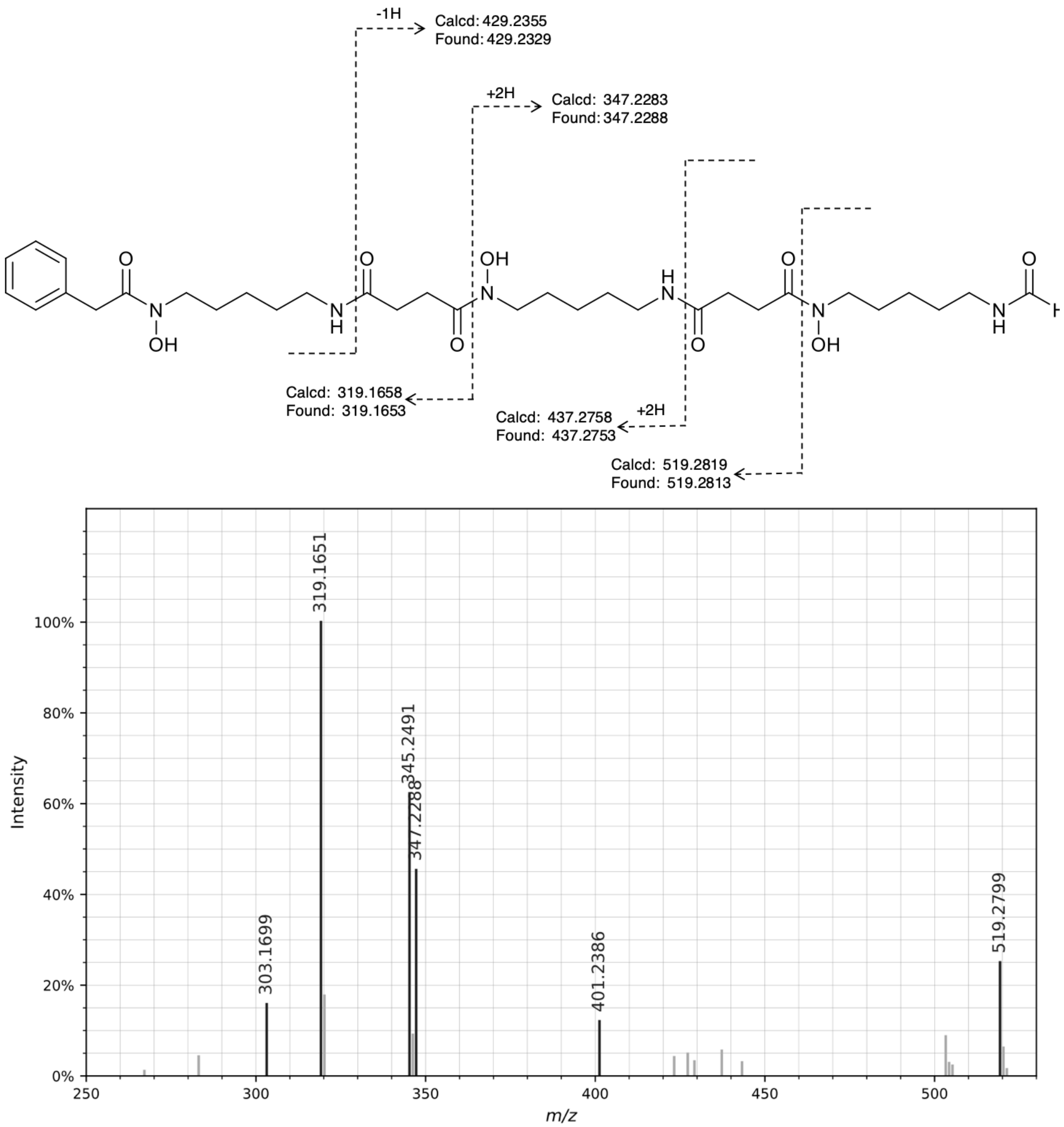


CID MS/MS spectrum of the $[M+H]^+$ ion of legonoxamine H. All fragment ions in the annotated structure match ions in the spectrum. Other fragment ions present in the spectrum are products of non-sequential fragmentation and are present in all DFOs.

Figure S18. Annotation of MS/MS for reported compounds: **legonoxamine A glycoside**

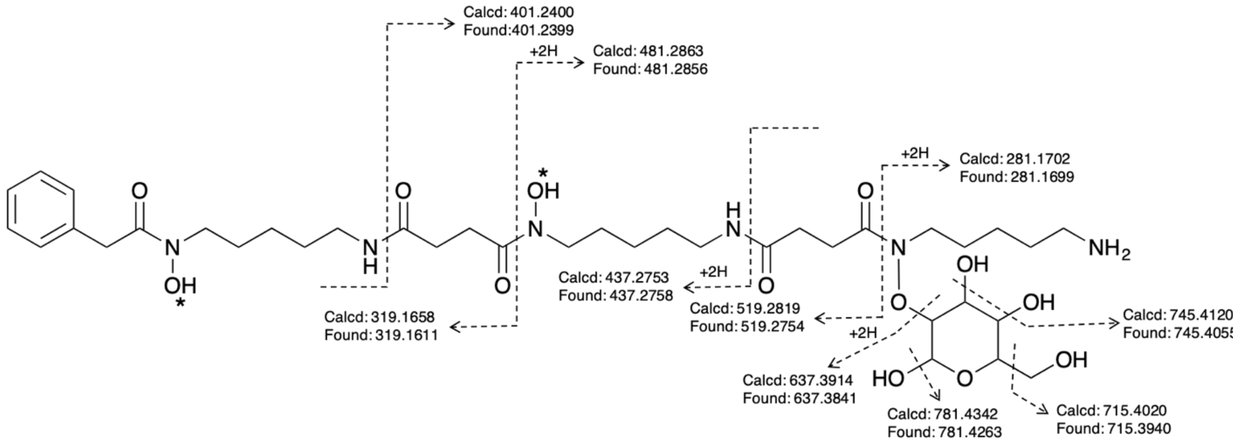


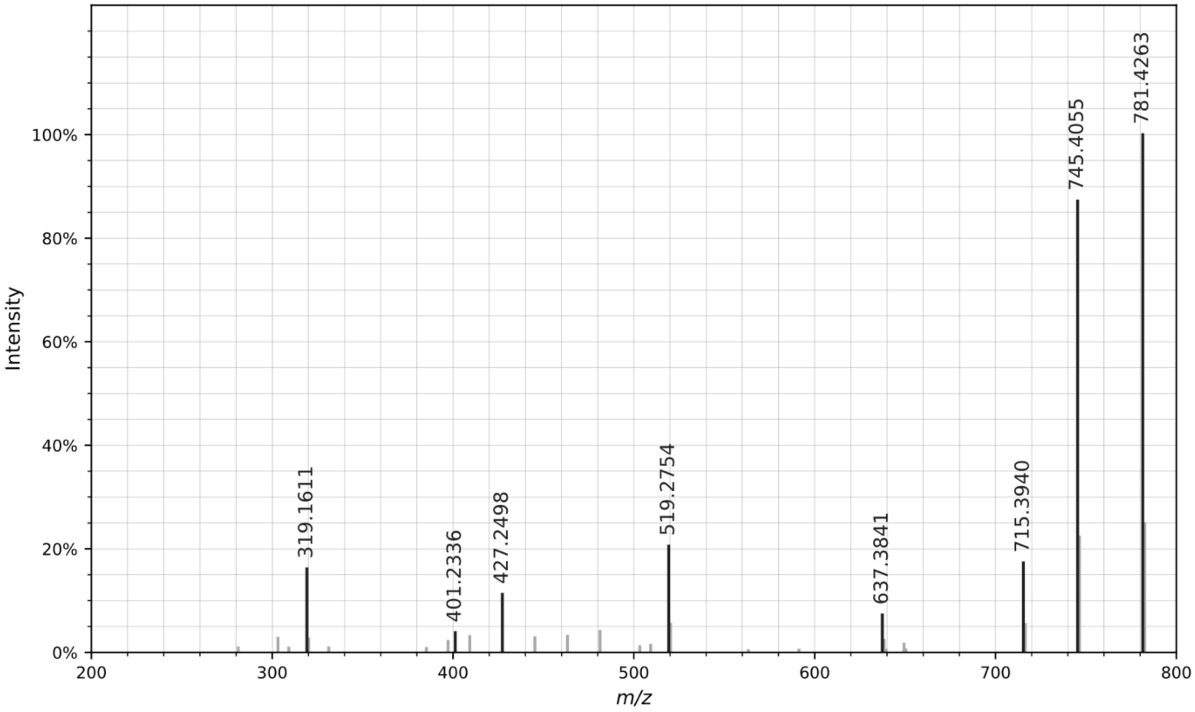


CID MS/MS spectrum of the $[M+H]^+$ ion of legonoxamine A glycoside. All fragment ions in the annotated structure match ions in the spectrum. Other fragment ions present in the spectrum are products of non-sequential fragmentation and are present in all DFOs.

### DFO-Bs

Figure S19. Annotation of MS/MS for reported compounds: **un-pre[5+5*]**

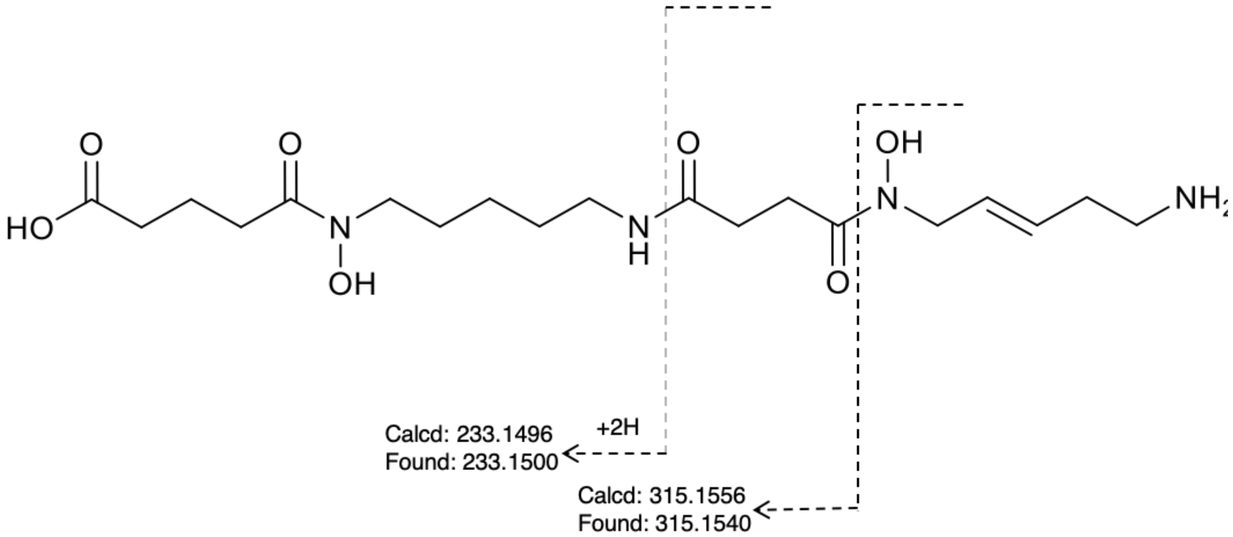


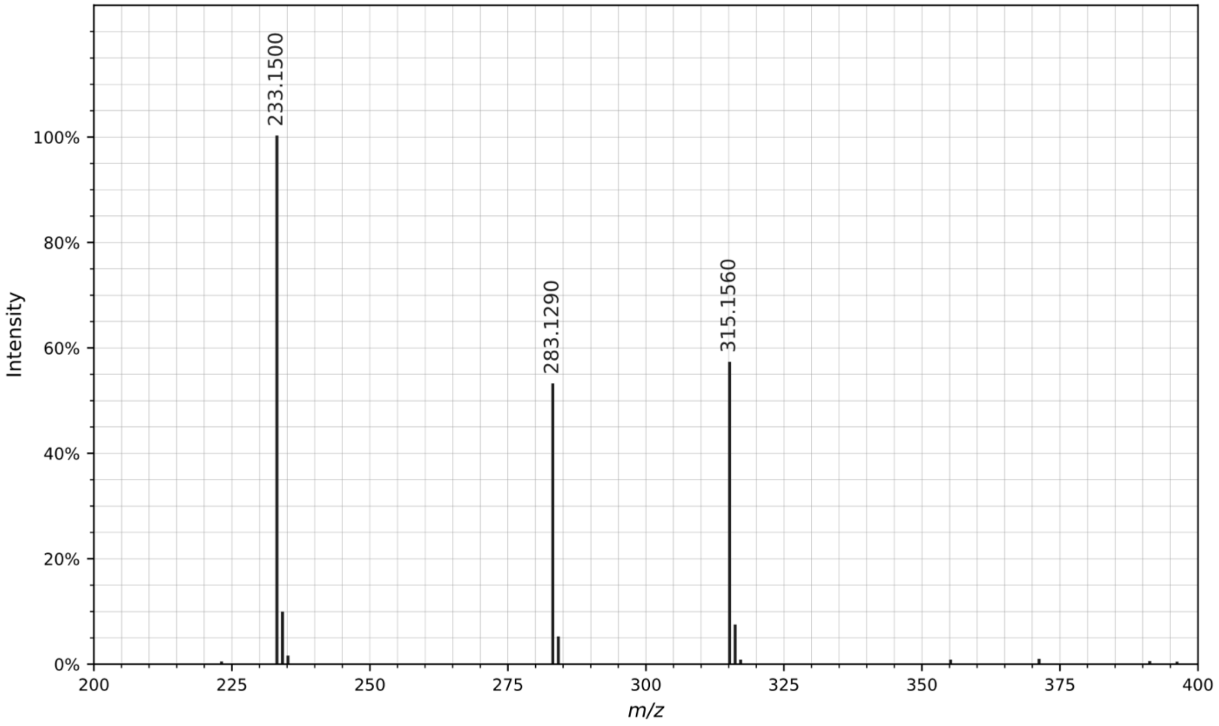


CID MS/MS spectrum of the $[M+H]^+$ ion of un-pre[5+5*]. All fragment ions in the annotated structure match ions in the spectrum. Other fragment ions present in the spectrum are products of non-sequential fragmentation and are present in all DFOs.

Figure S20. Annotation of MS/MS for reported compounds: **pre[5+5*] aldehyde**

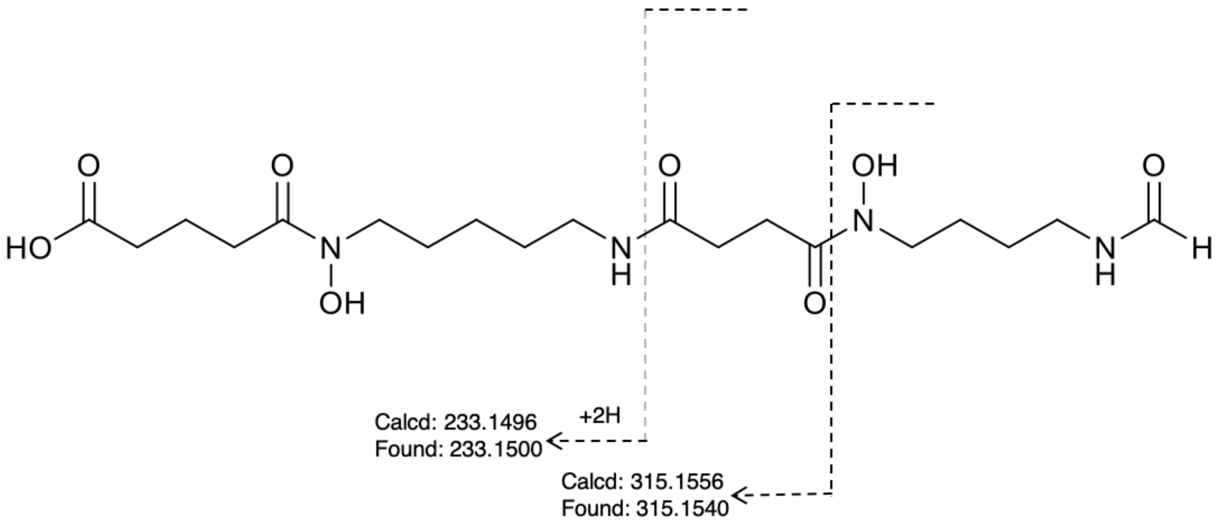


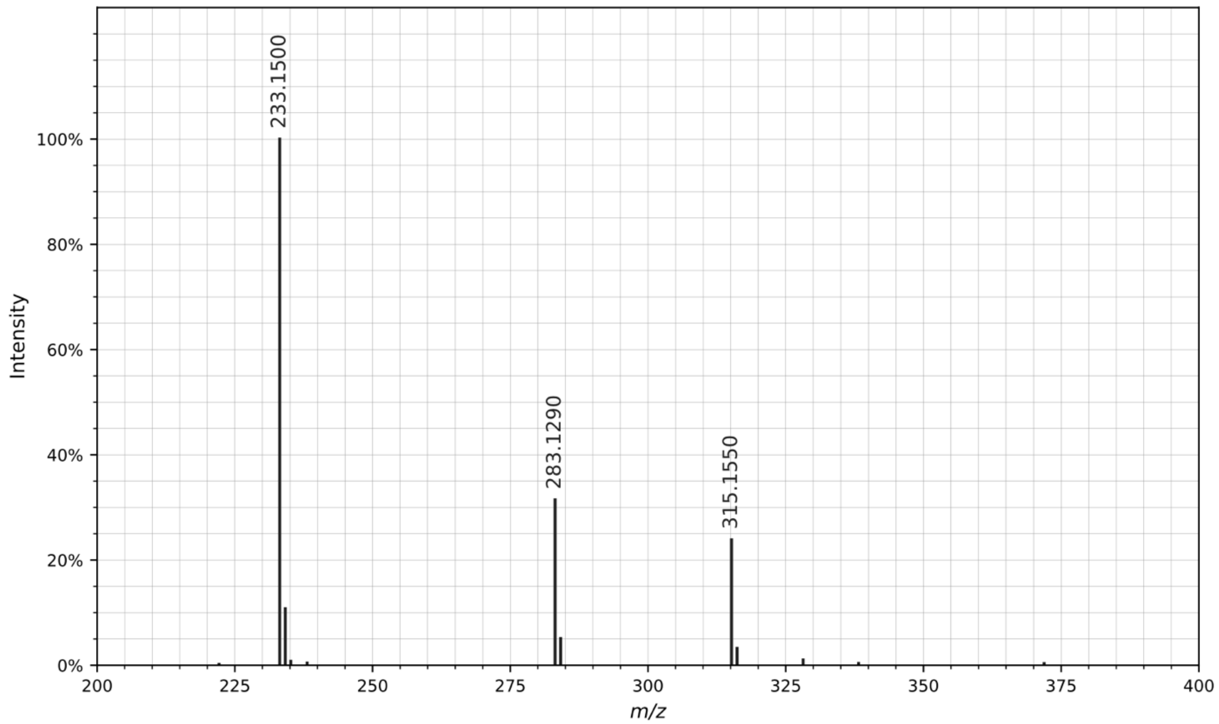


CID MS/MS spectrum of the $[M+H]^+$ ion of pre[5+5*] aldehyde. All fragment ions in the annotated structure match ions in the spectrum. Other fragment ions present in the spectrum are products of non-sequential fragmentation and are present in all DFOs.

Figure S21. Annotation of MS/MS for reported compounds: **desferrioxamine D4**

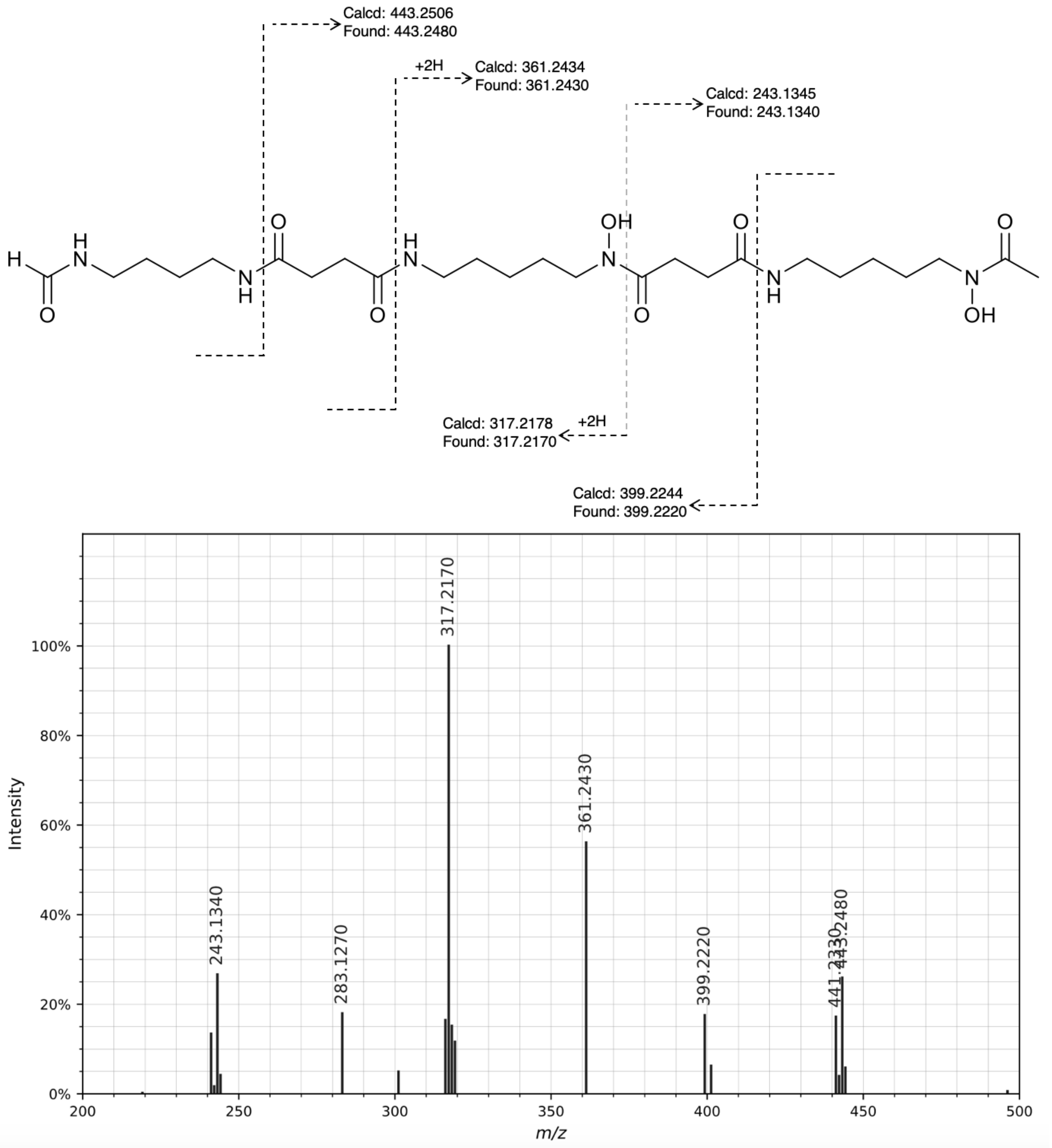


CID MS/MS spectrum of the $[M+H]^+$ ion of desferrioxamine D4. All fragment ions in the annotated structure match ions in the spectrum. Other fragment ions present in the spectrum are products of non-sequential fragmentation and are present in all DFOs.

Figure S22. Annotation of MS/MS for reported compounds: **deoxydesferrioxamine D3**

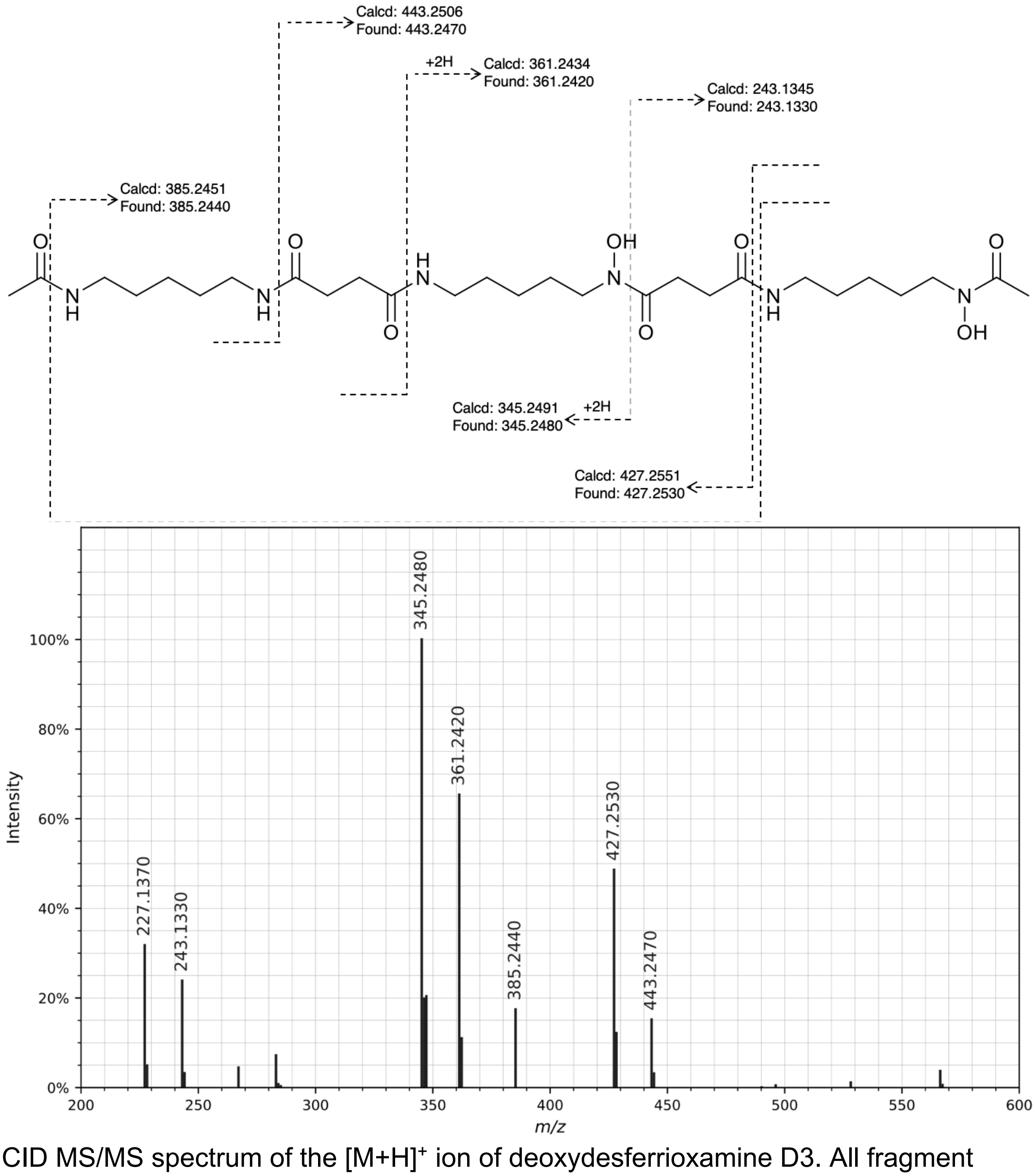


CID MS/MS spectrum of the $[M+H]^+$ ion of deoxydesferrioxamine D3. All fragment ions in the annotated structure match ions in the spectrum. Other fragment ions present in the spectrum are products of non-sequential fragmentation and are present in all DFOs.

Figure S23. Annotation of MS/MS for reported compounds: **C3 acyl DFO-B**

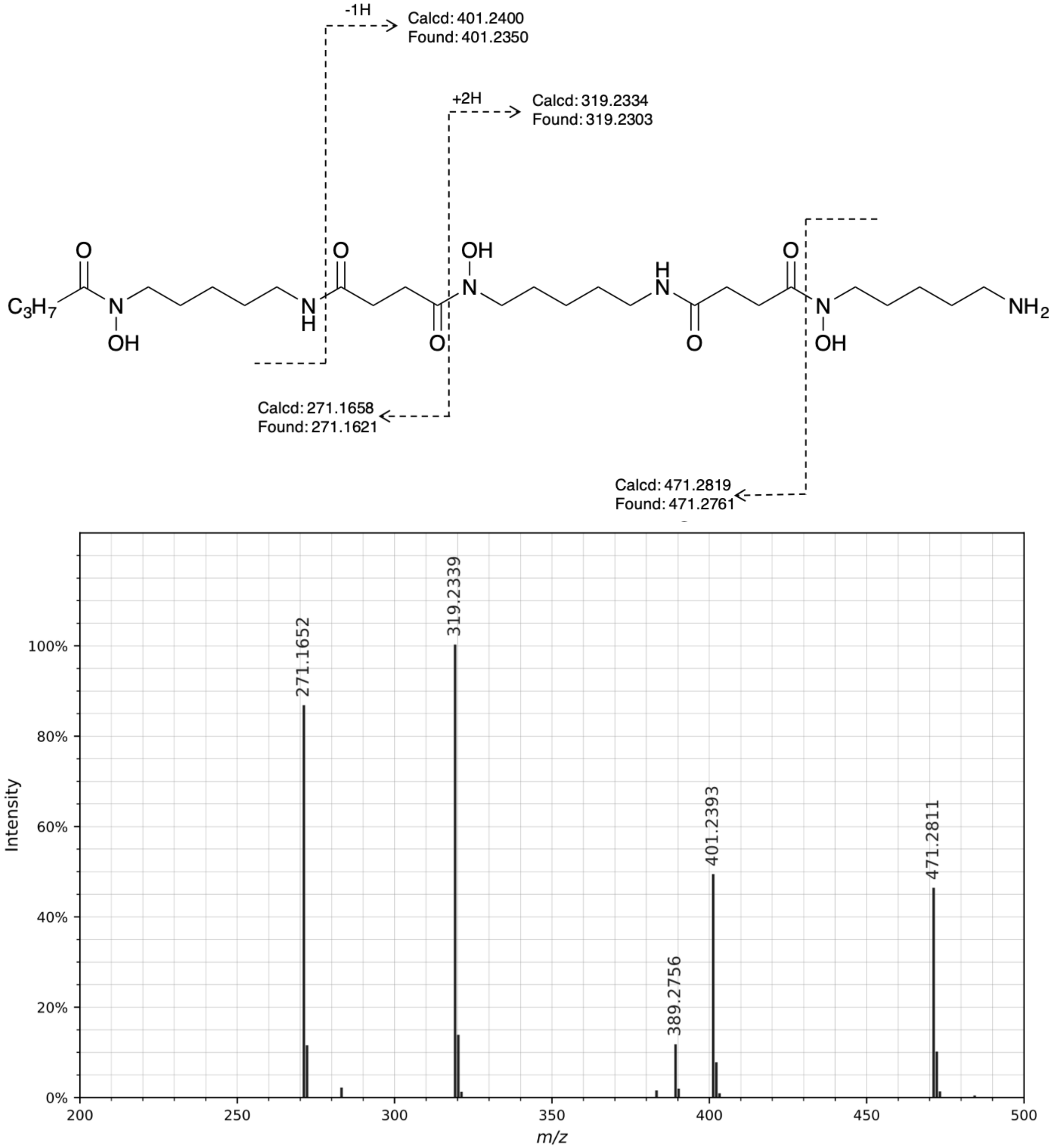


CID MS/MS spectrum of the $[M+H]^+$ ion of C3 acyl DFO-B. All fragment ions in the annotated structure match ions in the spectrum. Other fragment ions present in the spectrum are products of non-sequential fragmentation and are present in all DFOs.

Figure S24. Annotation of MS/MS for reported compounds: **C4 acyl DFO-B**

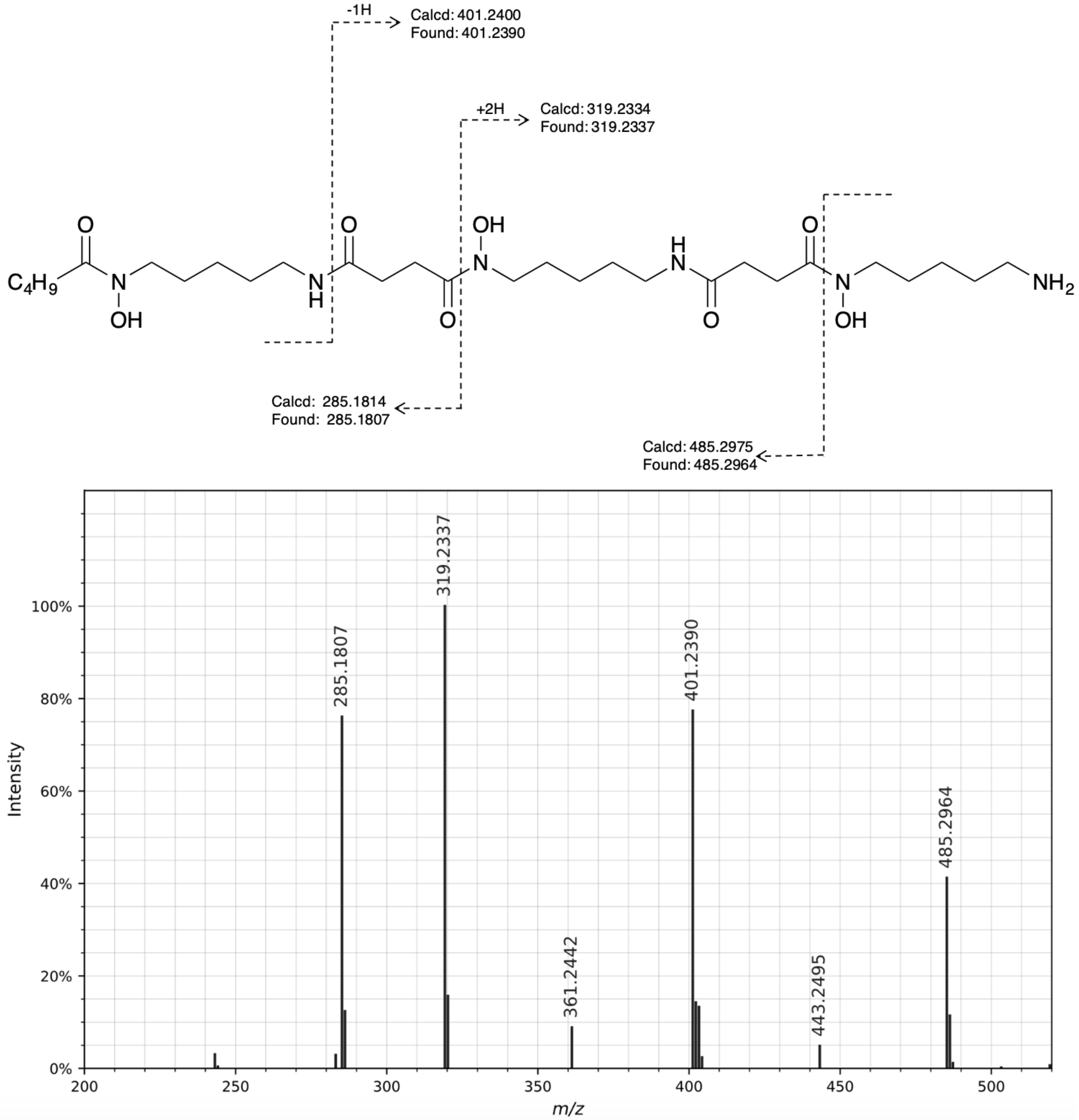


CID MS/MS spectrum of the $[M+H]^+$ ion of C4 acyl DFO-B. All fragment ions in the annotated structure match ions in the spectrum. Other fragment ions present in the spectrum are products of non-sequential fragmentation and are present in all DFOs.

Figure S25. Annotation of MS/MS for reported compounds: **C5 acyl DFO-B**

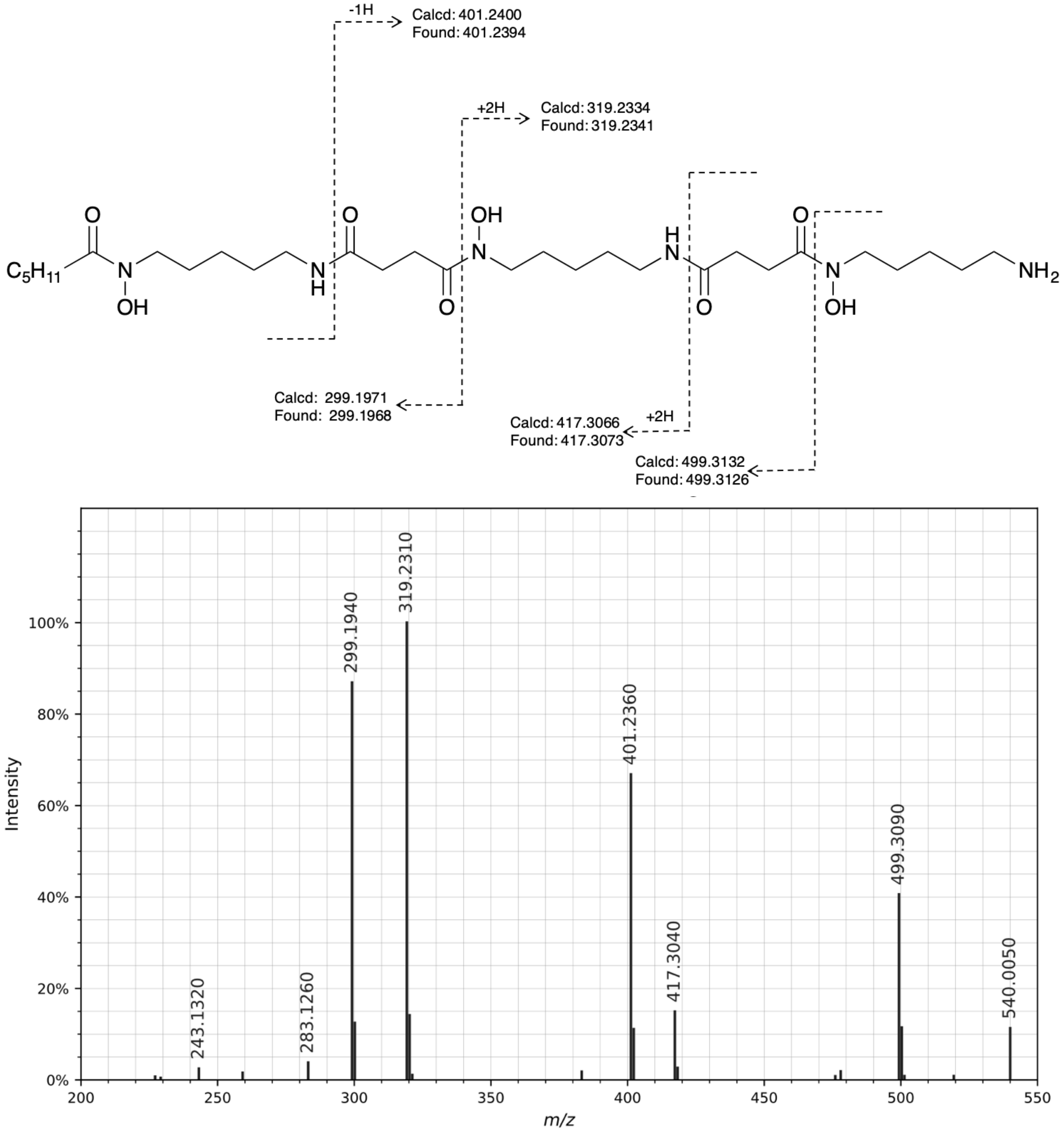


CID MS/MS spectrum of the $[M+H]^+$ ion of C5 acyl DFO-B. All fragment ions in the annotated structure match ions in the spectrum. Other fragment ions present in the spectrum are products of non-sequential fragmentation and are present in all DFOs.

Figure S26. Annotation of MS/MS for reported compounds: **C4 acyl hydroxylated DFO-B**

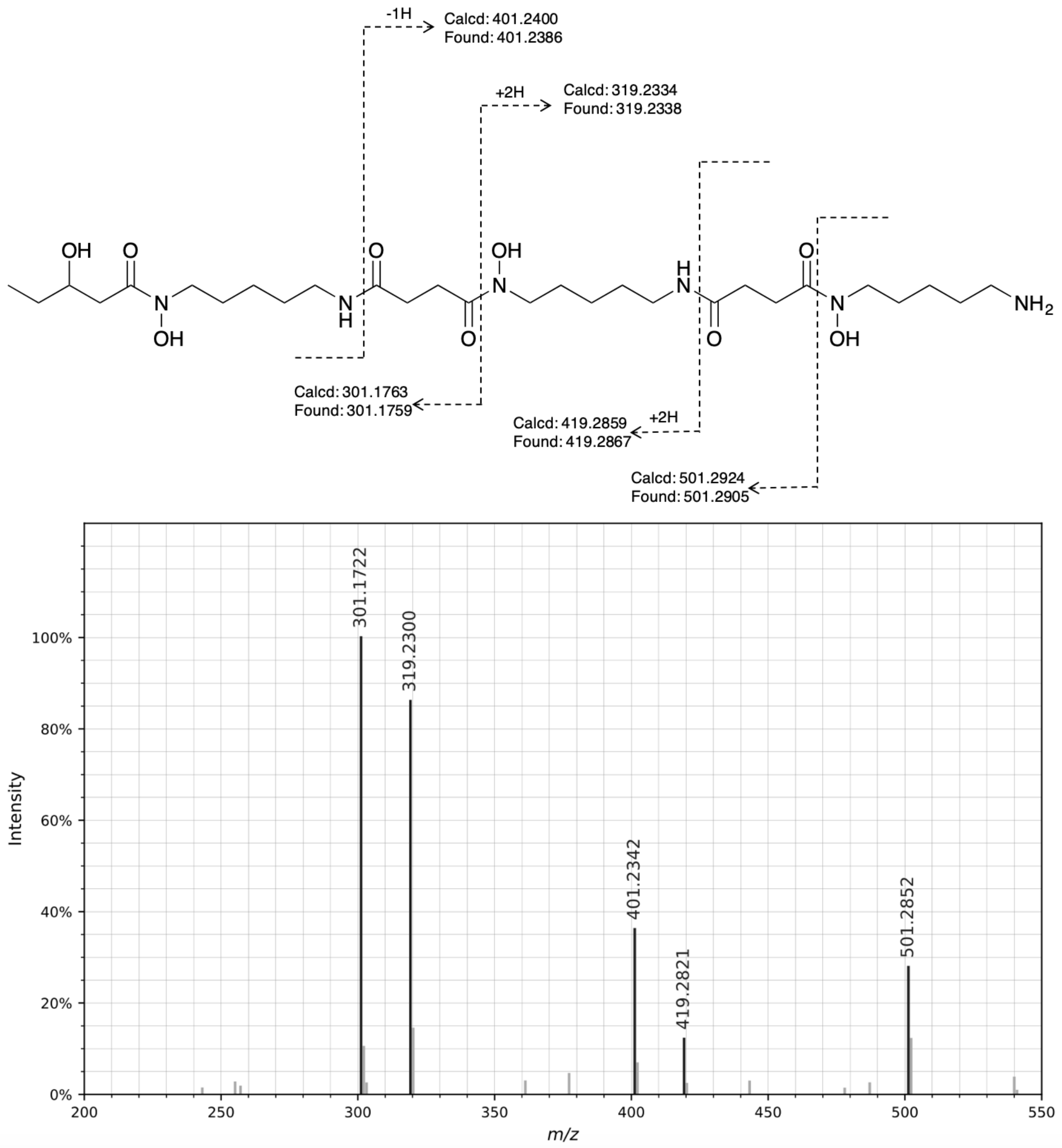


CID MS/MS spectrum of the $[M+H]^+$ ion of C4 acyl hydroxylated DFO-B. All fragment ions in the annotated structure match ions in the spectrum. Other fragment ions present in the spectrum are products of non-sequential fragmentation and are present in all DFOs.

Figure S27. Annotation of MS/MS for reported compounds: **uC6 acyl DFO-B**

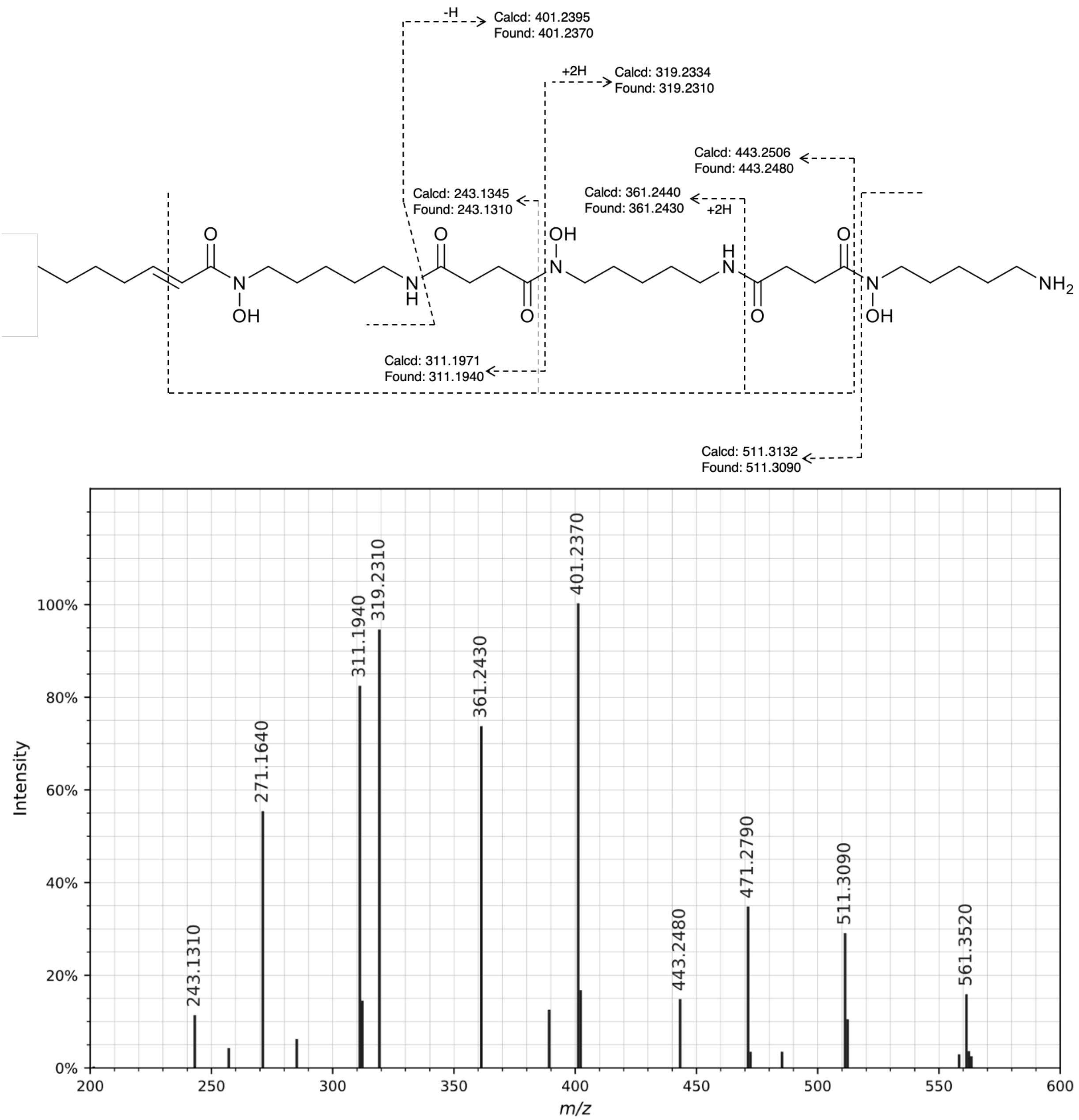


CID MS/MS spectrum of the [M+H]+ ion of uC6 acyl DFO-B. All fragment ions in the annotated structure match ions in the spectrum. Other fragment ions present in the spectrum are products of non-sequential fragmentation and are present in all DFOs.

Figure S28. Annotation of MS/MS for reported compounds: **C6 acyl DFO-B**

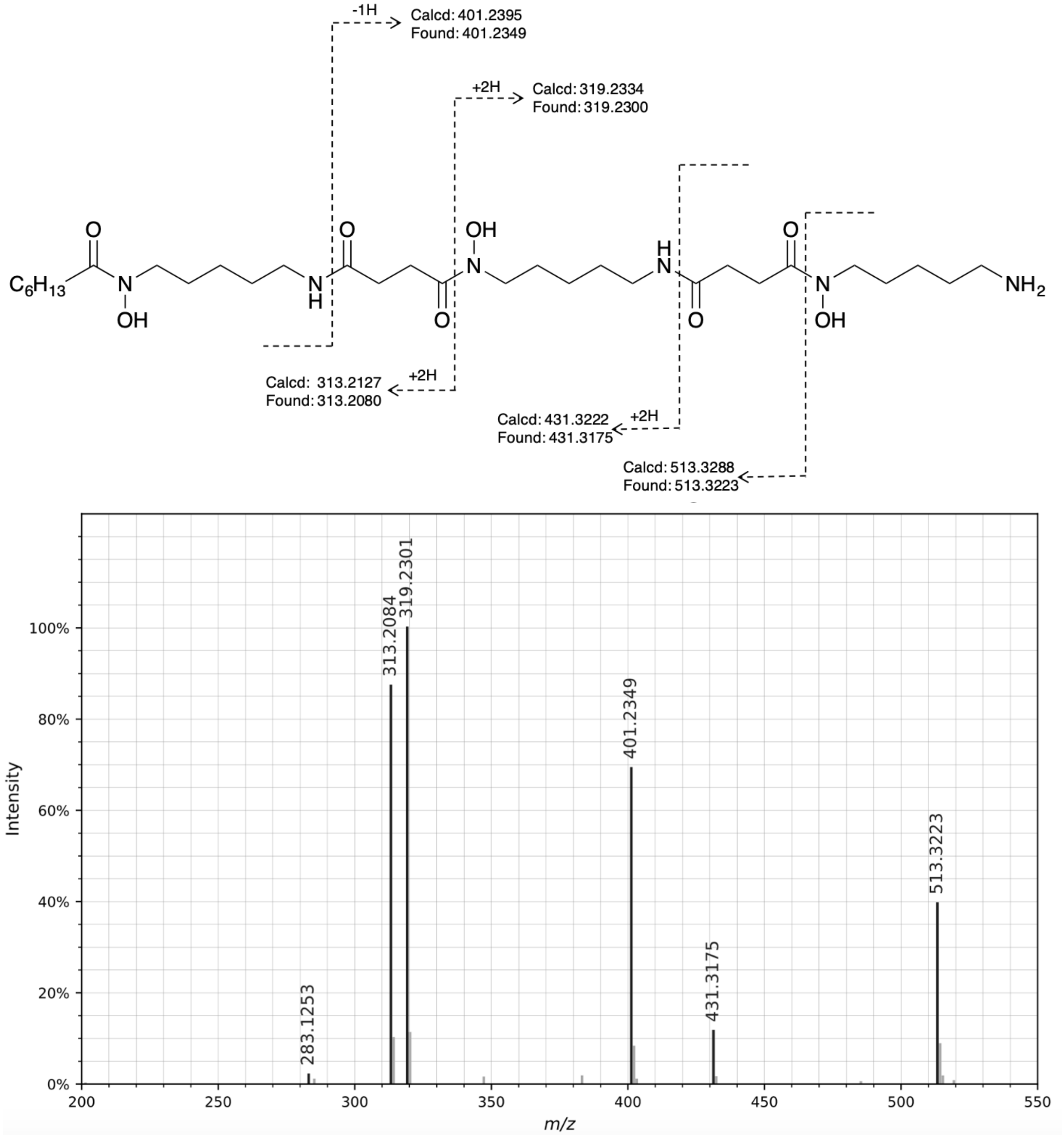


CID MS/MS spectrum of the $[M+H]^+$ ion of C6 acyl DFO-B. All fragment ions in the annotated structure match ions in the spectrum. Other fragment ions present in the spectrum are products of non-sequential fragmentation and are present in all DFOs.

Figure S29. Annotation of MS/MS for reported compounds: **uC7 acyl DFO-B**

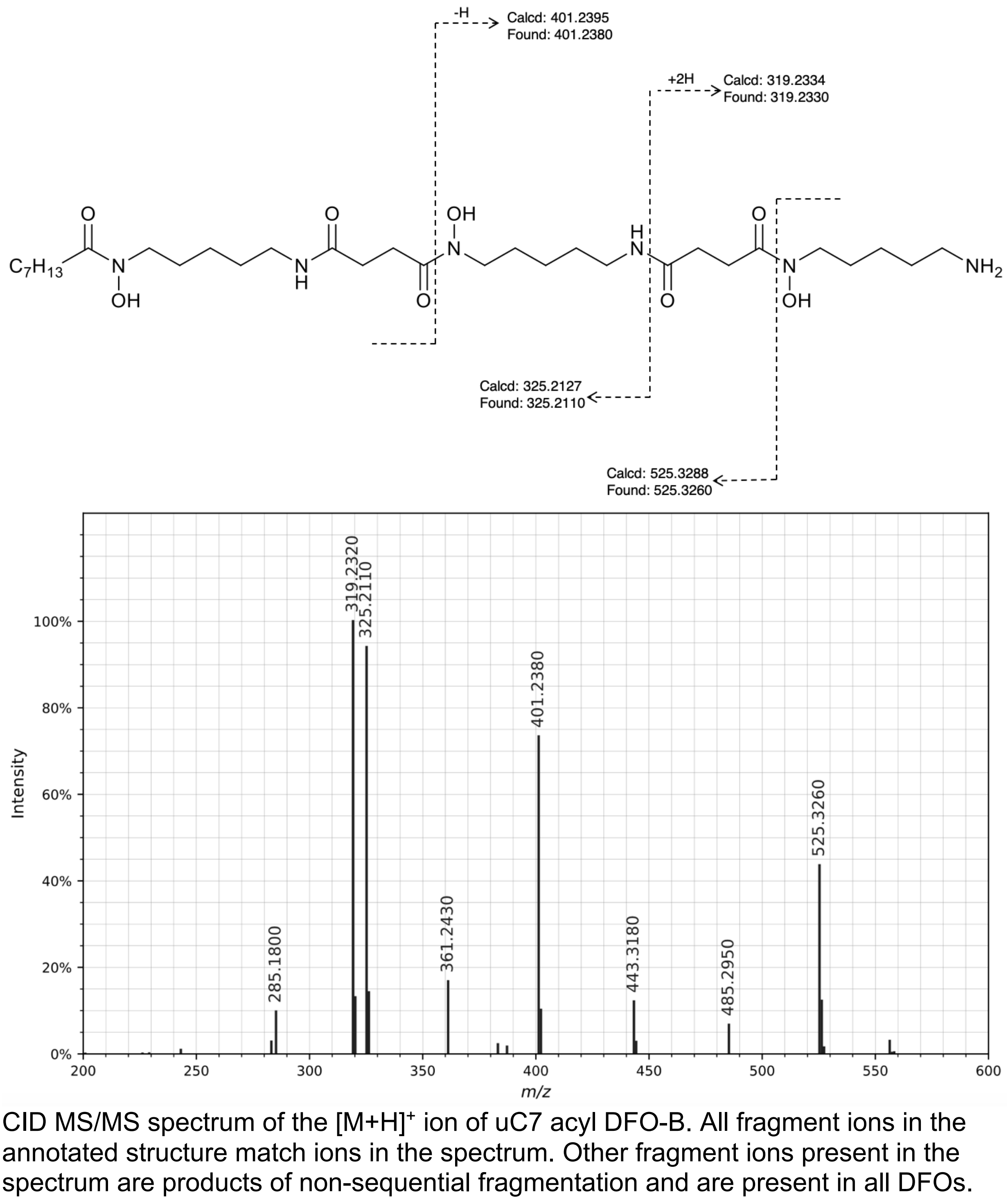


CID MS/MS spectrum of the $[M+H]^+$ ion of uC7 acyl DFO-B. All fragment ions in the annotated structure match ions in the spectrum. Other fragment ions present in the spectrum are products of non-sequential fragmentation and are present in all DFOs.

Figure S30. Annotation of MS/MS for reported compounds: **uC10 acyl DFO-B**

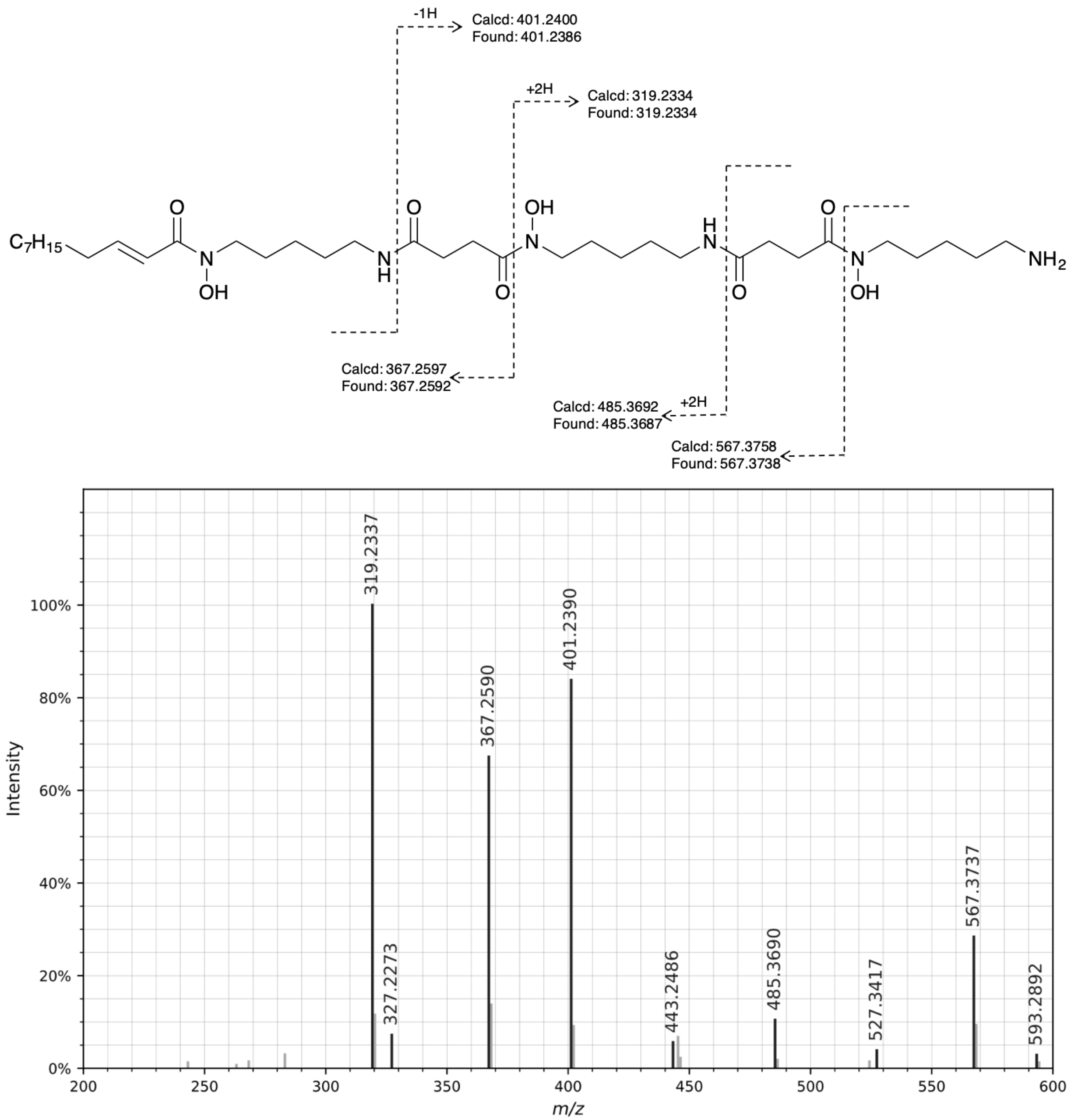


CID MS/MS spectrum of the $[M+H]^+$ ion of uC10 acyl DFO-B. All fragment ions in the annotated structure match ions in the spectrum. Other fragment ions present in the spectrum are products of non-sequential fragmentation and are present in all DFOs.

Figure S31.   Annotation of MS/MS for reported compounds: **uC11 acyl DFO-B**

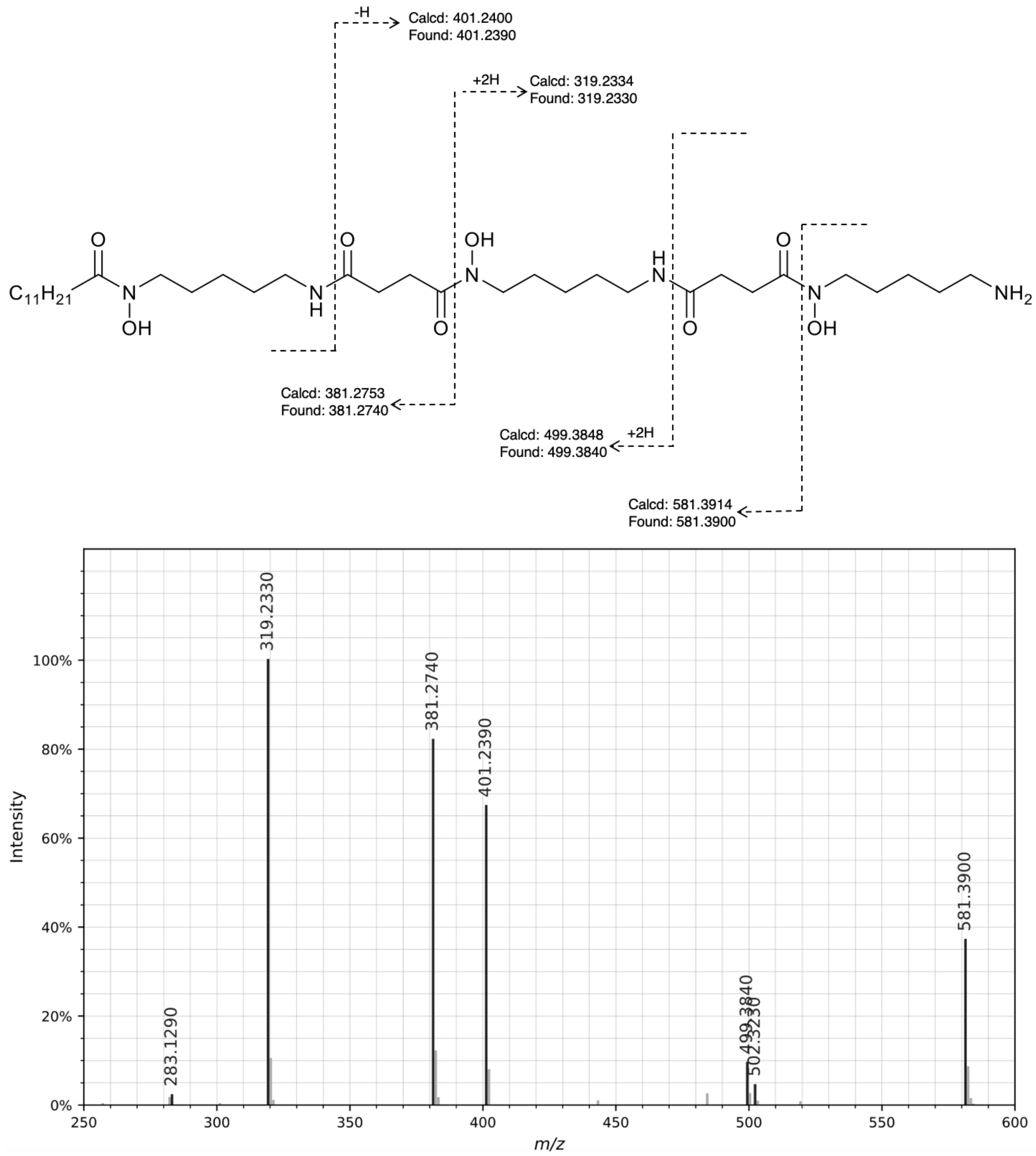


CID MS/MS spectrum of the $[M+H]^+$ ion of uC11 acyl DFO-B. All fragment ions in the annotated structure match ions in the spectrum. Other fragment ions present in the spectrum are products of non-sequential fragmentation and are present in all DFOs.

Figure S32. Annotation of MS/MS for reported compounds: **uC12 acyl FO-B**

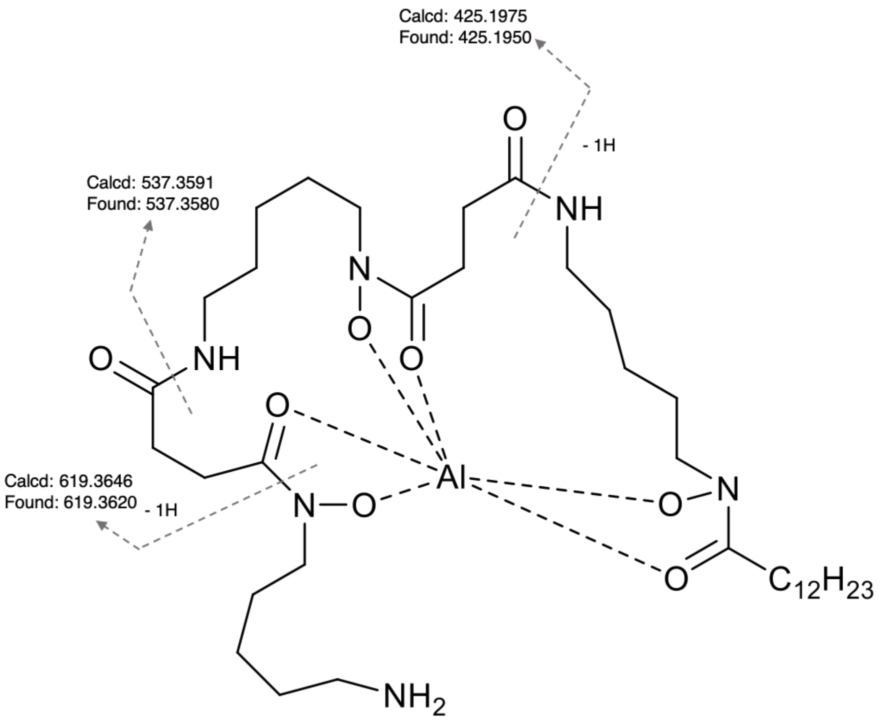


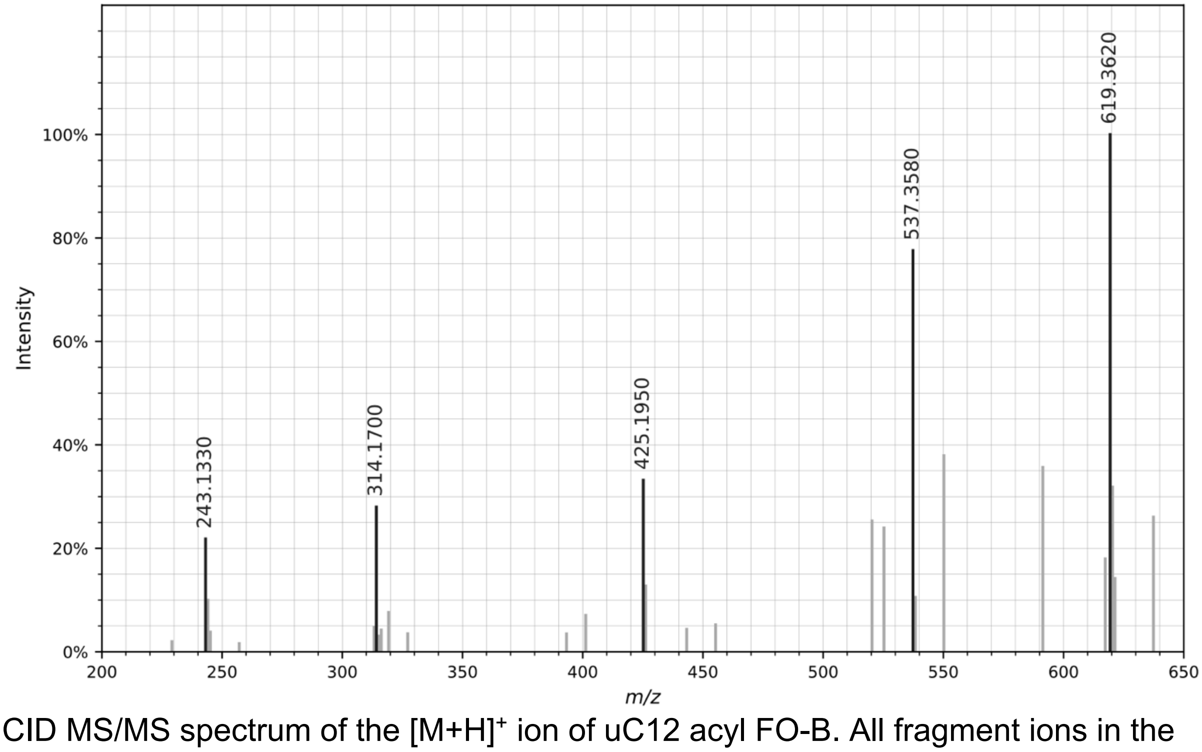


CID MS/MS spectrum of the $[M+H]^+$ ion of uC12 acyl FO-B. All fragment ions in the annotated structure match ions in the spectrum. Other fragment ions present in the spectrum are products of non-sequential fragmentation and are present in all DFOs.

Figure S33. Annotation of MS/MS for reported compounds: **dodecanedioic DFO-B**

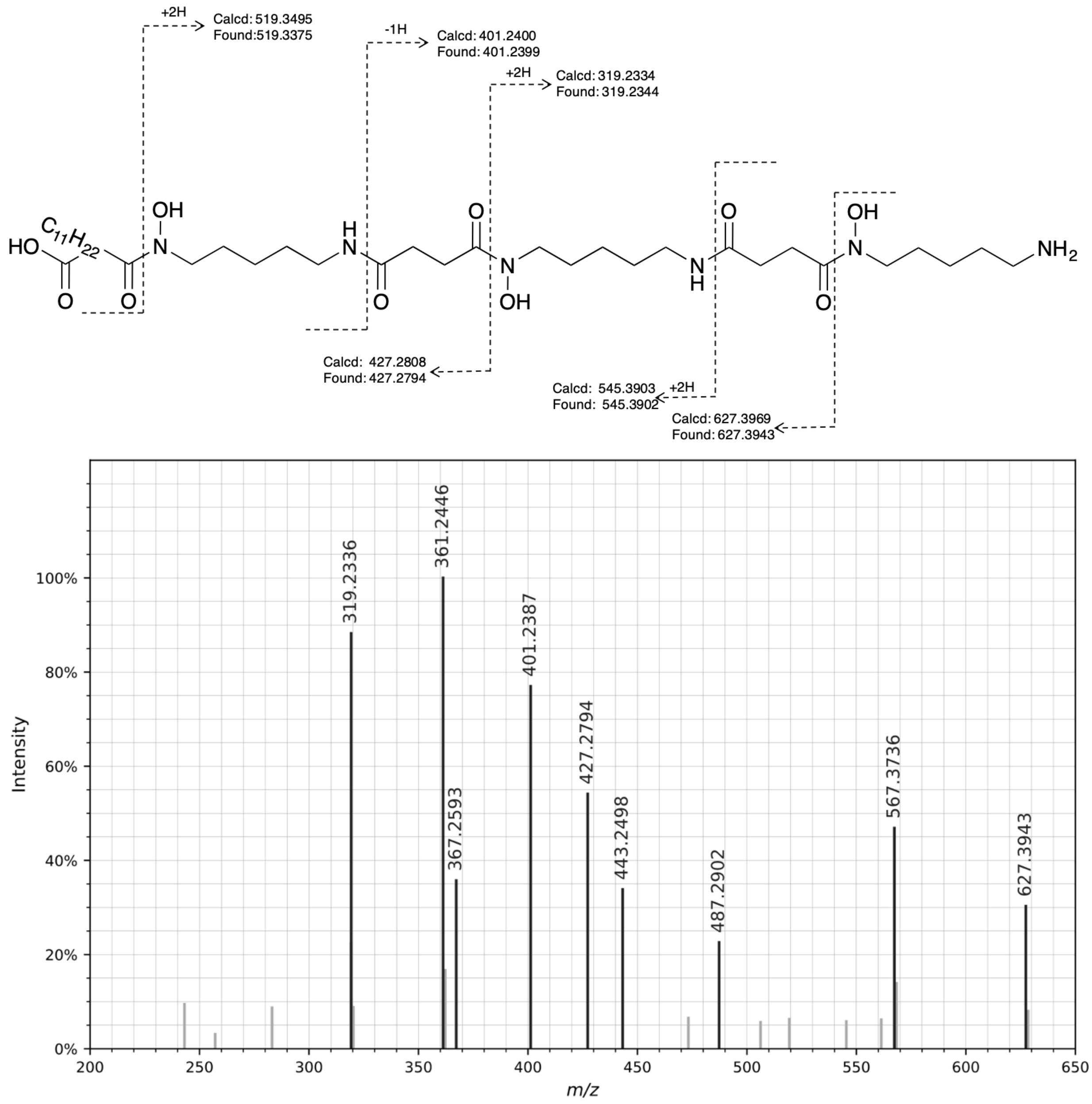


CID MS/MS spectrum of the $[M+H]^+$ ion of dodecanedioic DFO-B. All fragment ions in the annotated structure match ions in the spectrum. Other fragment ions present in the spectrum are products of non-sequential fragmentation and are present in all DFOs.

Figure S34. Annotation of MS/MS for reported compounds: **tridecanedioic DFO-B**

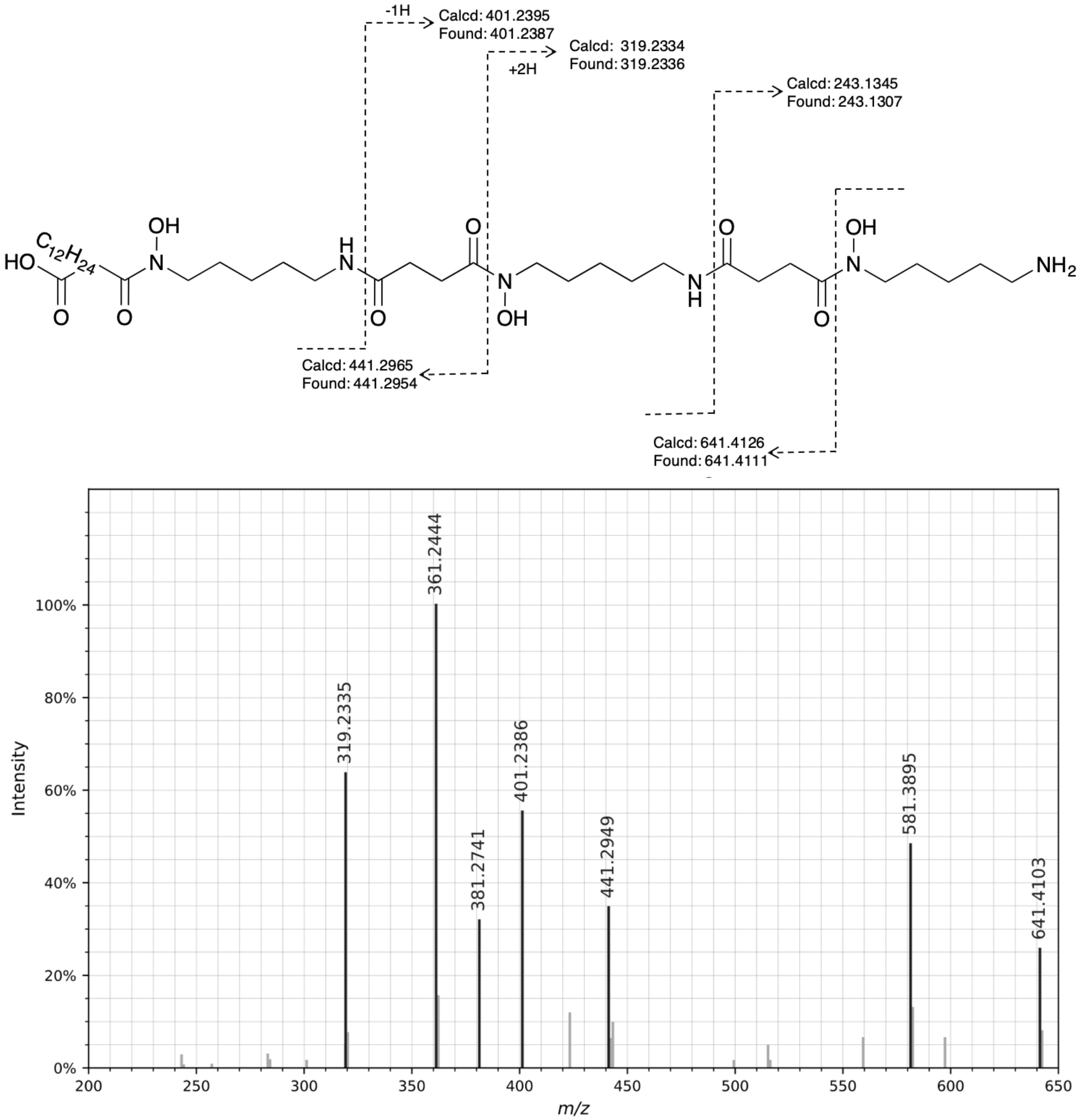


CID MS/MS spectrum of the $[M+H]^+$ ion of tridecanedioic DFO-B. All fragment ions in the annotated structure match ions in the spectrum. Other fragment ions present in the spectrum are products of non-sequential fragmentation and are present in all DFOs.

### DFO-D

Figure S35. Annotation of MS/MS for reported compounds: **glutaric desferrioxamine H**

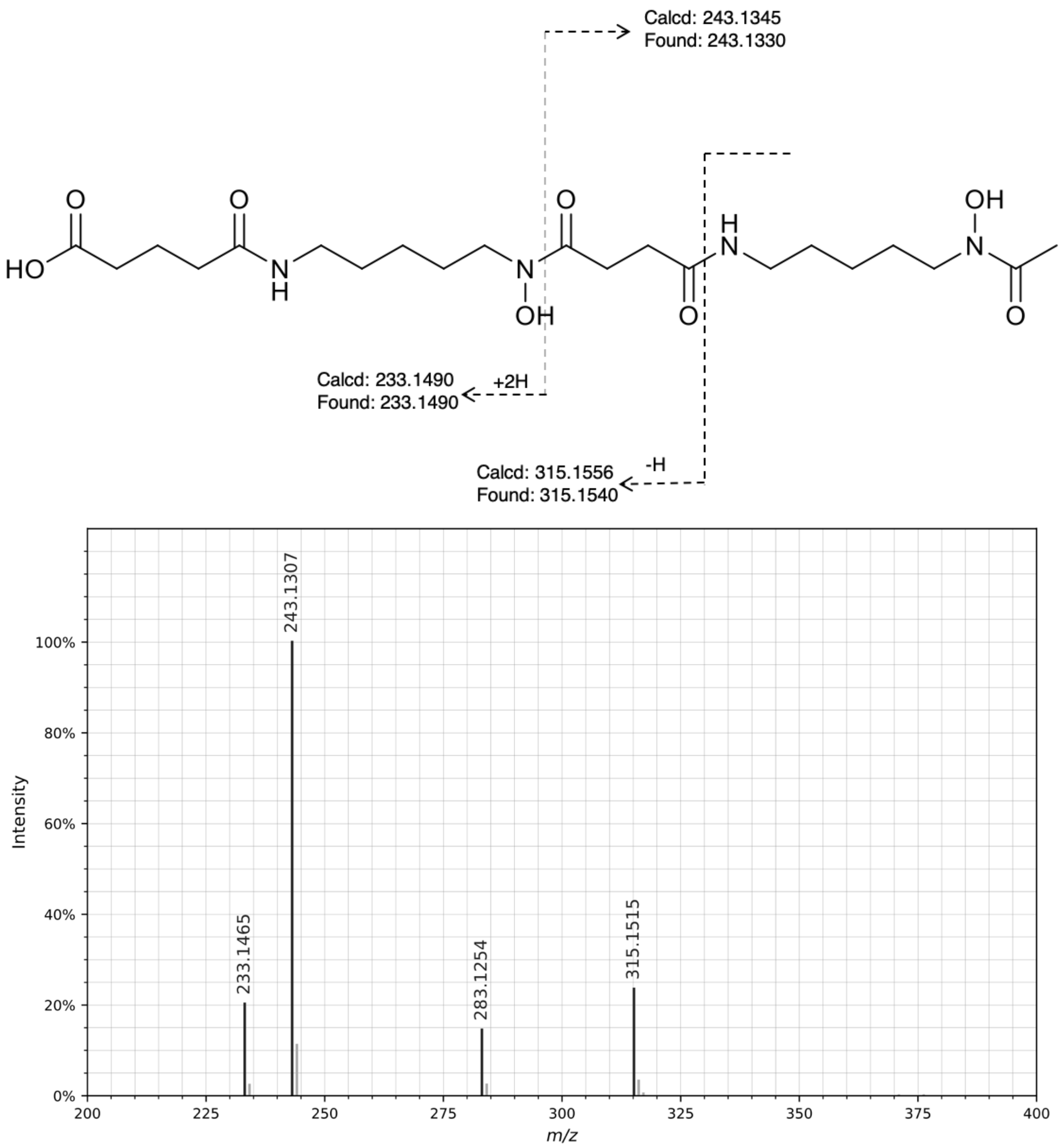


CID MS/MS spectrum of the $[M+H]^+$ ion of glutaric desferrioxamine H. All fragment ions in the annotated structure match ions in the spectrum. Other fragment ions present in the spectrum are products of non-sequential fragmentation and are present in all DFOs.

Figure S36. Annotation of MS/MS for reported compounds: **C2 acyl desferrioxamine H**

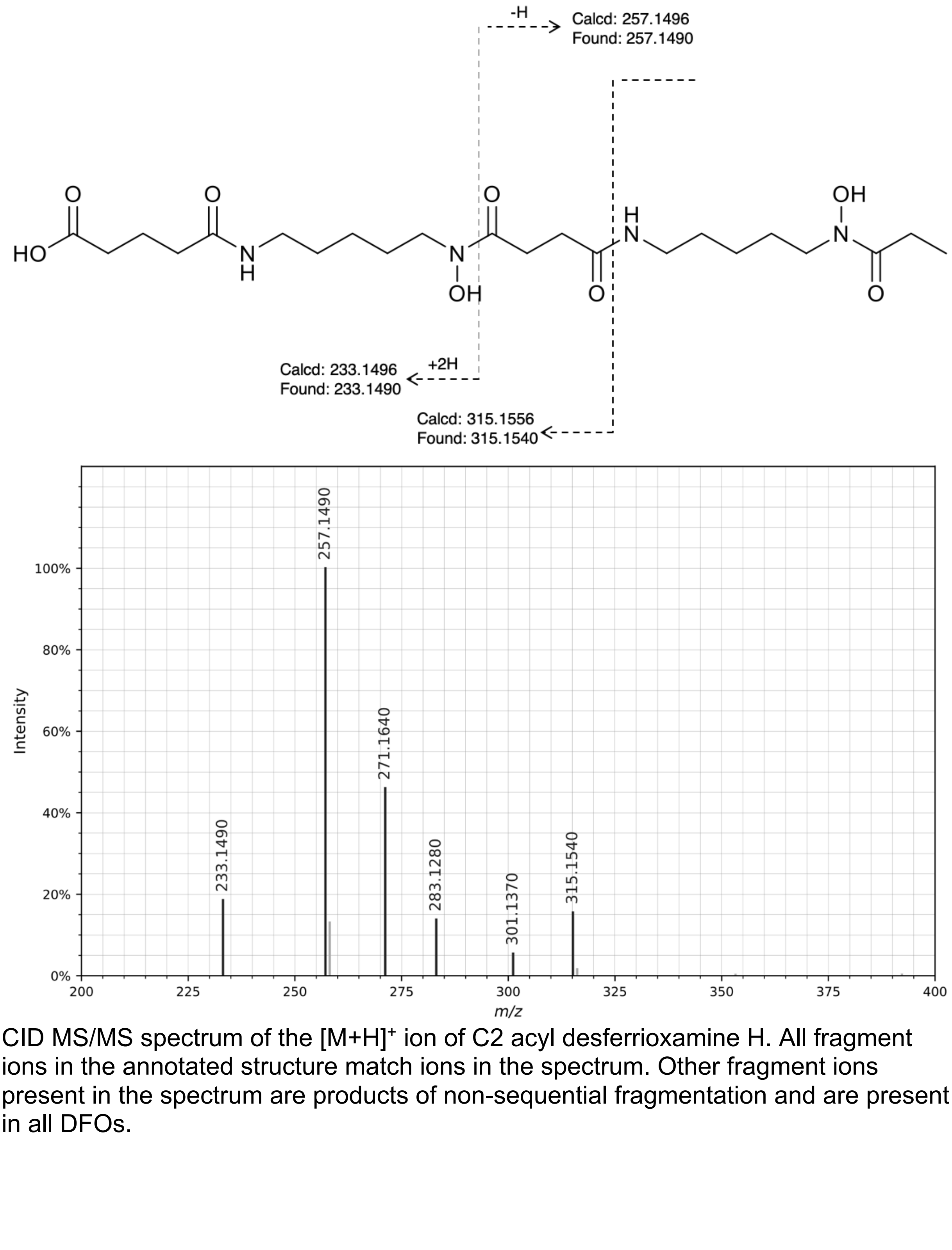


CID MS/MS spectrum of the $[M+H]^+$ ion of C2 acyl desferrioxamine H. All fragment ions in the annotated structure match ions in the spectrum. Other fragment ions present in the spectrum are products of non-sequential fragmentation and are present in all DFOs.

Figure S37. Annotation of MS/MS for reported compounds: **desferrioxamine D3**

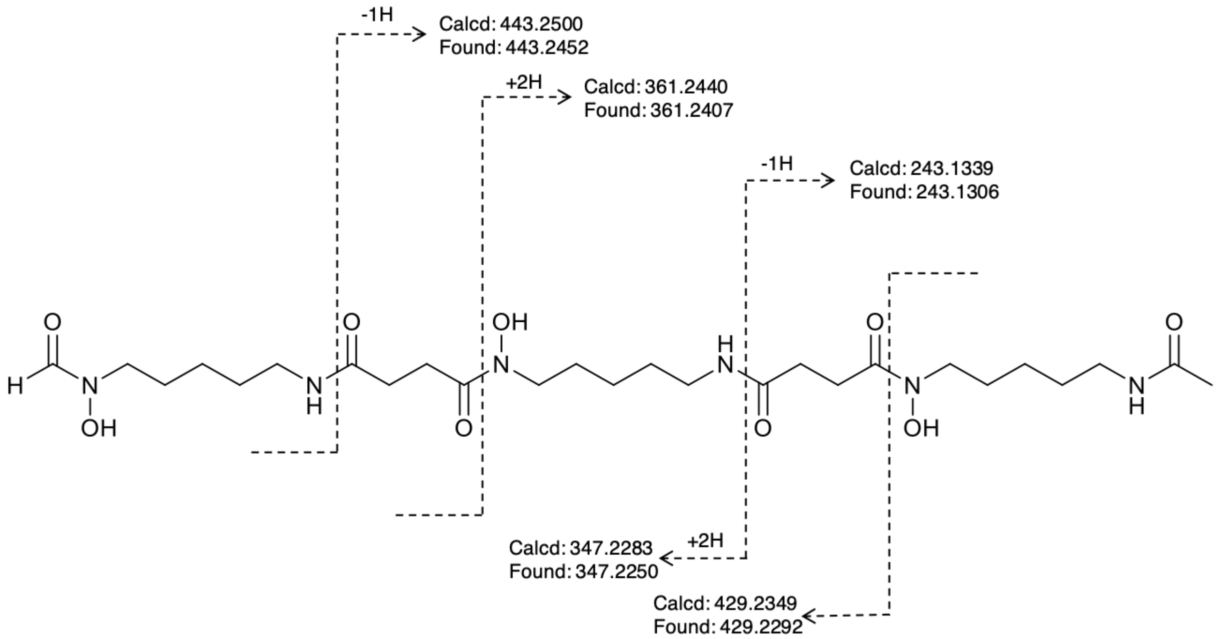


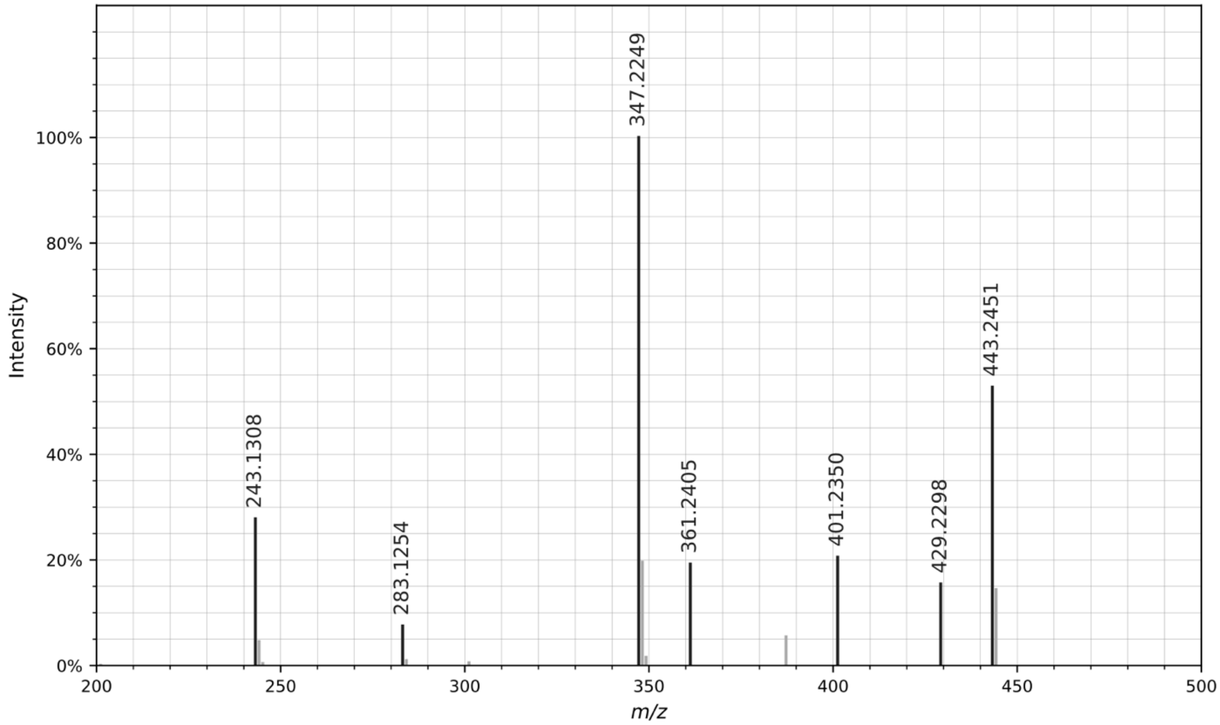


CID MS/MS spectrum of the $[M+H]^+$ ion of desferrioxamine D3. All fragment ions in the annotated structure match ions in the spectrum. Other fragment ions present in the spectrum are products of non-sequential fragmentation and are present in all DFOs.

Figure S38. Annotation of MS/MS for reported compounds: **C2 acyl DFO-D**

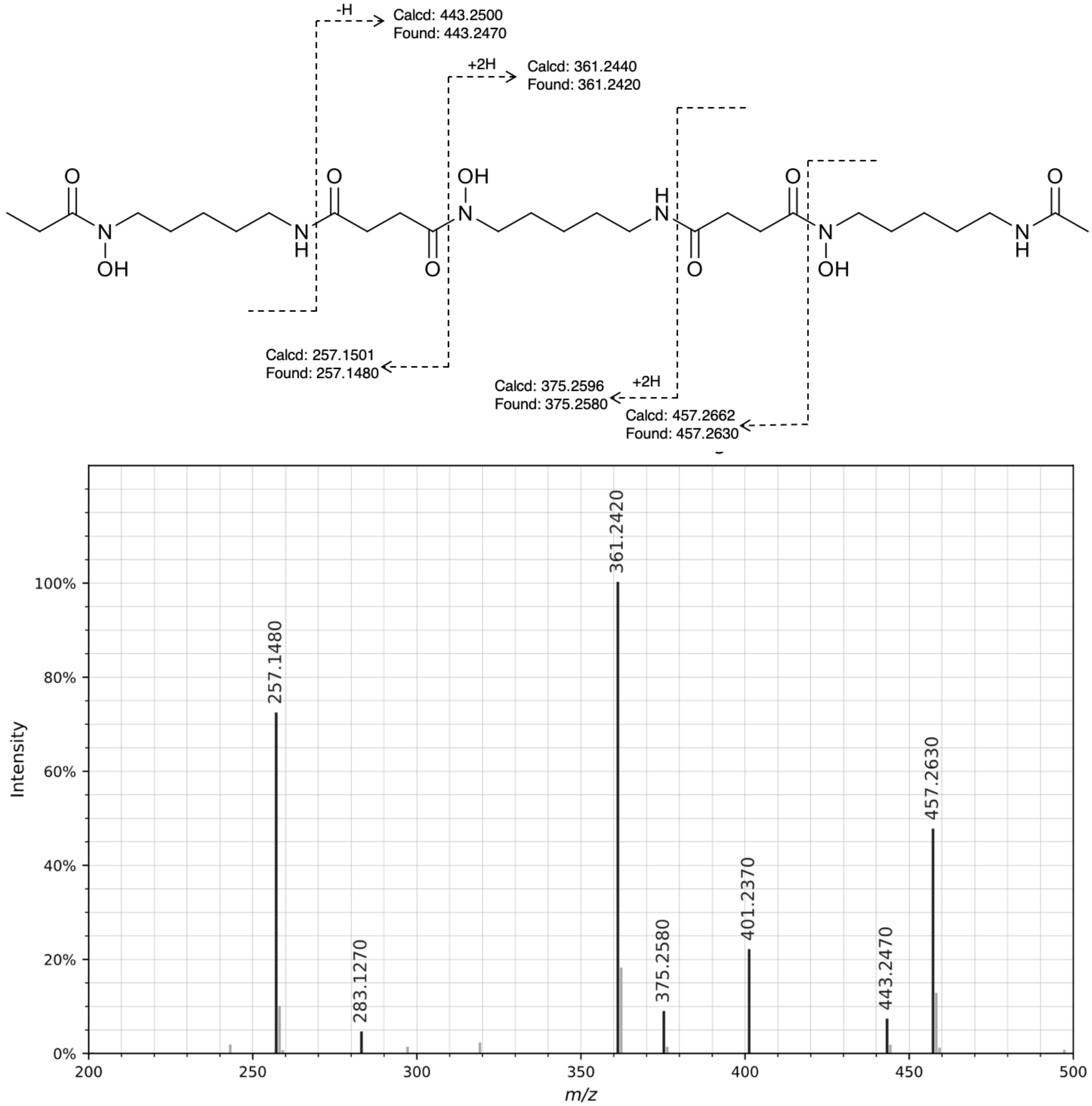


CID MS/MS spectrum of the $[M+H]^+$ ion of C2 acyl DFO-D. All fragment ions in the annotated structure match ions in the spectrum. Other fragment ions present in the spectrum are products of non-sequential fragmentation and are present in all DFOs.

Figure S39. Annotation of MS/MS for reported compounds: **C4 acyl DFO-D**

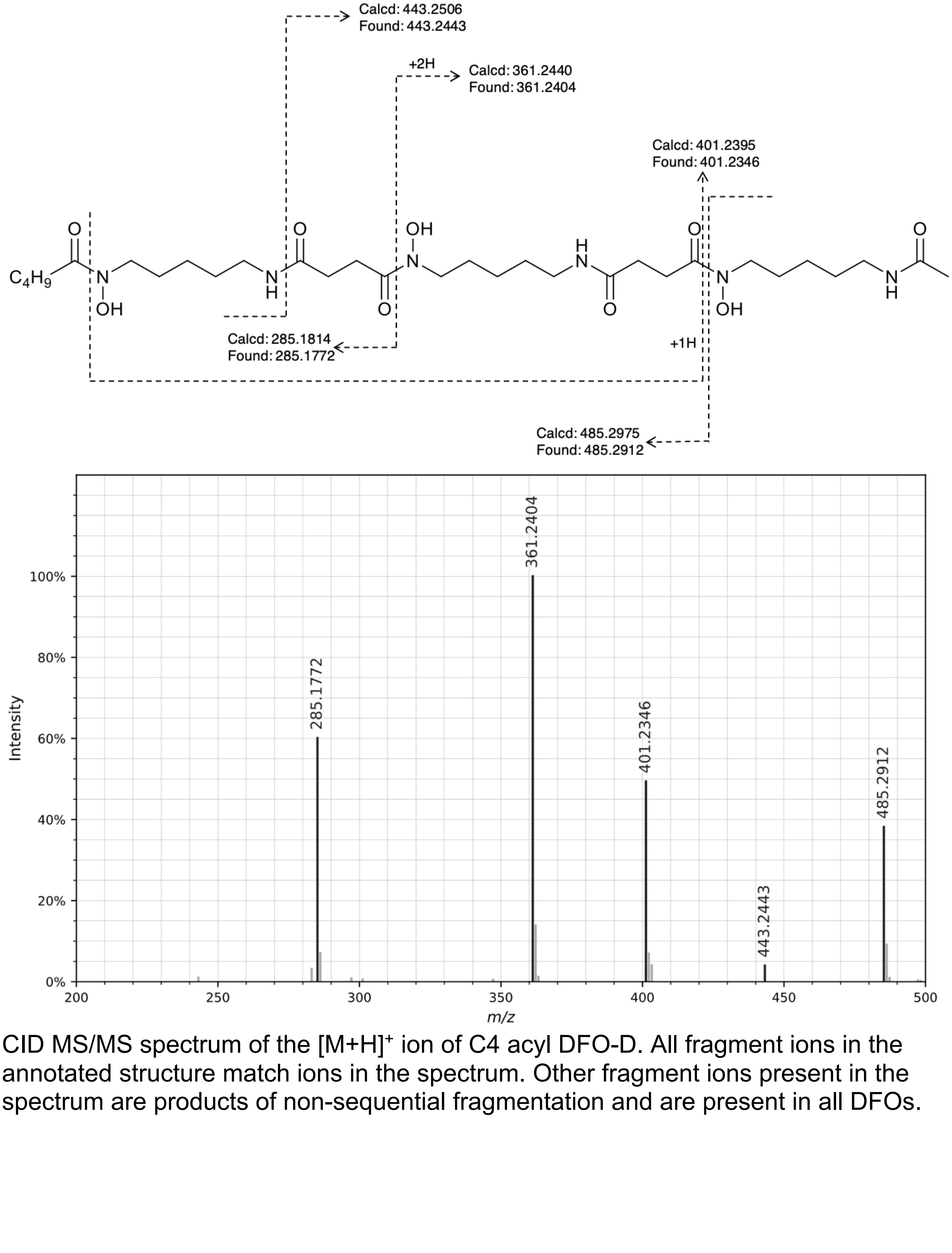


CID MS/MS spectrum of the $[M+H]^+$ ion of C4 acyl DFO-D. All fragment ions in the annotated structure match ions in the spectrum. Other fragment ions present in the spectrum are products of non-sequential fragmentation and are present in all DFOs.

Figure S40.   Annotation of MS/MS for reported compounds: **C5 acyl DFO-D**

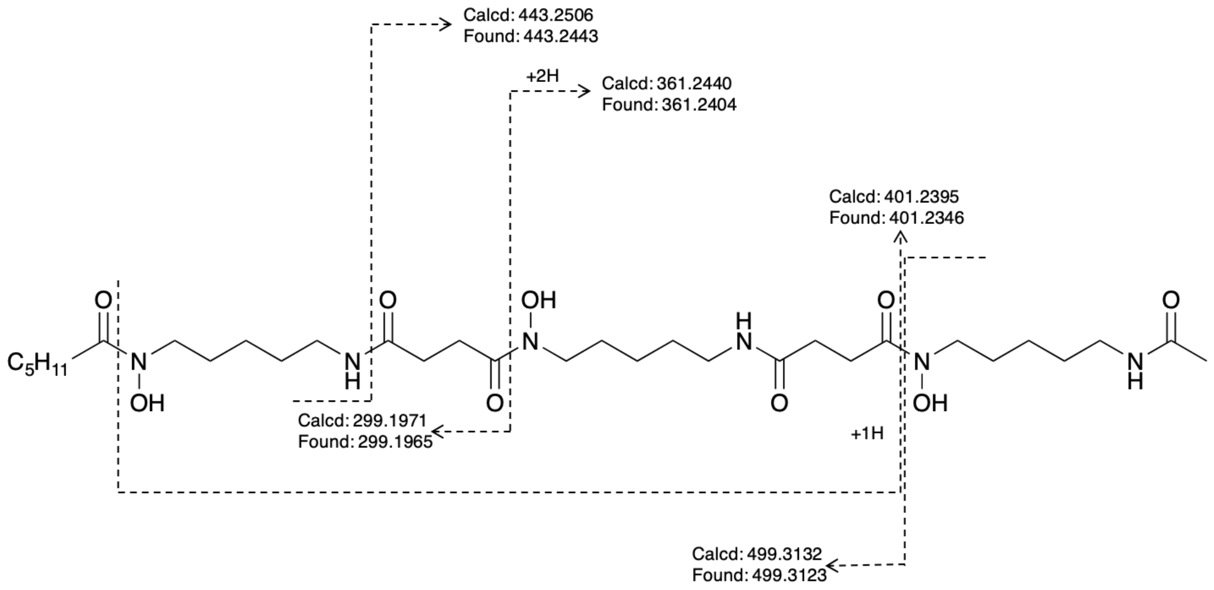


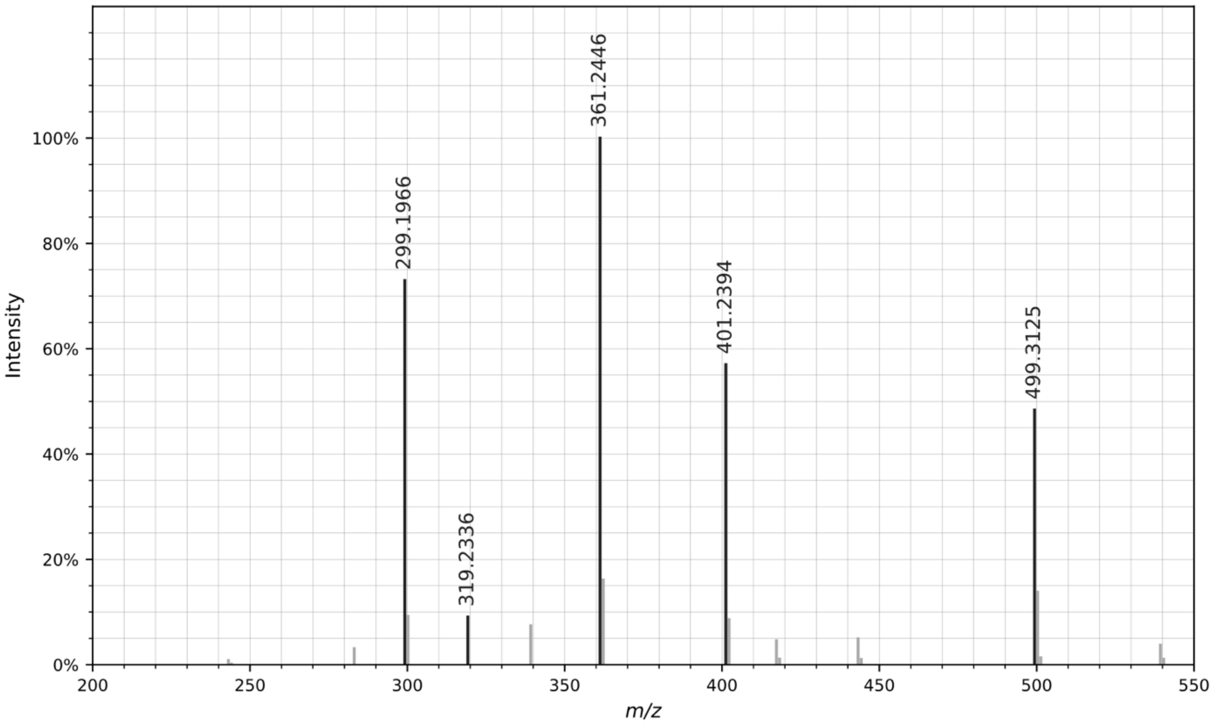


CID MS/MS spectrum of the $[M+H]^+$ ion of C5 acyl DFO-D. All fragment ions in the annotated structure match ions in the spectrum. Other fragment ions present in the spectrum are products of non-sequential fragmentation and are present in all DFOs.

Figure S41. Annotation of MS/MS for reported compounds: **C6 acyl DFO-D**

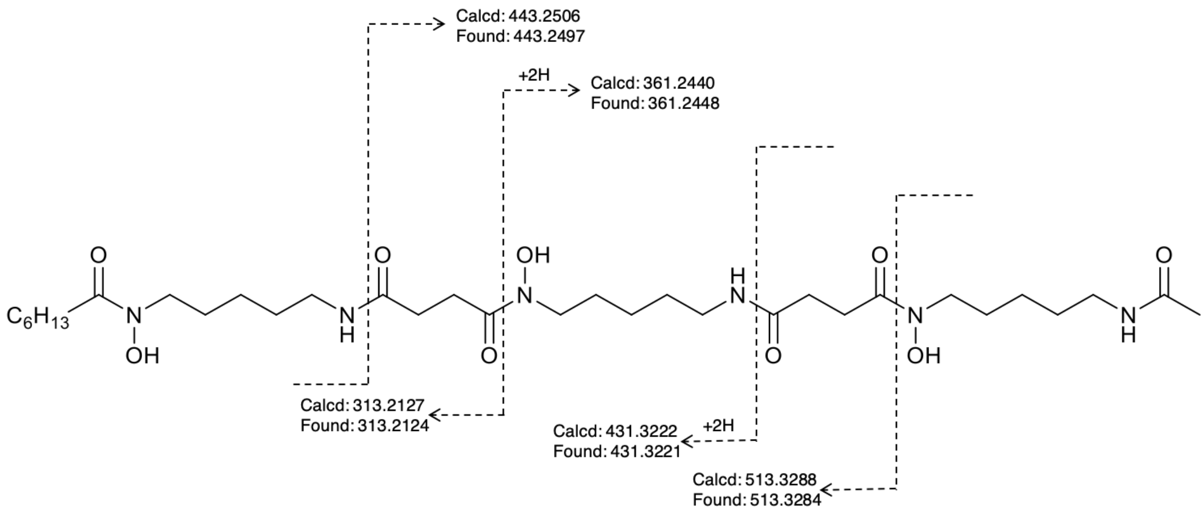


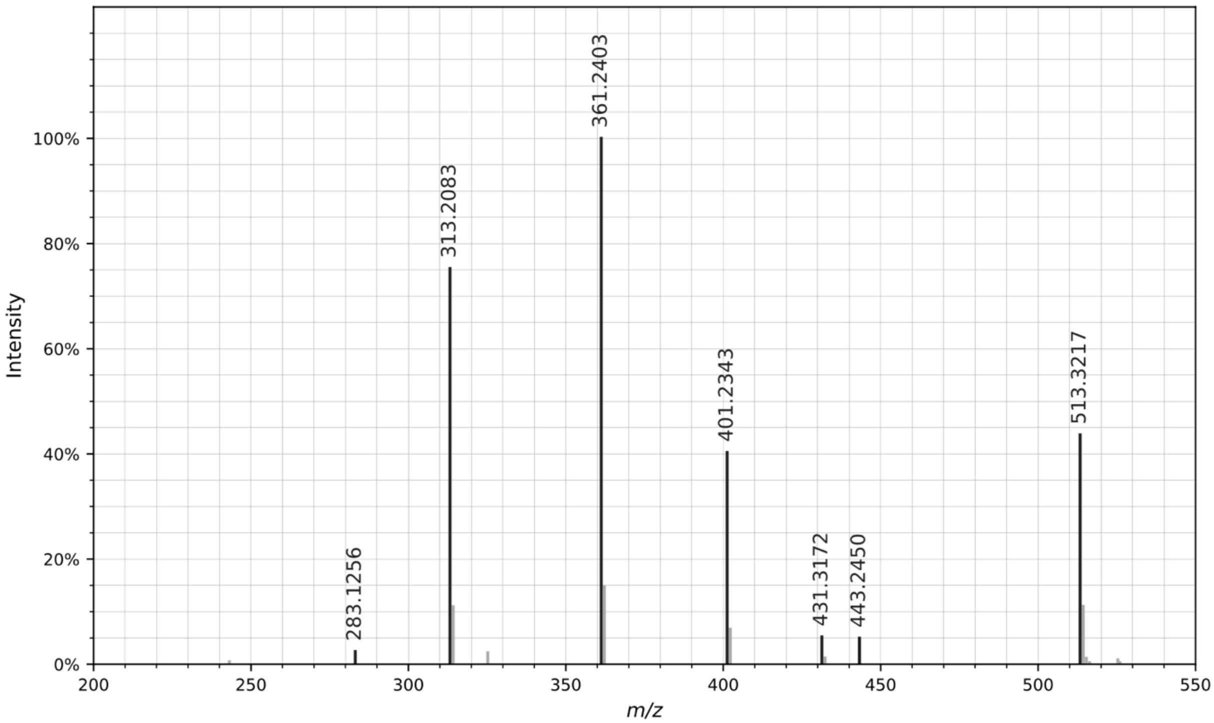


CID MS/MS spectrum of the $[M+H]^+$ ion of C6 acyl DFO-D. All fragment ions in the annotated structure match ions in the spectrum. Other fragment ions present in the spectrum are products of non-sequential fragmentation and are present in all DFOs.

Figure S42. Annotation of MS/MS for reported compounds: **C7 acyl DFO-D**

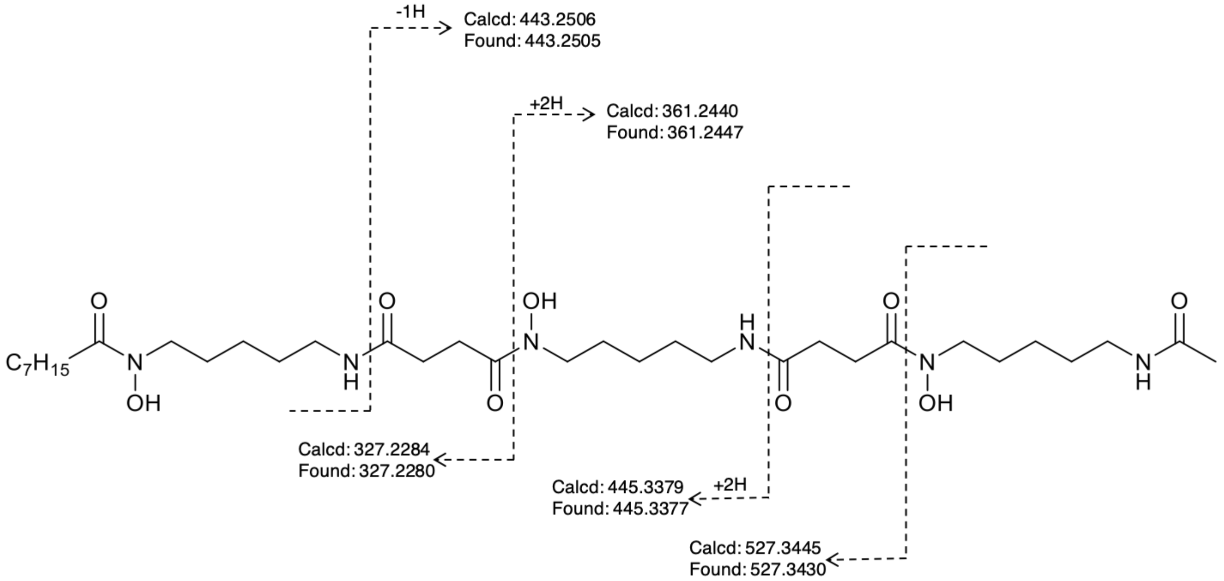


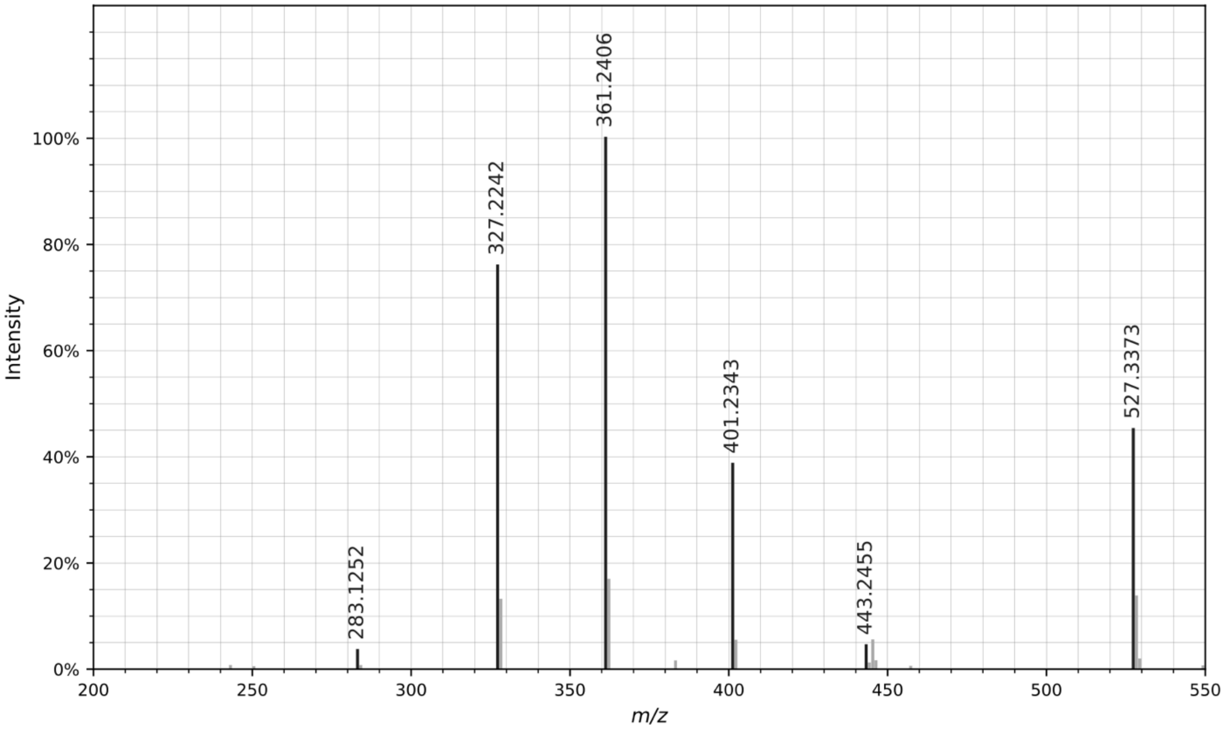


CID MS/MS spectrum of the $[M+H]^+$ ion of C7 acyl DFO-D. All fragment ions in the annotated structure match ions in the spectrum. Other fragment ions present in the spectrum are products of non-sequential fragmentation and are present in all DFOs.

Figure S43. Annotation of MS/MS for reported compounds: **C11 acyl DFO-D**

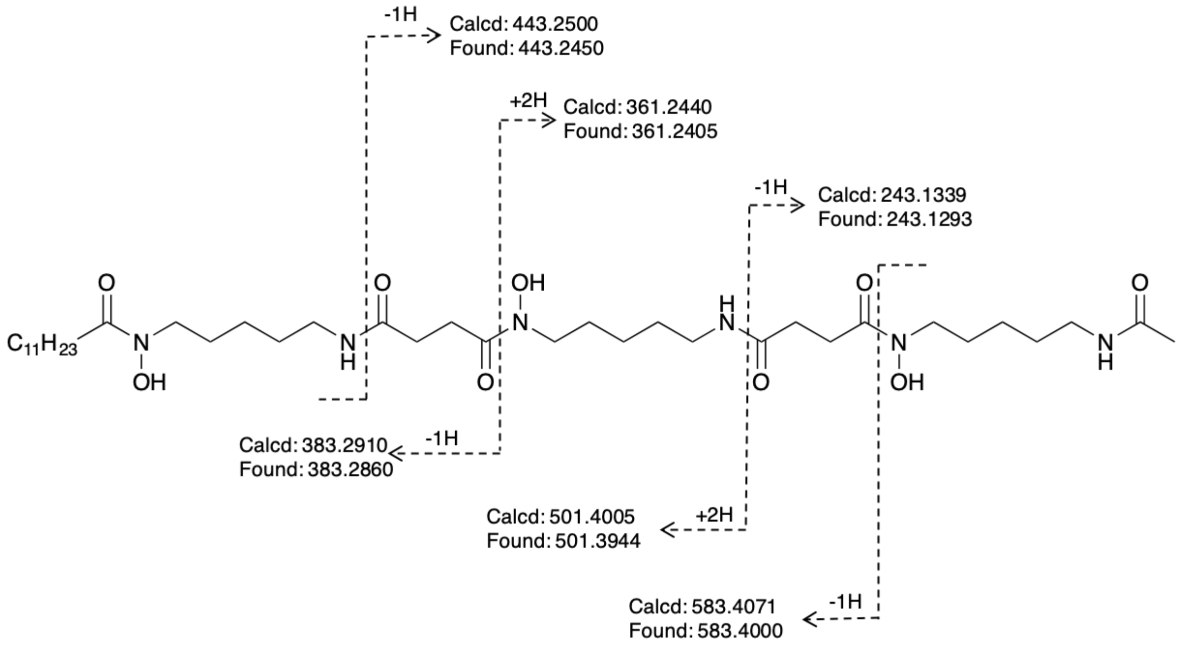


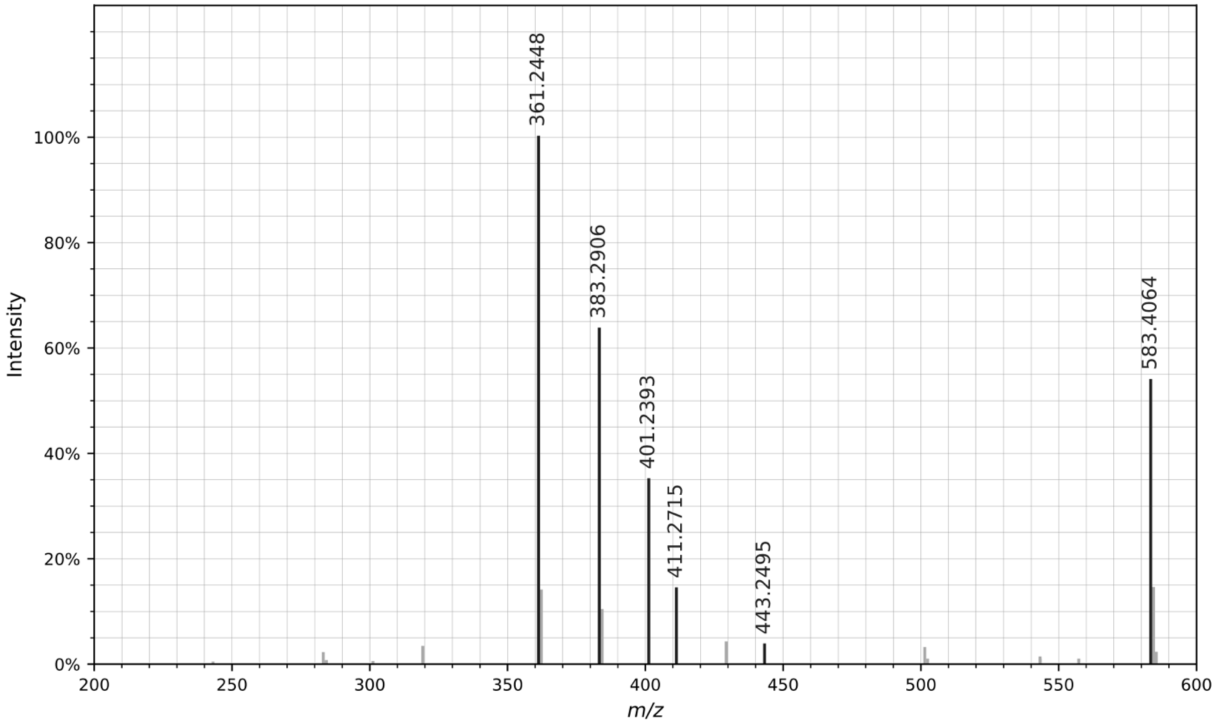


CID MS/MS spectrum of the $[M+H]^+$ ion of C11 acyl DFO-D. All fragment ions in the annotated structure match ions in the spectrum. Other fragment ions present in the spectrum are products of non-sequential fragmentation and are present in all DFOs.

Figure S44. Annotation of MS/MS for reported compounds: **C12 acyl DFO-D**

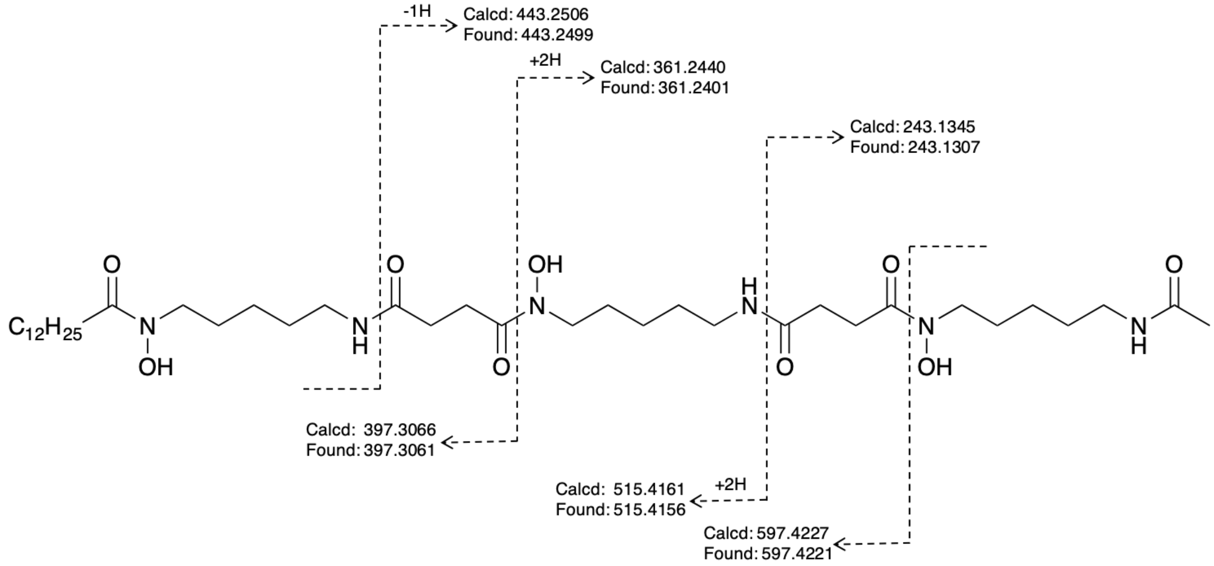


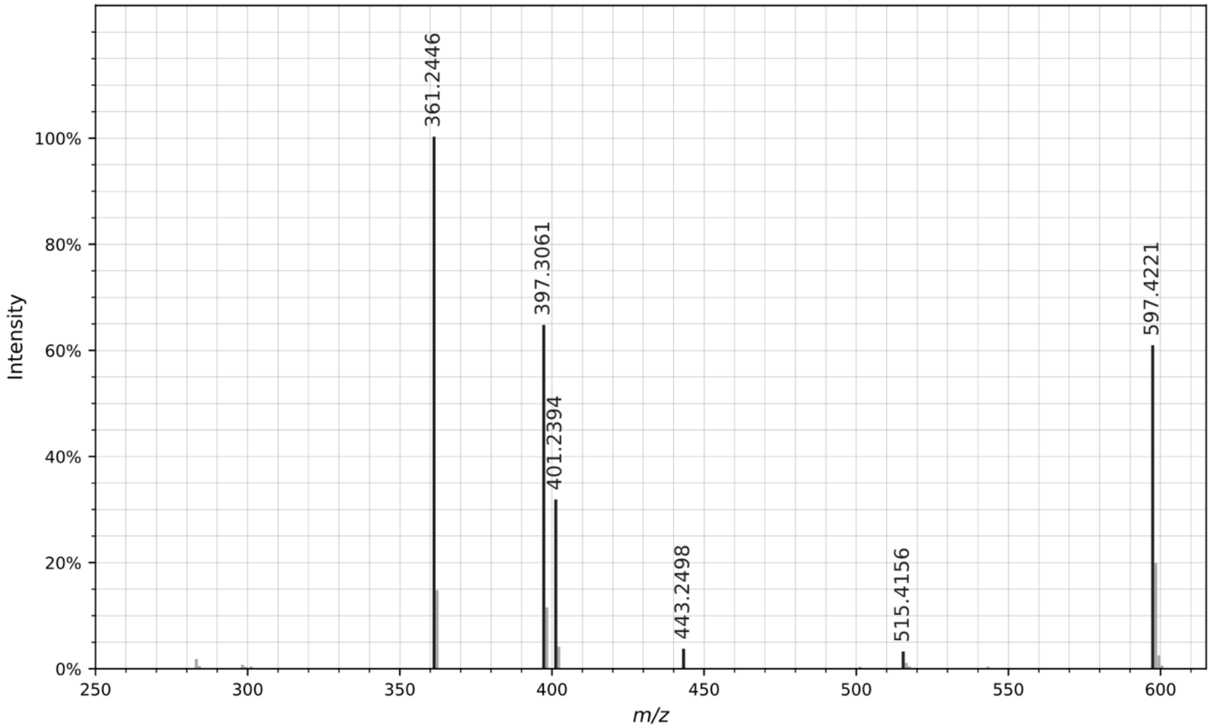


CID MS/MS spectrum of the $[M+H]^+$ ion of C12 acyl DFO-D. All fragment ions in the annotated structure match ions in the spectrum. Other fragment ions present in the spectrum are products of non-sequential fragmentation and are present in all DFOs.

Figure S45. Annotation of MS/MS for reported compounds: **C13 acyl DFO-D**

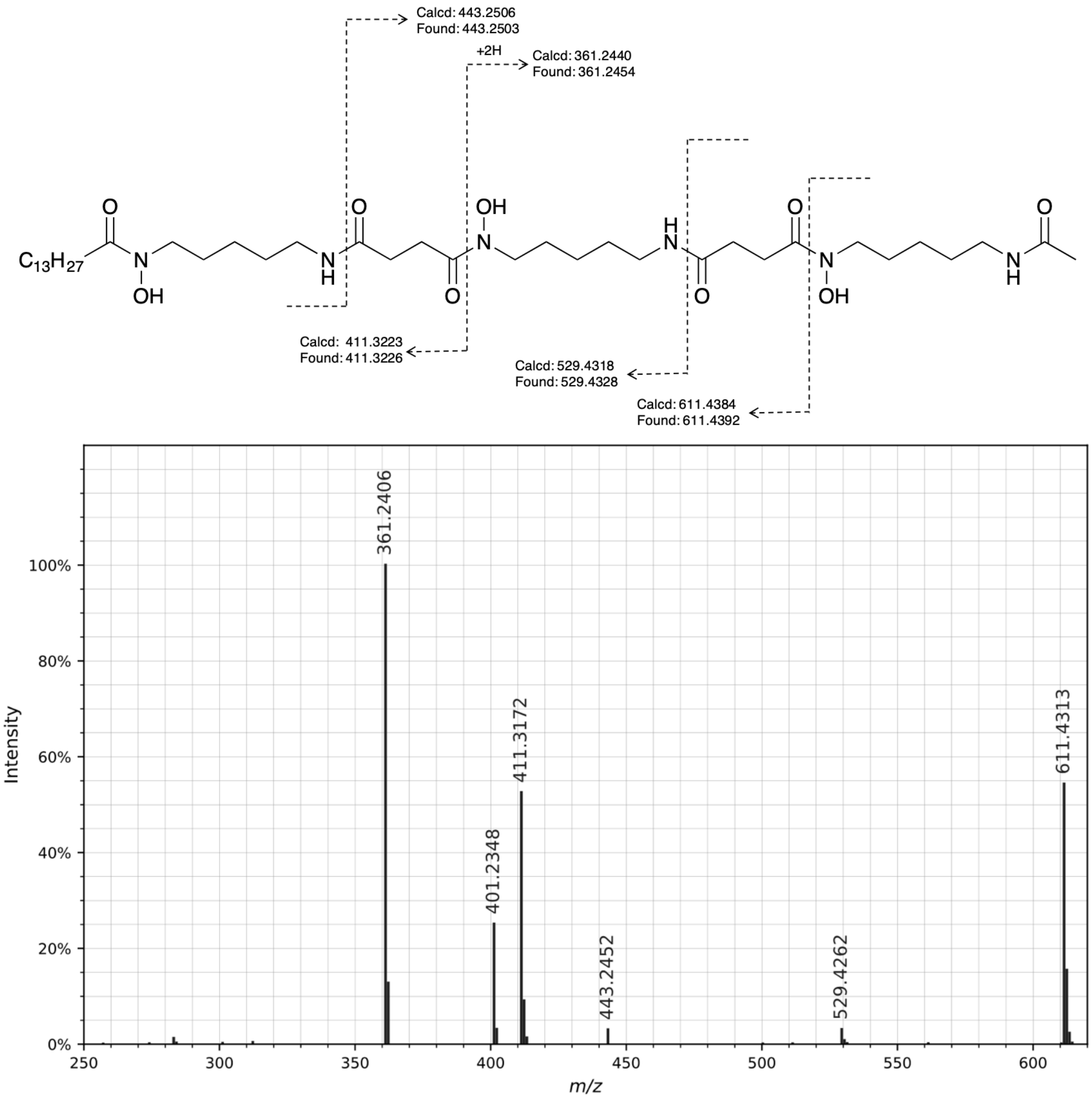


CID MS/MS spectrum of the $[M+H]^+$ ion of C13 acyl DFO-D. All fragment ions in the annotated structure match ions in the spectrum. Other fragment ions present in the spectrum are products of non-sequential fragmentation and are present in all DFOs.

Figure S46.   Annotation of MS/MS for reported compounds: **C14 acyl DFO-D**

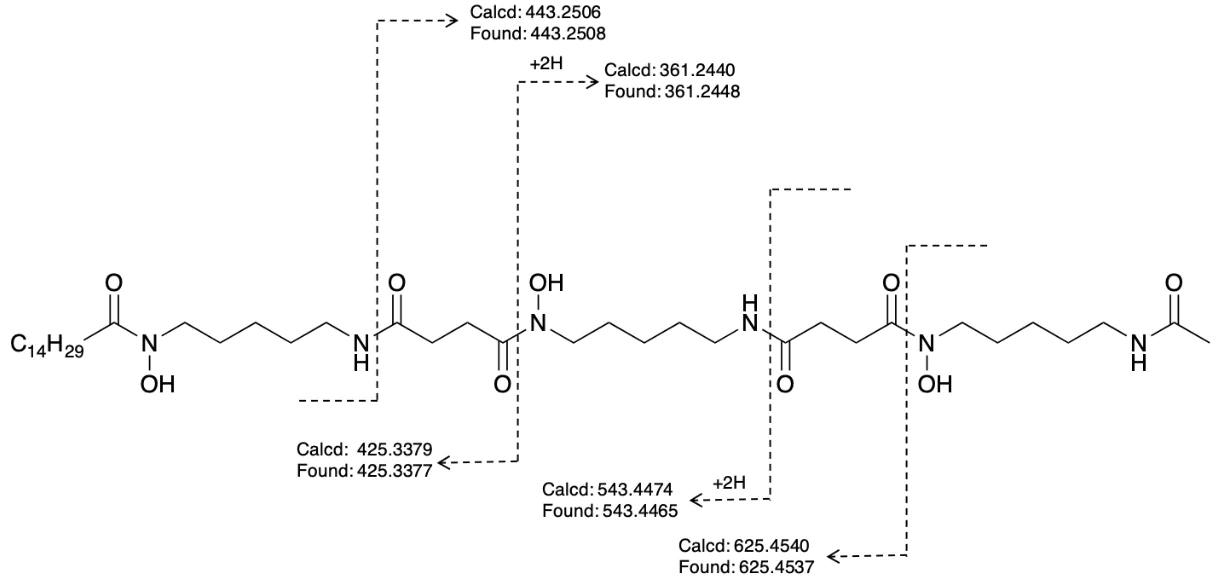


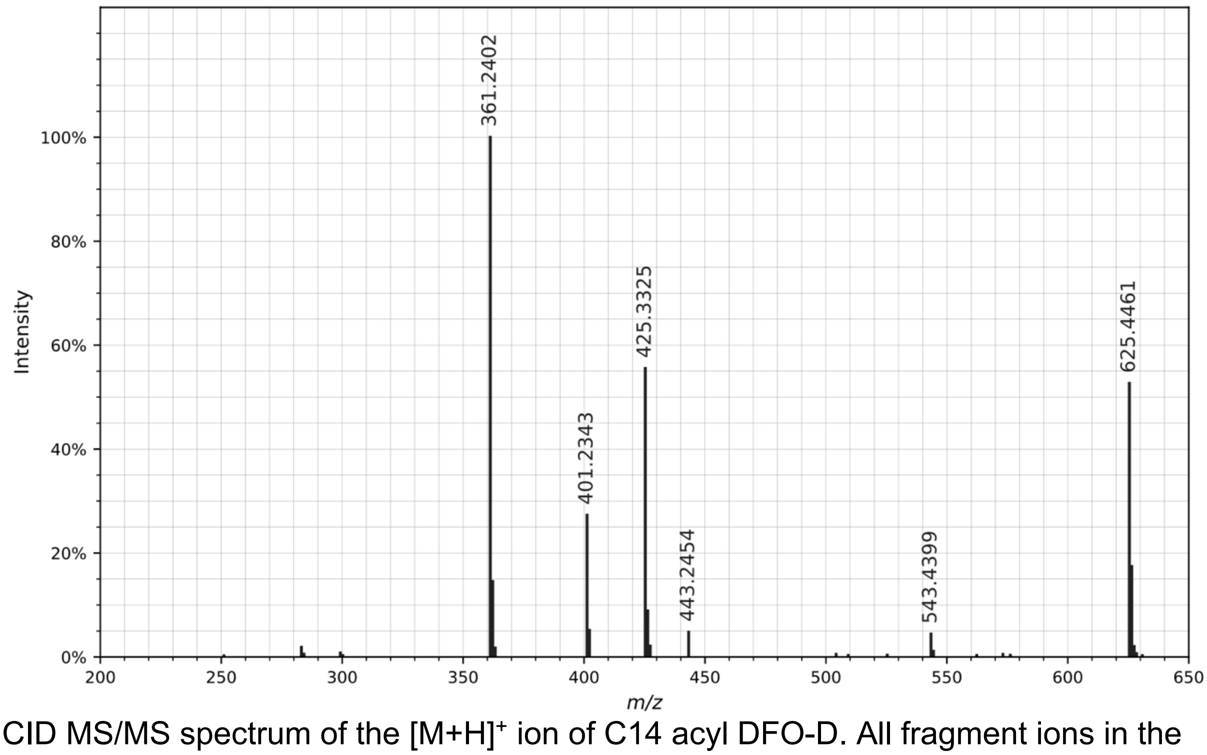


CID MS/MS spectrum of the $[M+H]^+$ ion of C14 acyl DFO-D. All fragment ions in the annotated structure match ions in the spectrum. Other fragment ions present in the spectrum are products of non-sequential fragmentation and are present in all DFOs.

Figure S47. Annotation of MS/MS for reported compounds: **tenacibactin E**

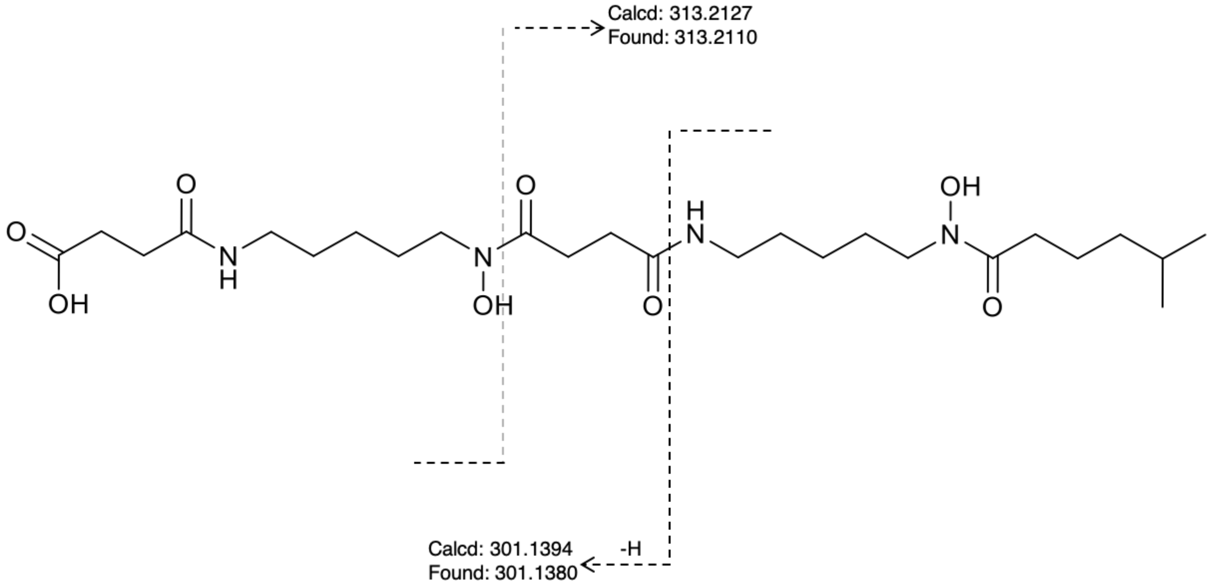


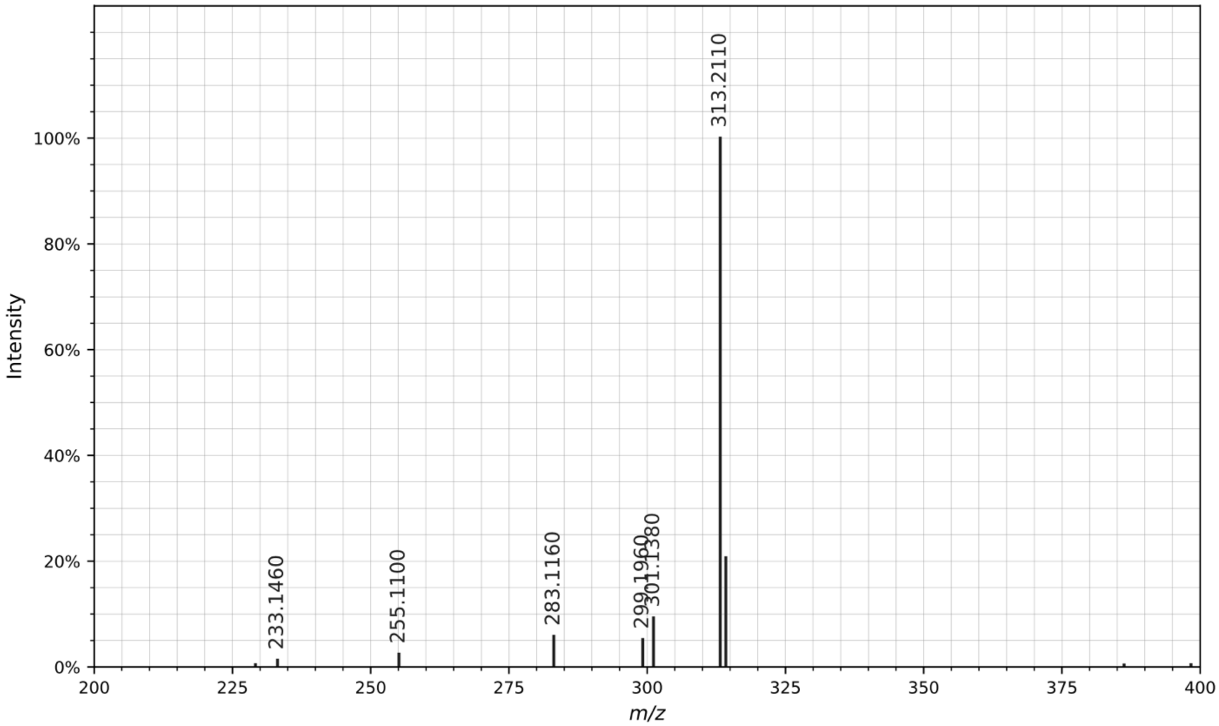


CID MS/MS spectrum of the [M+H]$^{+}$ ion of tenacibactin E. All fragment ions in the annotated structure match ions in the spectrum. Other fragment ions present in the spectrum are products of non-sequential fragmentation and are present in all DFOs.

Figure S48. Annotation of MS/MS for reported compounds: **tenacibactin F**

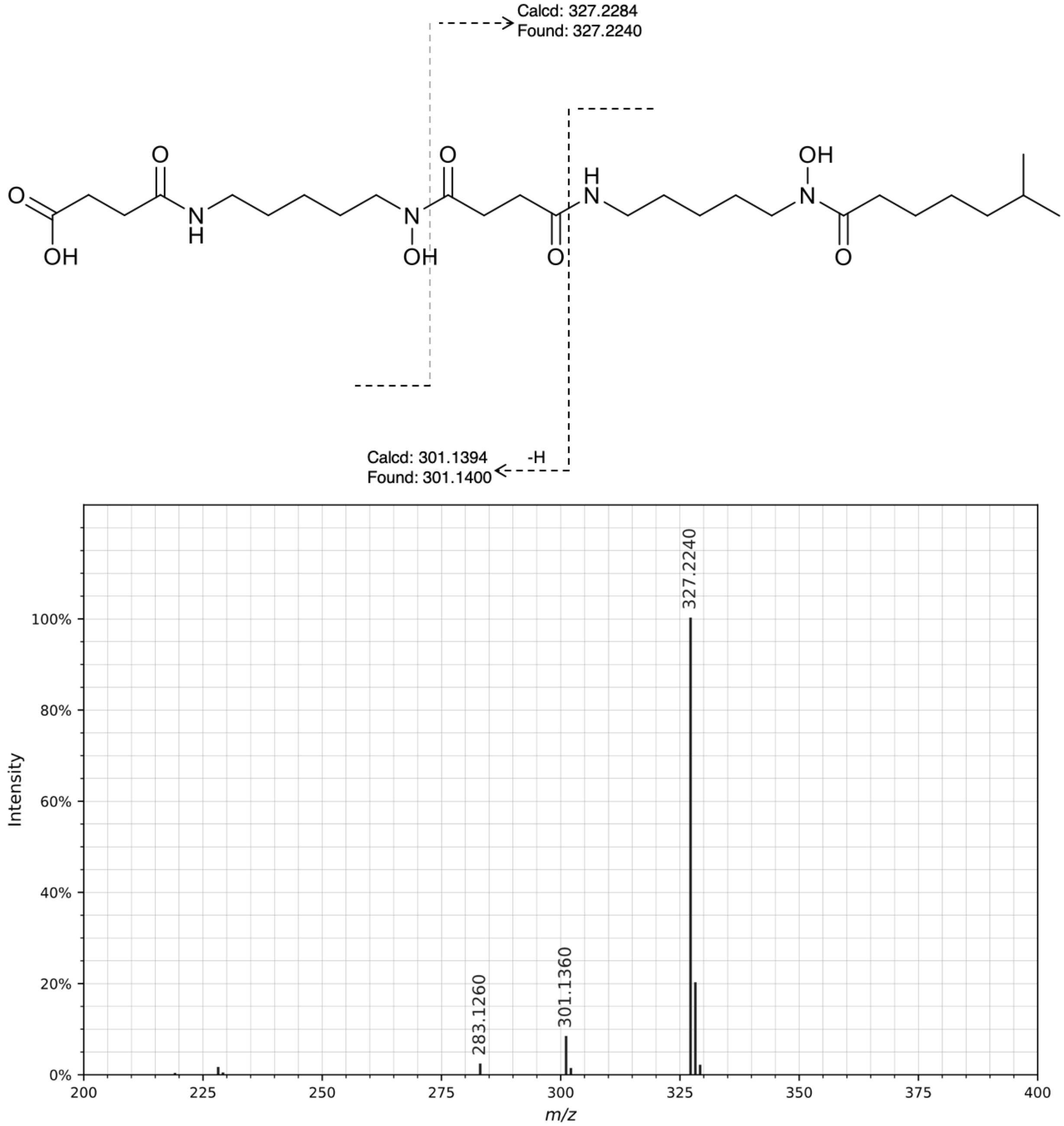


CID MS/MS spectrum of the $[M+H]^+$ ion of tenacibactin F. All fragment ions in the annotated structure match ions in the spectrum. Other fragment ions present in the spectrum are products of non-sequential fragmentation and are present in all DFOs.

Figure S49. Annotation of MS/MS for reported compounds: **tenacibactin G**

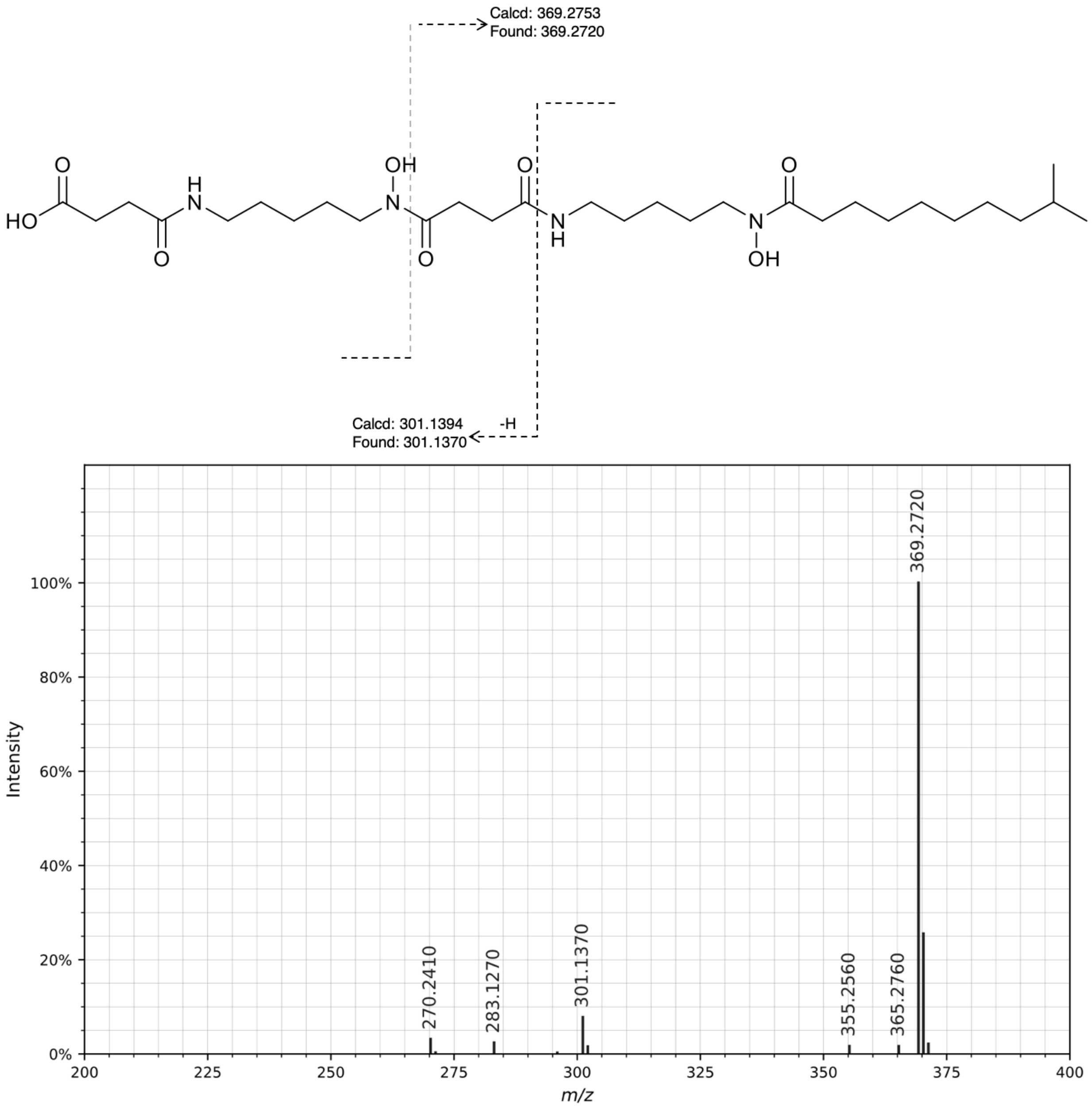


CID MS/MS spectrum of the $[M+H]^+$ ion of tenacibactin G. All fragment ions in the annotated structure match ions in the spectrum. Other fragment ions present in the spectrum are products of non-sequential fragmentation and are present in all DFOs.

Figure S50. Annotation of MS/MS for reported compounds: **tenacibactin H**

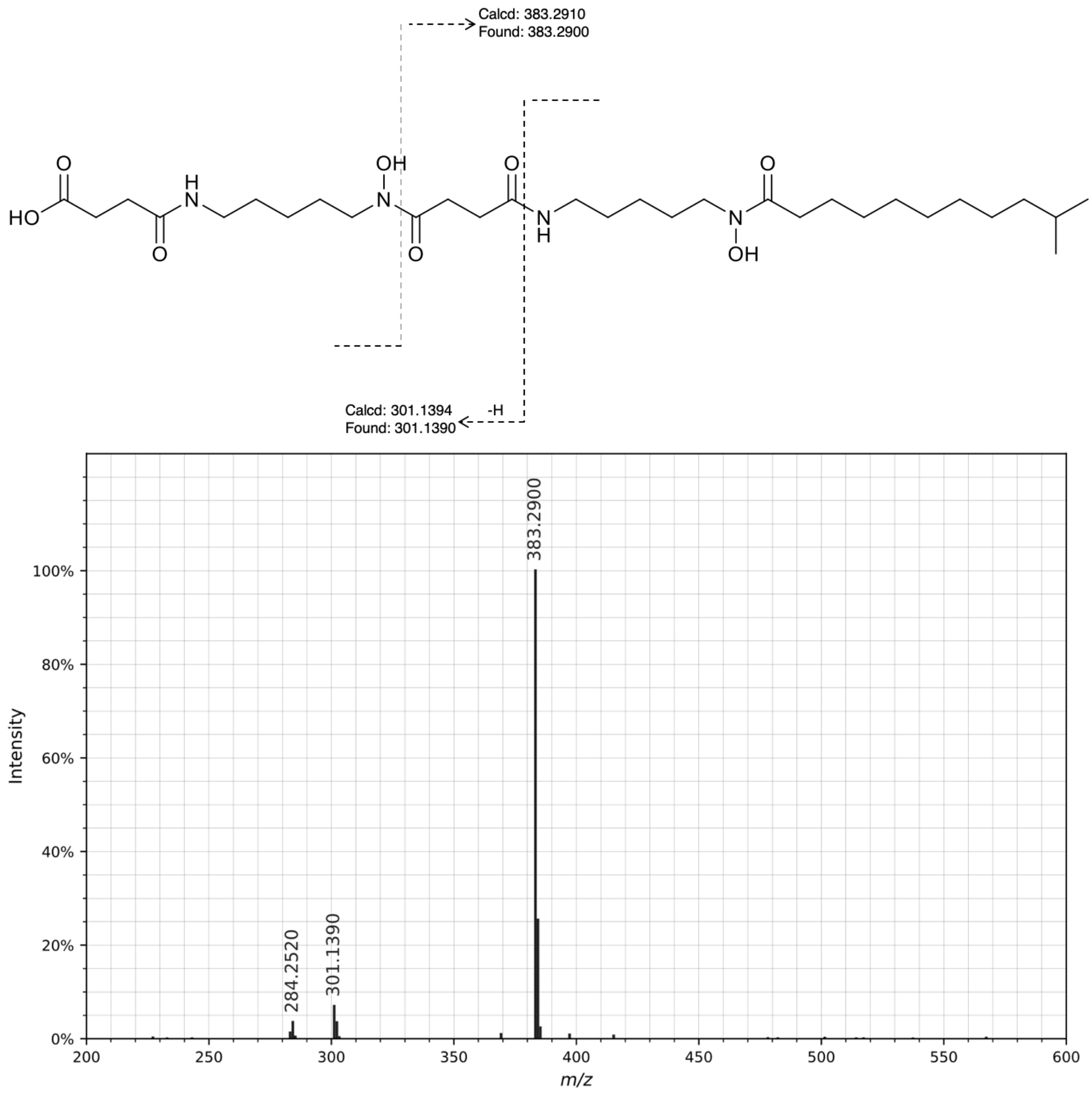


CID MS/MS spectrum of the $[M+H]^+$ ion of tenacibactin H. All fragment ions in the annotated structure match ions in the spectrum. Other fragment ions present in the spectrum are products of non-sequential fragmentation and are present in all DFOs.

Figure S51. Annotation of MS/MS for reported compounds: **tenacibactin I**

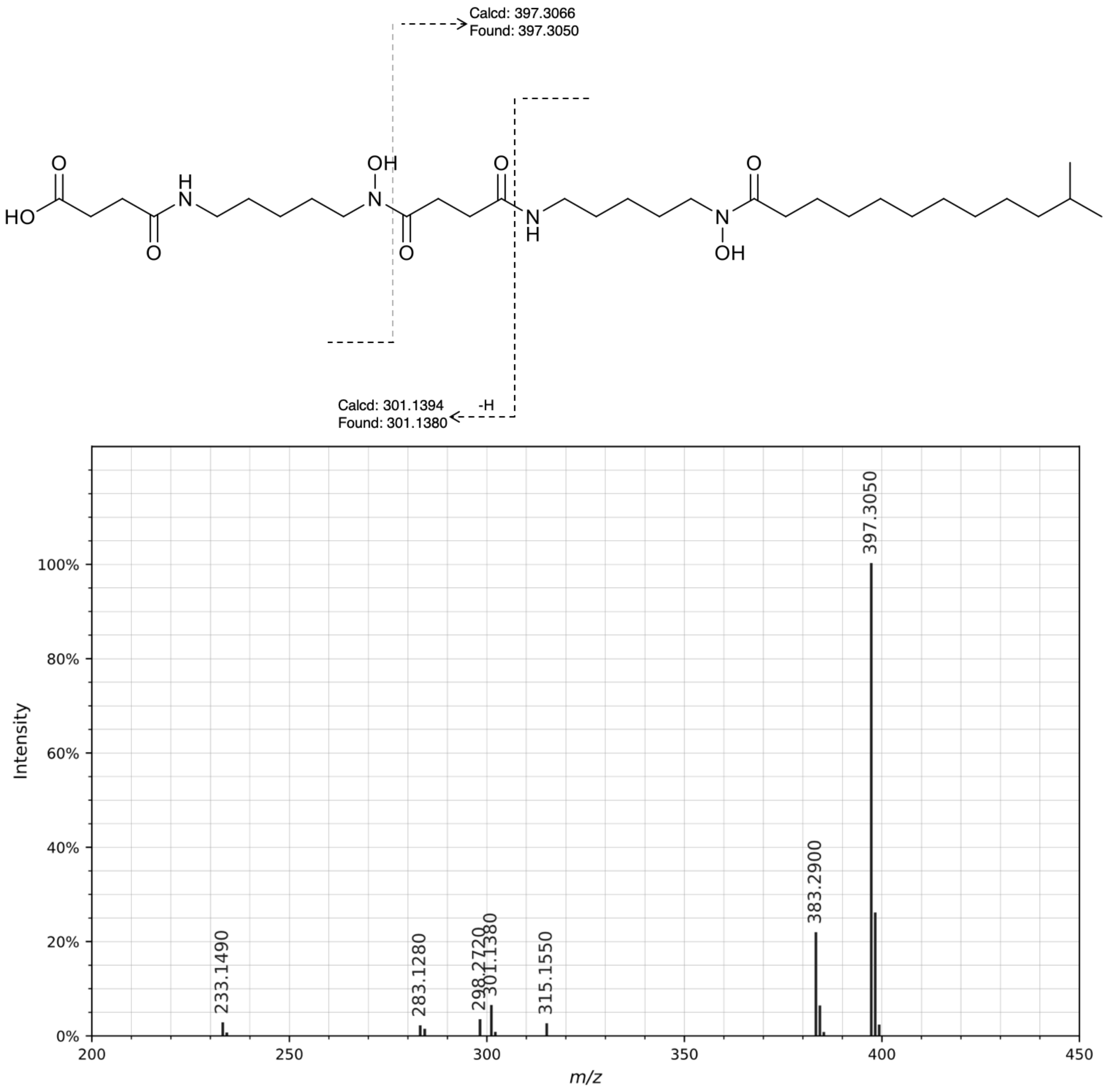


CID MS/MS spectrum of the [M+H]+ ion of tenacibactin I. All fragment ions in the annotated structure match ions in the spectrum. Other fragment ions present in the spectrum are products of non-sequential fragmentation and are present in all DFOs.

Figure S52. Annotation of MS/MS for reported compounds: **tenacibactin J**

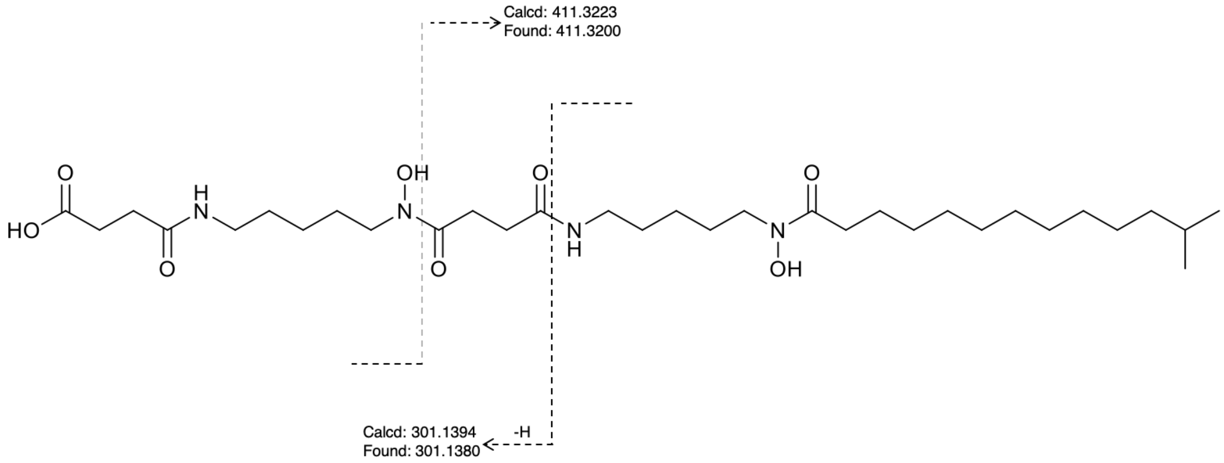


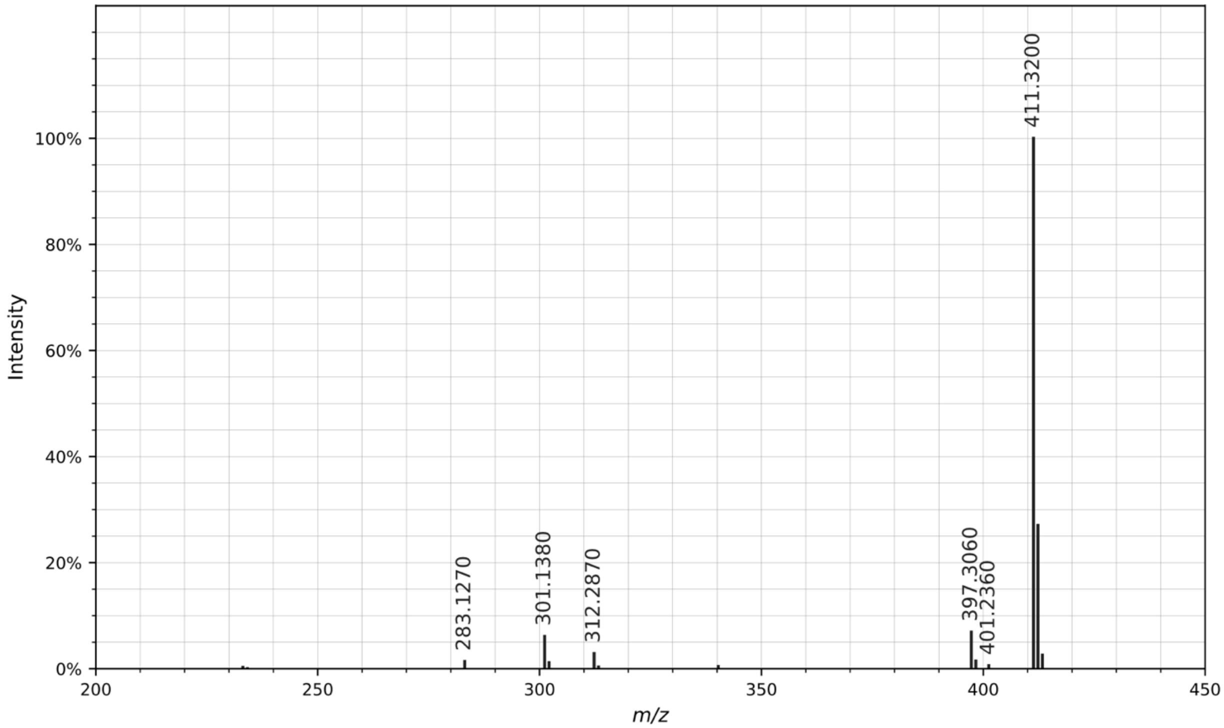


CID MS/MS spectrum of the $[M+H]^+$ ion of tenacibactin J. All fragment ions in the annotated structure match ions in the spectrum. Other fragment ions present in the spectrum are products of non-sequential fragmentation and are present in all DFOs.

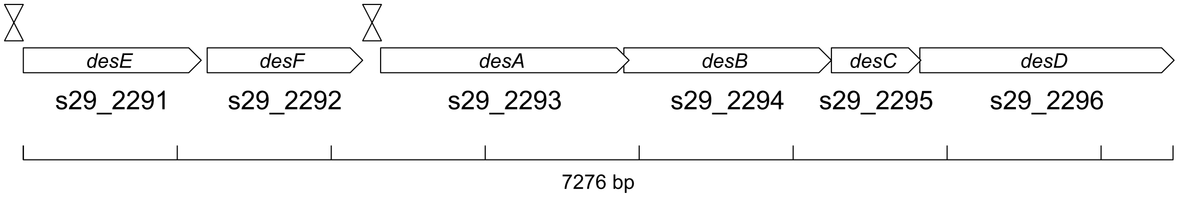


Figure S53. Region 7 - *des* gene cluster organization in *Streptomyces* sp. S29. The double arrowhead symbol indicates the location of iron boxes for DmdR.

>Botrytis cinerea – ITS region
TCCGTAGGGTGAACCTGCGGAAGGATCATTACAGAGTTCATGCCCGAAAGGGTAG
ACCTCCCACCCTTGTGTATTATTACTTTGTTGCTTTGGCGAGCTGCCTTCGGGCCT
TGTATGCTCGCCAGAGAATACCAAAACTCTTTTTATTAATGTCGTCTGAGTACTATA
TAATAGTTAAAACTTTCAACAACGGATCTCTTGGTTCTGGCATCGATGAAGAACGCA
GCGAAATGCGATAAGTAATGTGAATTGCAGAATTCAGTGAATCATCGAATCTTTGAA
CGCACATTGCGCCCCTTGGTATTCCGGGGGGCATGCCTGTTCGAGCGTCATTTCA
ACCCTCAAGCTTAGCTTGGTATTGAGTCTATGTCAGTAATGGCAGGCTCTAAAATC
AGTGGCGGCGCCGCTGGGTCCTGAACGTAGTAATATCTCTCGTTACAGGTTCTCG
GTGTGCTTCTGCCAAAACCCAAATTTTTCTATGGTTGACCTCGGATCAGGTAGGGA
TACCCGCTGAACTTAAGCATATCAATAAG

>Aspergillus niger – ITS region
GGTGAACCTGCGGAAGGATCATTACCGAGTGCGGGTCCTTTGGGCCCAACCTCCC
ATCCGTGTCTATTGTACCCTGTTGCTTCGGCGGGCCCGCCGCTTGTCGGCCGCCG
GGGGGGCGCCTCTGCCCCCCGGGCCCGTGCCCGCCGGAGACCCCAACACGAAC
ACTGTCTGAAAGCGTGCAGTCTGAGTTGATTGAATGCAATCAGTTAAAACTTTCAAC
AATGGATCTCTTGGTTCCGGCATCGATGAAGAACGCAGCGAAATGCGATAACTAAT
GTGAATTGCAGAATTCAGTGAATCATCGAGTCTTTGAACGCACATTGCGCCCCCTG
GTATTCCGGGGGGCATGCCTGTCCGAGCGTCATTGCTGCCCTCAAGCCCGGCTTG
TGTGTTGGGTCGCCGTCCCCCTCTCCGGGGGGACGGGCCCGAAAGGCAGCGGC
GGCACCGCGTCCGATCCTCGAGCGTATGGGGCTTTGTCACATGCTCTGTAGGATT
GGCCGGCGCCTGCCGACGTTTTCCAACCATTCTTTCCAGGTTGACCTCGGATCAG
GTAGGGATACCCGCTGAACTTAAGCATATCAAT

>S29 Omega-monooxygenase hit protein sequence 417 bp

MSCPALPEGFDATDPDLLQSRVPLPEFAQLRQTAPVWWCPQRRGVTGFDDEGYWAV
TRHADVKYVSTHPELFSSTVNTAIIRFNEHIPRDAIDAQRLIMLNMDPPEHTRVRQIVQR
GFTPRAIRGLETALRDRARKIAEEASAAAADGSFDFVTQVACELPLQAIAELIGVPQKDR
AKIFDWSNKMIAYDDPEYAITEEVGSNAAMELIGYAMNLSAQRKECPAQDIVTQLVAAE
GEGNLGSDEFGFFVLLLAVAGNETTRNAISHGMHAFLTHPEQWELYKRTRPSTAAEEI
VRWATPVVSFQRTATQDTELGGRKIKAGDRVGMFYSSANHDPEVFENPDVFDITRDP
NPHLGFGGGGPHFCLGKSLAVMEIDLIFNALADALPDLRLSGEPPRRLRAAWLNGIKEL
RVTSPGKD
